# Influence of predation mortality on past and future dynamics of Pacific Herring: implications for stock status and future biomass

**DOI:** 10.1101/2024.07.12.603178

**Authors:** Beau Doherty, Samuel D. N. Johnson, Ashleen J. Benson, Sean P. Cox, Jaclyn S. Cleary, Jim Lane

## Abstract

The recovery of marine mammals from historical over-exploitation in the 1970s represents one of the largest changes in trophic structure in the northeast Pacific Ocean over the last century, for which the impacts on key forage species such as Pacific Herring (*Clupea pallasii*) are poorly understood. This has prompted hypotheses that increasing marine mammal populations are the primary cause for productivity declines for some fish stocks and their lack of recovery to historical abundance levels. In this study, we evaluate such a hypothesis for Pacific Herring by quantifying historical predation rates by key predators including cetaceans (Pacific Humpbacks, Grey Whales), pinnipeds (Stellar Sea Lions, Harbour Seals), and piscivorous fish (Pacific Hake). Predation mortality is quantified via a novel approach that integrates a single-species catch-at-age model with estimates of predator consumption derived from bioenergetic models. We found that predator consumption, largely driven by Humpback Whales, explained increasing Pacific Herring natural mortality rates in recent years and could be used to forecast future mortality. Incorporating higher future natural mortality rates produced higher estimates of current stock status (1.09-1.2B0) based on lower estimates of equilibrium unfished biomass (17.5-20.3 kt). Conversely, models that assumed mortality was more like the historical average had lower stock status (0.63B0) and higher estimates of unfished biomass (32.4 kt). We demonstrate a practical approach for ecosystem modelling that can be used to develop operating model scenarios for management strategy evaluation, improving scientific defensibility by removing an element of analyst choice for future mortality scenarios. We discuss how simpler modifications to single-species model assumptions can be more pragmatic for providing fisheries management advice, while more complex multi-species or ecosystem models might provide more nuanced insights for exploring research questions related to multi-species ecosystems and fisheries interactions.

## 1. Introduction

Changes in the trophic structure of marine ecosystems can influence survival and recruitment of commercially exploited fishes and how their populations respond to fishing. The recovery of marine mammals such as Pacific Humpback Whales (*Megaptera novaeangliae*), seals (*Phoca spp*.), and sea lions (Family *Otariidae*) from historical over-exploitation represents one of the largest changes in trophic structure in the northeast Pacific Ocean over the last century, for which the impacts on key forage species are poorly understood. Specifically, it is unclear how prey consumption from increasing marine mammal populations may be contributing to increases in natural mortality observed for Pacific Herring (*Clupea pallasii*). Understanding how predators influence Pacific Herring mortality could provide key information for estimating stock status and future population trajectories under changing ecosystem conditions.

Pacific Herring play a key role in ecosystem function and food provisioning for commercial, social, and cultural uses throughout the northeast Pacific Ocean (Pikitch et al., 2012; Levin et al., 2016). Estimated fluctuations in productivity of Pacific Herring populations over the past hundred years were mainly attributed to exploitation and variable recruitment (Christensen et al., 2010; Martell et al., 2012); however, more recent declines in productivity are driven primarily by increasing trends in natural mortality (Benson et al., 2023; DFO, 2021; Johnson et al., 2024). For example, the exploitation rate for Pacific Herring off the West Coast of Vancouver Island (WCVI), British Columbia, Canada (Fig. 1) was relatively low over the 1980-2002 period (prior to fishery closures), ranging from 0% 26% (26% +/-4%), while estimated natural mortality rates increased from 0.48 /yr to 0.74 /yr over the same period (DFO, 2020). While interannual fluctuations in natural mortality are routinely estimated in assessments for this stock, the increasing trend presents a novel challenge for determining fishery reference points and understanding future population responses to fishing.

**Figure 1:**
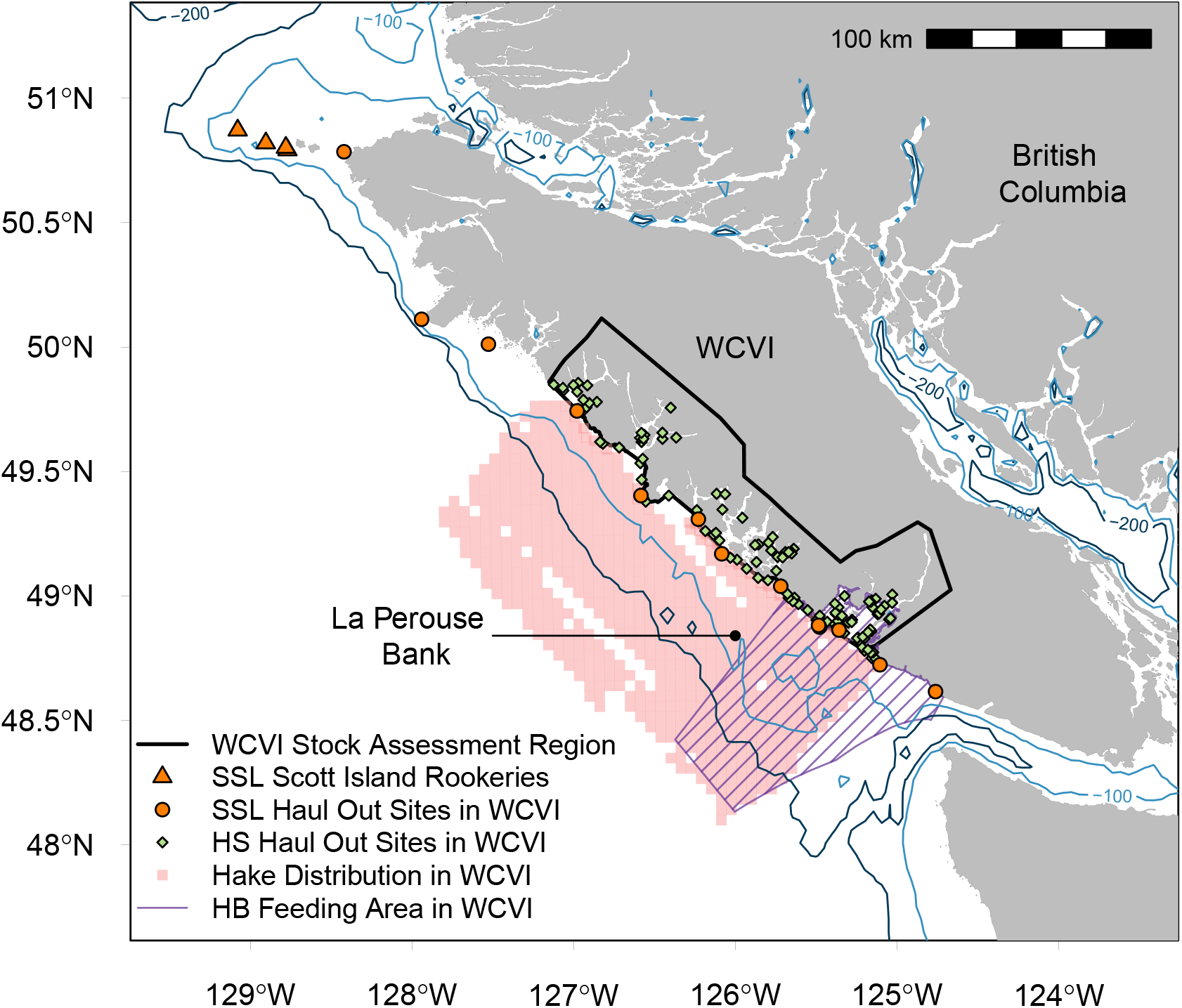
West Coast Vancouver Island (WCVI) Pacific Herring Stock Assessmet Region (SAR) and overlap with predator distributions for Humpback Whales (HB), Stellar Sea Lions (SSL), Harbour Seals (HS), and Pacific Hake (Hake). The Hake distribution reflects Hake presence for 2.5 nm grids from the acoustic survey from 1995-2019 for Pacific Fishery Management Areas (PFMAs) 123, 124, and 125. The HB feeding area is critical habitat identified in Nichol et al. 2010.

Despite commercial fishing closures since 2006 (DFO, 2024a,b), the recovery of WCVI Pacific Herring has been slow due to elevated natural mortality rates limiting production. The current regime of low biomass and high natural mortality could occur due to an Allee effect, whereby predation mortality rates increase for declining prey biomass (Gascoigne and Lipcius, 2004; Kronlund et al., 2018; Neuenhoff et al., 2019; Johnson et al., 2024). Indeed, positive correlations between WCVI Pacific Herring mortality rates and recovering marine mammal populations (Ford et al., 2009; Olesiuk, 2009; Chasco et al., 2017; Olesiuk, 2018) appear to support this hypotheses.

Most marine mammal populations in the northeast Pacific grew exponentially over the latter half of the 20th century in response to bans on commercial harvest and removal of predator control programs (Magera et al., 2013; Ford et al., 2009; Olesiuk, 1993). For example, recent estimates of Humpack Whales in the North Pacific range from 25,000-33,000 whales between 2007-2021 (Cheeseman et al., 2024), a remarkable recovery from the population lows of around 1,000 in the 1970s at the end of commercial whaling (Rice, 1978; Johnson and Wolman, 1984). The eastern stock for North Pacific Steller Sea Lions (*Eumetopias jubatus*) shares a similar story, which increased from around 10,000 to 35,000 individuals on breeding rookeries in British Columbia and Southeast Alaska between 1970 and 2013 following the end of commercial harvest and culling programs in the late 1960s (Olesiuk, 2018). Harbour Seals (*Phoca vitulina*) have also benefitted from increased legal protection for marine mammals and an end to seal harvests (Olesiuk, 1999), increasing 10-fold in British Columbia since the 1960s (DFO, 2022). These marine mammals feed on Pacific Herring, which has potential to produce sustained increases in mortality rates relative to historical levels with lower predator populations. Such an increasing trend in natural mortality, contradicts stationarity assumptions underlying fishery reference points from stock assessments, which are used to infer stock status relative to unfished equilibrium states. These shifting ecosystem dynamics present new challenges for fishery science and management to develop reliable estimates of stock status and forecasts for ecosystem conditions that differ from historical observations.

In this paper, we investigate the potential role of marine mammal and fish predators in driving historical changes in natural mortality rates and the implied effects on fishery reference points for WCVI Pacific herring. We construct a model of WCVI Herring that includes predation mortality from 5 predators (Plagányi et al., 2014, 2022), including Humpback Whales, Steller Sea Lions, and Harbour Seals, mentioned above, as well as Grey whales (*Eschrichtius robustus*), which feed on herring spawn deposited on benthic substrate, and Pacific Hake (*Merluccius productus*), which have more of a size-dependent predation. For each predator, herring consumption is estimated via a bioenergetic approach (Chasco et al., 2017), accounting for average predator size, estimates of predator abundance, and herring energy content. Predators are then modeled like a fishery in a Bayesian statistical catch-at-age model of the herring population, where consumption is treated as ‘catch’ and where each predator ‘fleet’ has a herring size preference. The predation mortality model is compared to a more typical assessment model where herring mortality is modeled as a random walk (Martell et al., 2012), as in regular herring stock assessments (Benson et al., 2023). Models are then projected into the future to estimate unfished equilibrium conditions that incorporate uncertainty in future predator populations. Our results show how explicitly accounting for predation can i) explain historical natural mortality similar to a random walk, ii) improve credibility of future natural mortality projection scenarios used for estimating reference points and developing harvest strategies, iii) improve understanding of ecosystem requirements when developing management advice frameworks, and iv) compare estimates of unfished equilibrium states and current stock status under alternative natural mortality assumptions, with and without separate components for predation mortality.

## 2. Methods

### 2.1. Catch-at-age model with predation mortality

We develop a statistical catch-at-age (SCAH) model for Pacific Herring that expands on previous WCVI herring stock assessment models to explicitly incorporate mortality from major predators (Benson et al., 2023; Johnson et al., 2024; DFO, 2024a). SCAH is an age structured population dynamics model that includes four fishing fleets (reduction, seine-roe, gillnet, spawn-on-kelp) and seven predator groups targeting Pacific Herring (Summer Humpback Whales, Winter Humpback Whales, Steller Sea Lions, Harbour Seals, Small Hake, Large Hake) and their eggs (Grey Whales). Predators are treated the same way as fishing fleets and require annual consumption estimates (i.e., predator ‘catch’) for each predator as model inputs, which are derived using bioenergetic models and annual abundance estimates of each predator.

A full description of the SCAH model can be found in supplementary material A, including notation (Table A.1), process model equations (Table A.2), and observation models and objective functions (Table A.3). Data inputs (Fig. A.1) for fitting the SCAH model include: (i) fleet-specific landed catch and age-composition data (1951 - 2022) from the commercial reduction fishery, seine-roe, and gillnet fisheries; (ii) closed-pond spawn on kelp (SOK) landings, (iii) fishery independent spawner indices that blends surface (1951 - 1987) and dive (1988 - 2022) survey designs, (iv) annual estimates of Herring biomass consumed by a set P of seven predator classes, namely two Pacific Hake size classes (hakeLt50 and hakeGt50, *Merluccius productus*), two Humpback Whale classes broken up by feeding behaviour in the winter and summer (hbW and hbS, *Megaptera novaeangliae*), Steller Sea Lions (SSL, *Eumetopias jubatus*), Harbour Seals (HS, *Phoca vitulina*), and Grey Whales (GW, *Eschrichtius robustus*), and (v) model estimates of total numbers for all predators (described in supplementary material B) except Pacific Hake, for which we use estimates of biomass at-age from the stock assessment to derive predator size class biomass (Johnson et al., 2021).

The main novelties in our SCAH model for WCVI Pacific Herring are the inclusion of predator consumption (whole herring predation) and removals of fertilised herring eggs from both SOK fisheries and predation (Grey Whales). Since predators are treated similar to commercial fisheries, with input catch data and assumed selectivity functions, most of the population dynamics for SCAH are governed by standard age-structured equations (Appendix A). In the following sections we outline the modelling approaches used for including the removals of fertilised herring spawn by SOK fisheries and Grey Whales, estimating predation mortality on WCVI herring, and the estimation of long-term equilibrium unfished biomass via simulation under uncertainty about future modeled predation mortality.

#### 2.1.1. Fertilised herring egg removals

This subsection described closed pond SOK fisheries and grey whale predation, and, briefly, how egg removals are incorporated into the SCAH model. A more detailed explanation along with the process model equations are provided in supplementary material A (Table A.1, eqns K.1-K.6 in Table A.2).

Closed pond fisheries involve the impoundment (ponding) of fish from the spawning stock. Pacific Herring are captured from spawning aggregations via purse seines, and then towed to and transferred into a closed net-pen (pond). Ponds have material suspended inside for herring to deposit eggs onto, primarily kelp blades (*Macrocystus integrifolia*) and also feather boa (*Egregia menziesii*), depending on the local practice (Schweigert et al., 2018). Additionally, hemlock (*Tsuga* spp.) and red cedar (*Thuja* spp.) boughs can be set in or near ponds as Indigenous food harvest practices. When fish are ponded they are effectively removed from the spawning stock biomass for the remainder of the year. A post-ponding mortality rate *M*_*SOK*_ = 0.35 is used to account for fish that do not survive impoundent (supplementary material A), while surviving fish are reunited with the main population upon release.

Grey Whale predation on fertilised eggs is treated slightly different to eggs removed through the spawn-on-kelp fisheries. Early in the spawn season, between late February and early March, Grey Whales feed on fertilised Herring spawn in the WCVI (Darling et al., 1998). Grey Whales feed on eggs by diving in areas with sufficient depth for their body size, and scraping their mouths along the bottom in spawn areas. To account for Grey Whale predation, the SCAH model temporarily reduces spawning biomass prior to recruitment by the equivalent biomass needed for generating the eggs consumed by Grey Whales, since their consumed eggs will not contribute to new recruitment. In contrast to closed-pond spawn fisheries, there is no additional mortality associated with the egg predation when adding spawners back into the main population because the fish are not physically impounded.

#### 2.1.2. Natural mortality models

SCAH models are fit under two natural mortality hypotheses

##### 1. noPredM

a time-varying mortality *M*_*t*_ that is the same for all age-classes *a* ∈ {1, …, 10} and includes all sources of natural mortality modelled as a random walk

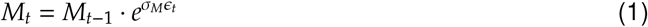

where *M*_*t*_ is the instantaneous natural mortality rate at time *t*, and *ϵ*_*t*_ ∼ N(0, 1) with *σ*_*M*_ = 0.1 as the random walk deviation.

##### 2. predM

A time-varying natural mortality *M*_*a,t*_ for age *a* at time *t*, divided into separate components

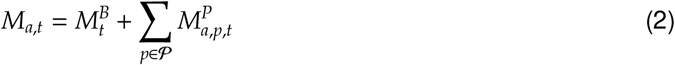

where 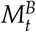 is a time-varying basal mortality rate (e.g., disease, senescence, environmental effects, starvation, unmodelled predators, unreported catch) modelled via random walk (as in (1) for *noPredM*), and 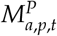 is time-varying predation mortality-at-age, summed over the set of modeled predators 𝓅.

### 2.2. Modelling historical predation mortality rates

This section details the components of the predator mortality model for the *predM* hypothesis. First, we describe the bioenergetic approach and selectivity curves for estimating annual predator consumption and age composition. Predator consumption is then used to estimate historical per-capita predation mortality rates via the Baranov catch equation, which is in turn used to estimate the relationship between predator density, prey density, and consumption under an assumed functional response or a foraging arena theory model (Holling, 1959; Ahrens et al., 2012).

#### 2.2.1. Predator consumption and bioenergetic models

Under the *predM* hypothesis, annual consumption of Pacific Herring by each predator is estimated via a model of each predator’s bioenergetic requirements (Chasco et al., 2017). We explain the models used for hbW, hbS, HS, and SSL first, which all share the same modeling approach. Second, we described the approach for Grey Whale egg consumption, where several assumptions are required due to data limitations on population sizes and feeding habits. Third, we explain the Hake model, for which energetic requirements are calculated for two age-class groups. Table 1 summarizes the data, assumptions, and literature sources used for estimating bioenergetic requirements for each predator.

**Table 1:** Parameter values and sources for bioenergetic models of Herring consumption for Harbour Seals (HS), Steller Sea Lions (SSL), and Humpbacks (HB, winter and summer feeding groups)

| Predator | Parameter | Value | Source |
| --- | --- | --- | --- |
| HS | $\alpha$ | 103 | Howard et al. 2013, Chasco et al. 2017 |
| | $\beta$ | 0.75 | Kleiber 1961 |
| | $W$ | 44.7 kg | Olesiuk 1993 |
| | $d$ | 365 days | year-round foraging, Olesiuk 1993 |
| | $\rho$ | 0.32 | Olesiuk 1993 |
| | $\lambda$ | 0.825 | Howard et al.2013; Chasco et al. 2017 |
| SSL | $\alpha$ | 216 | Weighted average by age-composition (Winship et al. 2002, Chasco et al. 2017) |
| | $\beta$ | 0.75 | Kleiber 1961 |
| | $W$ | 220.6 kg | Calculated from weight-at-age, proportions-at-age, and sex composition data from Chasco et al. 2017 |
| | $d$ | 365 days | year-round foraging, Tollit et al. 2015 |
| | $\rho$ | 0.086 | Estimates from variable percent energy contribution models (Tollit et al. 2015) |
| | $\lambda$ | 0.875 | Winship et al. 2002, Chasco et al. 2017 |
| hbS, hbW | $\alpha$ | 209 | Moran et al. 2018 |
| | $\beta$ | 0.75 | Kleiber 1961 |
| | $W$ | 25.9 t | Weight-length conversion for 12.3 m based on length distribution from BC Whaling records (Nichol and Heise 1992, Moran et al. 2018) |
| | $d$ | Summer: 76 days | Summer: Aug 10 - Oct 15 (McMillan 2014) |
|  |  | Winter: 151 days | Winter: Oct 16 - Mar 15 (Moran et al. 2018) |

Table 1: Parameter values and sources for bioenergetic models of Herring consumption for Harbour Seals (HS), Steller Sea Lions (SSL), and Humpbacks (HB, winter and summer feeding groups) (continued)
| Predator | Parameter | Value | Source |
| --- | --- | --- | --- |
| | $\rho$ | Summer: 0.62 | Mean proportions from Sept 15 - Oct 15 (Moran et al. 2018) |
|  |  | Winter: 0.65 | Mean proportions from Oct 16 - Mar 15 (Moran et al. 2018) |
| | $\lambda$ | 0.9 | Laidre et al. 2007 |
| hakeLt50,hakeGt50 | $d$ | 183 | Jun 1 - Nov 30 (Tanasichuk et al. 1991) |
| GW | $d$ | 60 | Feb 14 - Apr 15 (Darling et al. 1998) |
| | $\lambda$ | 0.8 | Highsmith and Coyle 1992 |

##### 2.2.1.1 Humpback Whales, Stellar Sea Lions, and Harbour Seals

For predator classes *p* ∈ {hbW, hbS, HS, SSL}, the the total annual energy *E*_*p*_ (kcal) that each individual predator derives from Herring is calculated as

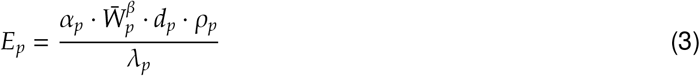

where *α*_*p*_ and *β* are parameters describing the relationship between daily metabolic rate (kcal/day) and body weight in kg (Kleiber, 1961), 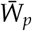 is the average predator weight in kg, *d*_*p*_ is the number of annual foraging days for Herring in WCVI, *ρ*_*p*_ is the fraction of energy the predator derives from Herring on herring foraging days, and *λ*_*p*_ is the average energy digestive efficiency. Harbour Seals and Steller Sea Lions feed on Pacific Herring year round, whereas Humpback Whales target herring during specific times of the year based on availability and each species’ seasonal migration patterns (Table 1).

Annual consumption for year *t* for each predator is calculated from annual energy requirements as

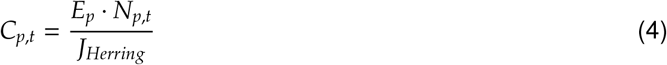

where *C*_*p,t*_ is the predator consumption, *N*_*p,t*_ is the number of predators (estimated from models described in supplementary material B), and *J*_*Herring*_ = 1.82 kcal/g is the average energy content for Pacific Herring (Moran et al., 2018).

##### 2.2.1.2. Grey Whale egg consumption

Grey Whales are modelled similarly to the other marine mammals above, but feed on herring eggs with a different energy content. An individual Grey Whale has a daily energy requirement estimate of 760,000 kCal/day (Highsmith and Coyle, 1992), which is adjusted for a digestive efficiency of *λ*_*GW*_ = 0.8 to give the daily energy requirement as 950,000 kCal/day during the spawning season. The same bioenergetic requirements from equations (3) - (4) are then applied, with a modification to replace diet proportion *ρ*_*p*_ with a scalar *r*_*GW*_, i.e.,

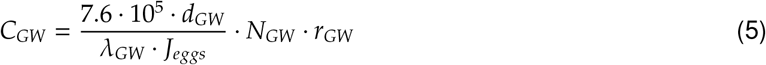

where *d*_*GW*_ = 60 days during the spawning season, *J*_*eggs*_ = 4.47 kcal/g is the average energy content for Pacific Herring eggs (Bishop and Green, 2001), and *N*_*GW*_ are abundance estimates for the Pacific Coast Feeding Group (PCFG) of Grey Whales *N*_*GW*_ (supplementary material B) derived from a recent recovery potential assessment (Gavrilchuk and Doniol-Valcroze, 2021). The scalar *r*_*GW*_ = 0.05 accounts for the proportion of PCFG Grey Whales that (a) are not present in WCVI during the Herring spawn season, and (b) do not feed exclusively on herring eggs, both of which are very uncertain based on limited quantitative data; however, recent tagging studies suggest that the proportion of Grey Whales feeding in the WCVI area is low (Lagerquist et al., 2019). The assumption of *r*_*GW*_ = 0.05 represents 5% of PCFG whales feeding exclusively on Herring eggs for the full 60 day spawning season, which is roughly in the middle of the range of two main qualitative observations. The first is from Darling et al. (1998), observing dozens to hundreds of whales utilizing the site (Clayquot Sound) usually for 2-3 weeks between mid February and early April; the second being from Ford et al. (2013) that fewer than 100 PCFG whales are likely to occupy BC waters in any given year. At current PCFG abundance, 5% represents 12.5 whales feeding exclusively for 60 days.

##### 2.2.1.3. Pacific Hake

The Pacific Hake bioenergetic model takes advantage of the age-structured population dynamics parameters available from the stock assessment model (Johnson et al., 2021). Combined sex biomass-at-age estimates from the Hake stock assessment are converted into piscivorous Hake biomass-at-age for males and females in the offshore areas (PFMAs 123, 124, and 125, Fig. 1) adjacent to WCVI Pacific Herring spawning areas each year, as described in Supplementary B. Consumption of WCVI herring by each Hake age-/sex-class is

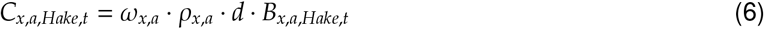

where *ω*_*x,a*_ is the amount of food consumed as a percentage of Hake body-weight per day (Francis, 1983), *ρ*_*x,a*_ is the proportion of an age-*a* and sex-*x* Hake’s diet that is Herring, *d* is the number of days per year where Hake and Herring overlap off the coast of WCVI, and *B*_*x,a,Hake,t*_ is the piscivorous biomass of sex-*x* and age-*a* Hake in the WCVI region in year *t*. The sex- and age-specific diet proportion *ρ*_*x,a*_ is derived by integrating the observed diet proportion *ρ*_*l*_, where *l* is the length classes *l >* 50 cm or *l* ≤ 50 cm (fork lengths), over the estimated uncertainty in Hake von Bertalanffy growth models (supplementary B.1.4). The number of days feeding on WCVI Herring is fixed at *d* = 183 to account for feeding between June 1 - November 30 (Tanasichuk et al., 1991).

The *ω*_*x,a*_ for each age-class and sex is calculated as:

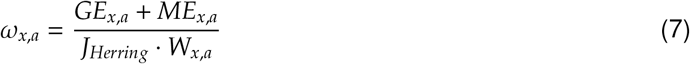

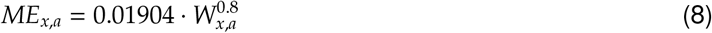

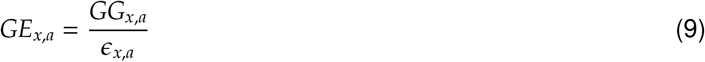

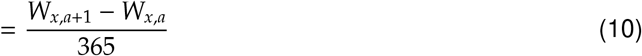

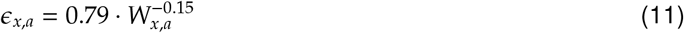

where *W*_*x,a*_ is mean weight-at-age, *J*_*Herring*_ = 1.82 kcal/g is the energy content of Herring, *ME*_*x,a*_ is energy required for maintenance in kcal/day, *GE*_*x,a*_ is energy required for growth and reproduction in kcal/day, *GG*_*x,a*_ is the gross growth per day in g/day, and *ϵ*_*x,a*_ is grow growth efficiency in g/kcal (Francis, 1983; Jones and Johnston, 1977). The weight at-ages *W*_*x,a*_ for males and females are drived from allometric length/weight and von Bertalanffy growth models (supplementary material B).

Finally, the consumption of WCVI Pacific Herring by each Hake predator size class *p* ∈ {hakeLt50, hakeGt50} is estimated by

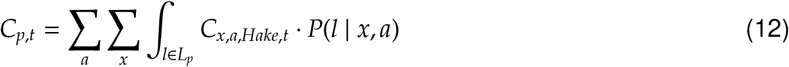

where *L*_*p*_ is the length interval for each Hake predator size class *p*, and *P*(*l* | *x, a*) is the probability a Hake is length-*l* given that it is sex-*x* and age-*a*.

#### 2.2.2. Predator Selectivity

Selectivity curves *s*_*p*_ for predators are based on a literature review of each predator’s size preferences and diet composition.

Steller Sea Lion (SSL) selectivity is modeled as a logistic function of Herring length *l*

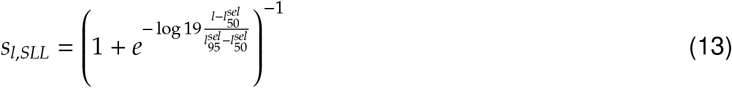

where 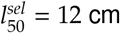 (ages 1-2) and 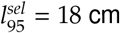 (age 3) are the lengths at 50% and 95% selectivity, respectively, based on Pacific Herring sizes in SSL diets in Southeast Alaska (Tollit et al., 2015).

For Harbour Seals, we assume a dome-shaped length-based selectivity curve

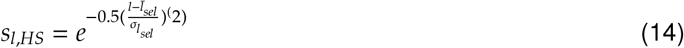

where 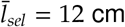 is the mean selectivity with standard deviation 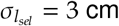, based on Harbour seal size preferences for smaller herring ranging from 10-14 cm (Tollit et al., 1997). SSL and HS selectivity-at-length *s*_*l,p*_ are converted to selectivity-at-age *s*_*a,p*_ via a Herring age-length key, estimated from herring age and length data collected from commercial sets and scientific test fishery sets in the WCVI area (DFO, 2024a).

Both Humpback Whales and Pacific Hake are split into two predation groups with different size selectivity. Humpback Whale size preference is seasonal, with a diet mostly made up of young-of-year and juveniles with sizes less than 15 cm during summer months (McMillan, 2014), switching to larger fish (age 2+) once mature Pacific Herring begin migrating inshore for over-wintering in October (Megrey et al., 2007). Therefore, two Humpback Whale feeding groups are defined with different selectivities reflecting the availability of herring during the summer feeding period from August 1 - October 15, where Humpback Whales are feeding only on age-1 and age-2 fish (McMillan, 2014), and the winter feeding period from October 16 - March 15 (Moran et al., 2018), when Humpback Whales are feeding on a broader range of sizes. Selectivity is fixed at 100% for ages 1-2 and 0% for ages 3+ for the summer humpback feeding group (McMillan, 2014), while selectivity for the winter humpback feeding group is fixed at 100% for ages 2-10 and 0% for age-1.

Pacific Hake target offshore herring from July to November prior to spawning migrations, and there are significant differences in the size frequency of herring observed for different Hake predator size classes (Fig B.9). Therefore, as mentioned above, Hake are split into two size classes of fish less than 50 cm in length (hakeLt50) and above 50 cm in length (hakeGt50). Additionally, the maximum size (length) of prey is limited by the size of the predator, with stomach volume and jaw openings being limiting factors. Therefore, a double-asymptotic dome-shaped size-selectivity is assumed for each Hake predator class, defined as

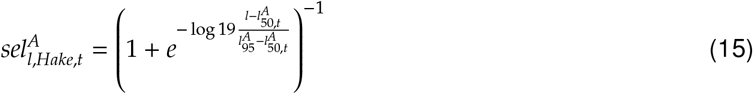

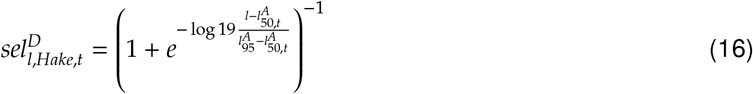

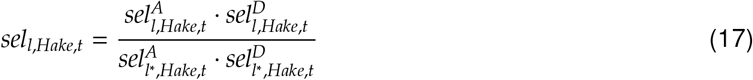

where 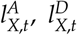 is the selectivity-at-length for ascending and descending limbs, respectively, in year *t*. The final size selectivity *sel*_*l,Hake,t*_ is then the product of the ascending and descending limbs, normalized so that the modal length bin *l*^∗^ is 100% selected. Time variation in Pacific Hake size selectivity is from scaling the length-at-50% and -95% selectivity for both ascending and descending limbs by the mean length of Pacific Hake within each size class for year *t*, which is derived from the sex-specific piscivorous Hake biomass-at-age and growth model (Figure B.8).

Size-selectivity model parameters are estimated by fitting to stomach contents data (supplementary material B.1.4, Fig. B.9) from survey observations of Pacific Hake feeding on Pacific Herring on La Perouse bank (Tanasichuk et al., 1991), where WCVI Herring feed during the summer and early fall. Given ad-hoc timing of the La Perouse Bank survey and relatively low sample sizes of stomach contents data across years, the data likelihood for each predator size class aggregates samples over all years as a mixed model, with annual samples weighted by sample size. Additionally, to account for variable year class strength, the expected values for yearly prey length compositions are scaled by WCVI stock-assessment numbers-at-length, using the same Pacific Herring age-length key described at the top of this section (DFO, 2020).

There is no size-selectivity for Grey Whales. Grey Whale consumption of fertilised herring eggs is converted to an equivalent mature spawning biomass estimate following the same conversion methods used in annual DFO spawn surveys (Grinnell et al., 2023).

#### 2.2.3. Estimating historical predation mortality rates

The consumption of individual herring by the main predators (i.e., Hake, HS, SSL, hbW, and hbW) is treated as a continuous process with an instantaneous predation mortality rate that is applied continuously over the entire year. While temporal overlap between some predators and WCVI herring is less than an entire year, the effect of lower average predation mortality rates over a longer time period on estimates of spawning biomass and fishery catch will be minor, given that spawning and commercial removals occur towards the end of the time step. In contrast to most predators that occur year-round or over several months, commercial fisheries and Grey Whale predation tend to occur over one or two month periods during the spawning season. Therefore, fisheries removals (described in supplementary material A) and Grey Whale predation are modeled as discrete mortality processes where individual herring or eggs are removed as a single pulse event occurring at a fractional time of year *δ*_*g*_ during the spawning season, where *δ*_*reduction*_ = 0.50 for the reduction fishery, *δ*_*seineroe*_ = 0.65 for the seine-roe fishery, *δ*_*gillnet*_ = 0.67 for the gillnet fishery, *δ*_*SOK*_ = 0.68 for the SOK fishery, and *δ*_*GW*_ = 0.69 for Grey Whale predation.

The Baranov equation is modified to account for continuous mortality applied between discrete pulse removals of commercial fisheries and Grey Whale predation. This modification is necessary because, especially in early years, the biomass can change a lot at the moment a commercial fishery takes its entire catch, so the order that mortality is applied is important. Fisheries are indexed by *g* = 1, …, 4 and remove fish at the fractional time steps *δ*_*g*_, *δ*_*g*_ ≤ *δ*_*g*_′ when *g* ≤ *g*^′^ (i.e., fractional time steps are ordered from 1 to 4). We also include *g* indices to represent the start (*g* = 0) and end (*g* = 5) of the year with fractional time steps *δ*_0_ = 0 and *δ*_5_ = 1 to model predator consumption before the first fishery (*g* = 0 to *g* = 1) and after the last fishery (*g* = 4 to *g* = 5). Expected consumption of herring by predator *p* for a fractional time step ending at *δ*_*g*_ (i.e., between *δ*_*g*−1_ and *δ*_*g*_) is then estimated as

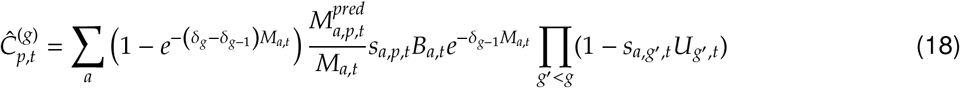

where 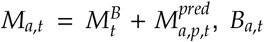 is the start of year herring biomass at age *a*, and *U*_*g,t*_ = *C*_*g,t*_*/B*_*g,t*_ is the harvest rate for gear *g* at time *t* (see Table A.2 for discrete fishery equations). For the closed-pond spawn fishery removals, harvest rates are based on vulnerable biomass equivalents after transformation by the Ψ_*g,t*_ factor (described in supplementary material A, K.1 in Table A.2) used to convert between landed eggs and the equivalent biomass. Then, by stepping through fishing gears, total expected predator consumption is estimated by adding up consumption before and after each fishing gear *g* as

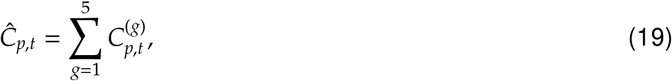

which is used as the mean value in the catch data likelihood (Co.1-Co.2, Table A.3).

Consumption estimates from predator bioenergetic models are treated as data inputs to SCAH. When fitting SCAH, an expected value of consumption is derived from input predator abundance, vulnerable herring biomass, and predator selectivity-at-age via the Baranov catch equation, modified to reflect the mix of discrete and continuous removals as described above. Total continuous mortality-at-age for the Baranov catch equation (described below) is the sum of basal mortality 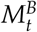 and predation mortality 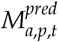 for predators *p* ∈ {hakeLt50, hakeGt50, hbW, hbS, HS, SSL} whose consumption is modelled as a continuous process, i.e.

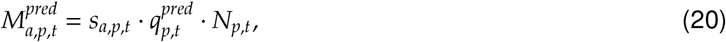

where *s*_*a,p,t*_ is herring selectivity-at-age by predator *p* at time *t* (only time-varying for Hake) derived from predator size preference discussed above, 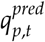 is the per-capita predation rate, and *N*_*p,t*_ is the abundance (numbers or biomass) of predator *p*. The per-capita predation rate 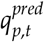 converts predator ‘effort’ (i.e., abundance) to instantaneous herring mortality, and is modeled as a simple random walk in log space

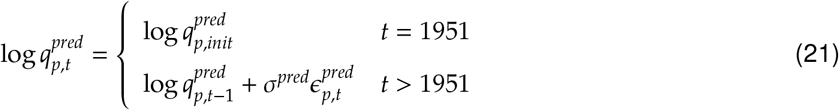

with interannual jumps *ϵ*_*p,t*_ ∼ *N*(0, 1). The initial value 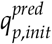 and jumps *ϵ*_*p,t*_ are model parameters, estimated during model fitting by comparing expected consumption from the Baranov catch equation to input consumption ‘data’ using a log-normal likelihood (Table A.3) with coefficients of variation (CVs) of 27% for hakeLt50, 13.5% for hakeGt50, 3% for HS, 12% for SSL, 15% fo hbW, and 24% for hbS (labelled as 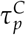 in Co.1 of Table A.3). The CVs were determined via the estimated log-scaled standard deviations from residuals between MLE model estimates of consumption and the expected predation consumption from bioenergetic models, which were then increased by 50% to allow for greater uncertainty when estimating posteriors via Hamiltonian Monte Carlo integration.

### 2.3. Modelling future predation mortality rates

To project predation mortality into the future, a model of the relationship between prey density, predator density, and predation mortality is required. Here, we use two different approaches to post-fitting predation mortality rate models to SCAH model estimates, with the approach depending on the predator.

The first approach, for Pacific Hake and Harbour Seals, is to model predation rates as a Type II or III functional response (Holling, 1959). Functional responses relate per-capita predation rates 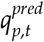 to prey biomass, for which we used vulnerable herring biomass *B*_*p,t*_ = Σ _*a*_ *s*_*a,p,t*_*B*_*a,t*_. Maximum posterior density estimates of 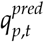 and *B*_*p,t*_ are used for post-fitting the functional response model

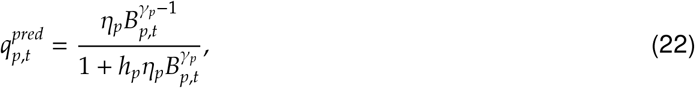

where *η*_*p*_ is the encounter rate between Pacific Herring and predator *p, h*_*p*_ is the handling time, and *γ*_*p*_ determines the shape of the functional response. A predator with *γ*_*p*_ = 1 has a Type II functional response, where predation rate continually increases as prey density declines, whereas *γ*_*p*_ *>* 1 produces a Type III functional response, where predation rate is maximised at a positive prey density, and then declines as prey density declines. A Type III functional response is commonly associated with a predator switching to prey that are easier to catch or more prevalent. For WCVI Herring, we assumed a Type III functional response (i.e., *γ*_*p*_ *>* 2) for both Pacific Hake predator size classes, and a Type II functional response for Harbour Seals. The second approach uses foraging arena theory to estimate absolute predation rates for Humpback Whales and Steller Sea Lions (Ahrens et al., 2012). Similar to a functional response, foraging arena theory relates predation rates, prey density, and predator density. The two main differences are (i) the inclusion of prey exchange between vulnerable states while feeding (i.e., in the foraging arena) and invulnerable states between feeding times, and (ii) predation rates are a function of predator abundance instead of prey biomass. The exchange between vulnerable and invulnerable states imposes an upper bound on the total predation mortality rate *M*_*p,t*_ or, equivalenty, imposes a dome-shaped relationship between per-capita rates 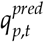 and *N*_*p,t*_, which decays towards zero as predator abundance increases. The limit on total predation mortality is required because preliminary simulations found that a Type II or III functional response always leads to extirpation of the WCVI herring stock in simulated projections. Extirpation occured because higher predator densities produced total mortality levels above a sustainable level for WCVI Herring, even while per-capita predation rates were limited over the simulated herring biomass levels.

Foraging arena theory relates absolute predation mortality rates to predator density as

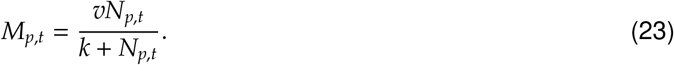

where the parameter *k* is the Michaelis constant, or the predator density at which predation mortality is half of the exchange rate *v* (Ahrens et al., 2012). This relationship does not hold exactly for the logistic modification in equation (24), but provides some valuable context.

We modified equation (23) to fit directly to predation mortality rates, by adding a logistic model factor to bound the absolute rate as predator density increases and match the sigmoidal shape of SCAH model estimates of predation mortality

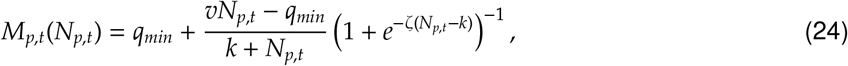

*where q*_*min*_ is the lower bound on predation rate when predator density approaches zero, *v* is the instantaneous rate of exchange from invulnerable to vulnerable states, *ζ* controls the steepness of the logistic model component, and *k* is the predator density level associated with the inflection point on the logistic component of the relationship.

### 2.4. Estimating equilibrium unfished biomass

We calculate unfished equilibrium spawning biomass *SB*_0_ for two scenarios for future *M* with *noPred* and one scenario of future *M* with *predM* models:

1. an average M (*noPred-avgM*) scenario using the average M over the full history (1951-2022) of 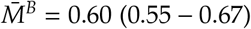 for all ages (posterior median with 95% CI);
2. a current M (*noPred-currM*) scenario using the average M in the last 10 years (2013-2022) of 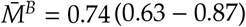 for all ages (posterior median with 95% CI)
3. a current M (*predM*) scenario using the average basal morality *M*^*B*^ in the last 10 years (2013-2022) of 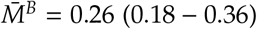 for all ages (posterior median with 95% CI), combined with projections of predator abundance to simulate predation mortality

The *SB*_0_ for i) is a leading parameter estimated in the SCAH model (EQ.1-5, Table A.2) and represents the standard WCVI Herring fishery stock assessment model assumptions for equilibrium states based on average historical ecosytem conditions, while ii) and iii) estimate *SB*_0_ for equilbrium states for future ecosystem conditions that reflect higher natural mortality rates observed in recent years. We use simulation to estimate *SB*_0_ for *noPred-currM* and *predM*, since analytical solutions based on average historical mortality represent states of WCVI Herring with lower predator abundance. Future *M*_*t*_ is modelled as a random walk

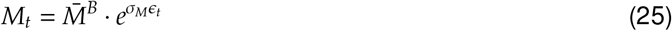

where *ϵ*_*t*_ ∼ N(0, 1) and *σ*_*M*_ = 0.1 is the random walk deviation, as in (1)

Simulation is also preferable for *predM* to analytical calculations based on estimated yield-per-recruit because per-capita predation mortality 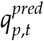 is related to vulnerable herring biomass via a functional response, making *M*_*a,p*_ difficult to solve analytically in the presence of changing fishing mortality. Simply fixing predation mortality (or the per-capita predation rate) to estimate reference points analytically would exclude the functional response relationship of predators to prey density, and would at times over- and under-represent future predation mortality.

WCVI Herring and all predator populations are projected 300 years into the future over 200 random seeds to estimate Pacific Herring equilibria in the absence of fishing. Each random seed is conditioned on a single draw of parameter values from the SCAH model posterior distribution. All Pacific Herring stock dynamics in the projection match the SCAH model specifications, with discrete removals of fertilised eggs by GW, and instantaneous predation mortality for hbW, hbS, HS, SSL, and both Pacific Hake size classes for *predM*, as described above. For Pacifc Herring and Pacific Hake, which are simulated using age-structured population dynamics models, future recruitments are simulated with random recruitment process errors in each replicate to simulate variability in future abundance. In addition, Pacific Hake biomass is simulated with a harvest rate of 30%, which is based on the *U*_*MSY*_ estimated from the operating model conditioned from the recent stock assessment (Johnson et al., 2021). Logistic models of marine mammal populations are also projected into the future, with future population size uncertainty incorporated via random sampling of parameters for the theta-logistic model parameters for all marine mammals (Supplementary B). For hbW, hbS, HS, and SSL, random parameters are drawn from estimated theta-logistic model posterior distributions, while parameters for GW are drawn from assumed distributions from Gavrilchuk and Doniol-Valcroze (2021).

Unfished equilibrium spawning biomass, mortality, and recruitment for each scenario are estimated for each random seed/posterior draw from long-term average simulated model states. For unfished spawning biomass 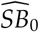 and unfished natural (basal + predation) mortality-at-age 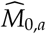 model states *SB*_*t*_ and *M*_*a,t*_ are averaged over 100 projection years *t* ∈ {2201, …, 2300}, i.e.

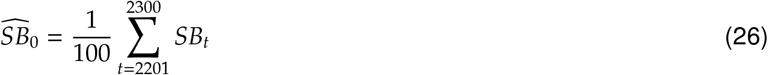

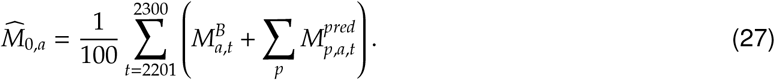

Average unfished recruitment 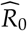 is then derived from 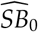 and 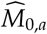 via spawning-stock-biomass-per-recruit as

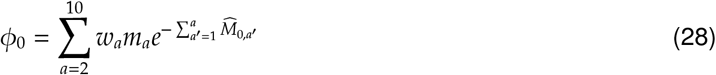

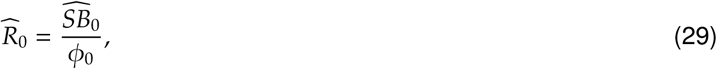

where *w*_*a*_ is mean weight-at-age for Pacific Herring, *m*_*a*_ is maturity-at-age, and survival is adjusted to include age-specific mortality 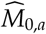 given that herring spawn at the end of the year. Note that *ϕ*_0_ calculation excludes age-1 herring, which are all assumed to be immature. The resulting estimates over 200 simulation replicates and posterior draws approximate the posterior distribution of each quantity. The 100-year period (2201-2300) used for equilibrium calculations begins after all predator populations have stabilized around their carrying capacity (Supplementary B), and provides sufficient time for Pacific Herring dynamics to equilibrate to the maximum assumed predation mortality.

## 3. Results

This section first describes the results of fitting the SCAH model to WCVI herring data under the *predM* and *noPredM* hypotheses, including a brief description of goodness of fit, current states, and unfished equilibrium states. The unfished equilibrium states reflect uncertainty in future predator population trajectories for the *predM* model and alternative future mortality scenarios (*currM, avgM*) for *noPredM*. Following this, we describe estimates of functional responses from foraging arena theory models, which are used to project predator dynamics for the *predM* model in simulations.

### 3.1. Current and equilibrium states

SCAH posterior estimates for 2022 spawning biomass are quite similar between the *noPredM* and *predM* hypotheses, with posterior medians of 20.5 kt (95% CI: 11.1-35.1) and 21.9 kt (95% CI: 11.5-38.8), respectively (Table 3). Although both models follow qualitatively similar trends over the entire time series, the median spawning biomass estimates from *predM* are on average 1.2 times that of *noPredM* for the 1951-2022 period. While *predM* and *noPredM* agree closely (+/- 20%) for most of the 1970-2022 period (Fig. 2), biomass estimates from *predM* are on average 1.5 times higher than those from *noPredM* during the earlier 1951 - 1969 period when surface spawn surveys were primarily used. The *predM* model also estimates higher biomass for the 2005-2013 period, where median spawning biomass for *predM* is on average 2.1 times that of *noPredM*. Strong agreement over most of the 1988-2022 period when the dive survey is the primary survey method (Fig. A.2) is unsurprising, given that the dive survey is treated as an absolute abundance index with catchability *q*_*d*_ = 1 (Appendix A).

**Figure 2:**
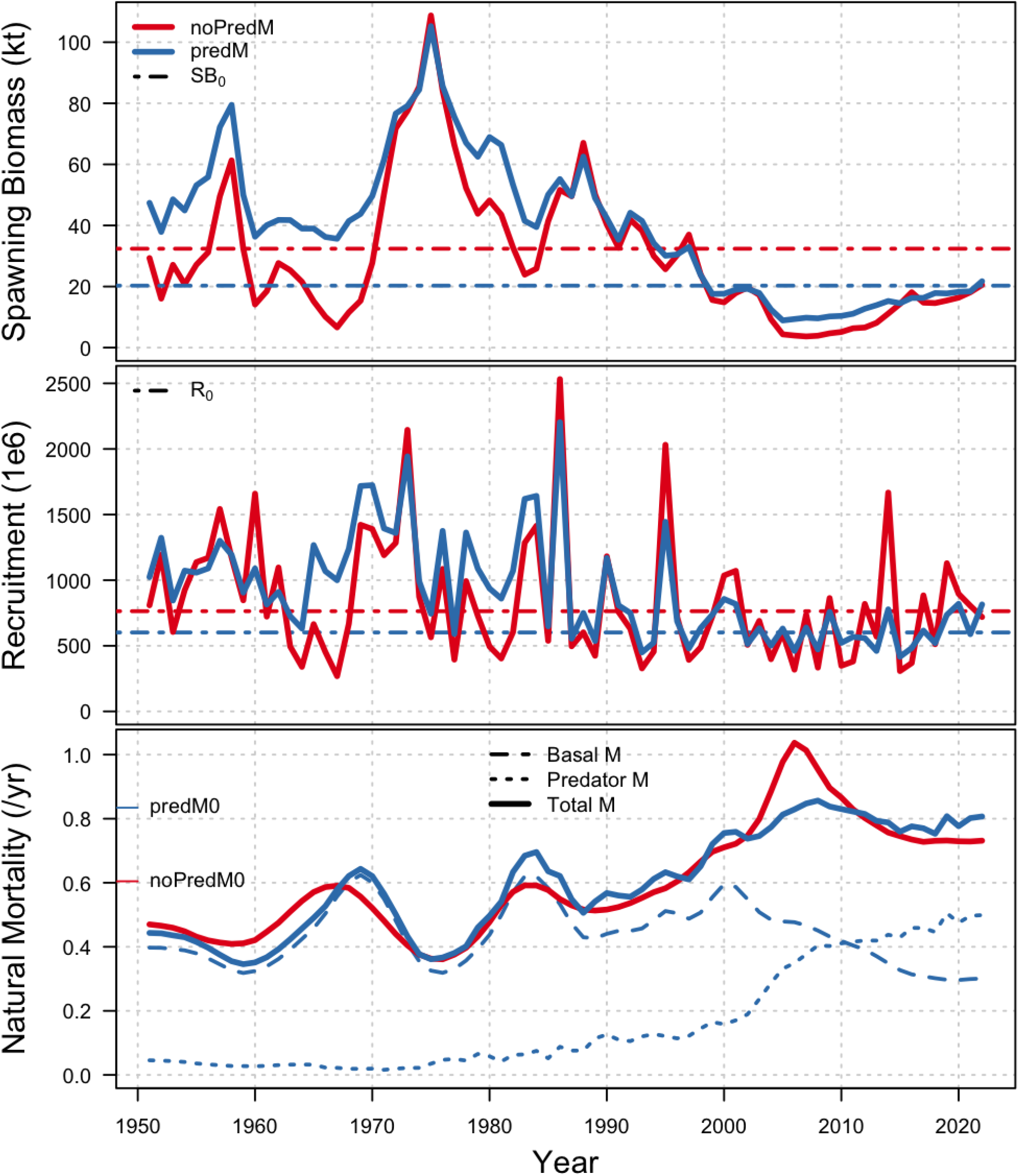
Posterior median spawning stock biomass (top), age-1 recruitment (middle), and age-2 natural mortality (bottom) estimates from *noPredM* (red) and *predM* (blue) SISCAH models. Unfished equilibrium states (*SB*_0_,*R*_0_,*M*_0_) for *noPred* are from the *avgM* projection scenario.

Despite strong agreement on historical biomass trends, the long-term average unfished values depend heavily on hypotheses about future natural mortality. The *noPredM-currM* and *predM* estimates for *SB*_0_ are quite similar, with posterior medians of 17.5 kt (95% CI: 8.8-28.8) and 20.3 kt (95% CI: 6.7-39.1), respectively (Table 3, Fig. 6). The 2022 spawning biomass is above *SB*_0_ for *noPredM-currM* and *predM* models with spawning biomass depletion estimates of 1.20 (95% CI: 0.71-2.45) for *noPredM-currM* and 1.09 (95% CI: 0.53-3.11) for *predM*. In contrast, *noPredM-avgM* estimates unfished equilibrium spawning biomass *SB*_0_ as 32.4 kt (95%CI: 11.1-35.1) with a 2022 spawning biomass depletion of 0.64 (95%CI: 0.34-1.1).

Time series of recruitment estimates under both *predM* and *noPredM* are typical of most fish stocks. Posterior median recruitments fluctuate above and below the unfished equilibrium level for both hypotheses (Fig. 2), with more recent estimates below unfished levels more often, commensurate with the historically low spawning biomass since 2005. Overall, the *predM* and *noPredM* recruitment time series follow qualitatively similar trends but the *noPredM* recruitments are higher on average (Table 3). The higher average recruitments under *noPredM* are due in part to the size-selectivity of predators under the *predM* model. Under *predM*, basal mortality applies to all fish but predation mortality rates are influenced by predator size-selectivity, with age-2 fish subject to the greatest mortality rates (Fig. A.5). Predation mortality for ages 3-10 is greatest from Winter Humpback Whales and Stellar Sea Lions, while predation mortality for age-1 is greatest from Summer Humpback Whales and Pacific Hake less than 50 cm. In contrast, mortality rates under the *noPredM* model apply to Pacific Herring of all ages, leading to lower survival rates for some age classes, driving recruitments higher on average.

Estimates of unfished equilibrium recruitment between hypotheses have a similar story to those for unfished biomass. Under the *predM* model, unfished recruitment *R*_0_ is 602 million (95%CI: 296-857) age-1 fish, around 21% lower than the 764 million (95%CI: 540-1084) under the *noPredM-avgM* scenario (Table 3). Under the higher future mortality in the *noPredM-currM* scenario, unfished recruitments drop to 702 million (95% CI: 450-1066).

Age-2 mortality estimates for both models follow similar trends from 1951-2022, with large increases in *M* begining in the mid 1990s (Fig. 2). Posterior median mortality for the *noPredM* model peaks in 2006 at 1.04, and then declines to a range of 0.73-0.78 over the last decade (2013-2022). Similarly, posterior median age-2 mortality for *predM* increases to a peak of 0.95 in 2007, before declining to a range of 0.75-0.80 from 2013-2022. Age-2 predation mortality under the *predM* model shows an increasing trend over the last decade that is expected to continue in the future, given that some predator populations are expected to continue growing (Supplementary B). The *predM* models has higher interannual variability in mortality, which is influenced by annual changes in predator consumption, particularly Pacific Hake.

Both model hypotheses indicate a sustained increase in mortality rates beginning in the 1990s, with the *predM* model attributing the majority of the increase to higher consumption by growing marine mammal populations (Fig. 3). Over the same time, the posterior median basal mortality rate declines from a maximum of 0.52 in 2000 to 0.26 from 2015-2022 to accommodate the large increases in predation mortality rates. Humpback Whales were the largest source of Pacific Herring predation over last decade (2013-2022), with average annual consumption estimates from bioenergetic models of 6.9 kt for winter and 2.2 kt for summer feeding groups (Fig. 4). Humpback Whales account for 62% of total predator consumption of Pacific Herring (excluding eggs consumed by grey whales) over that period. Other sources of predation mortality over the last decade were small in comparison, which included mean annual consumption of 2.4 kt by Pacific Hake, 2.3 kt by Steller Sea Lions, and 0.8 kt by Harbour Seals. The average Grey Whale egg consumption during the same period removes the eggs produced by 2.5-3.4 kt of spawning biomass, which corresponds to removals of 14-28% of annual egg production from 2013-2022.

**Figure 3:**
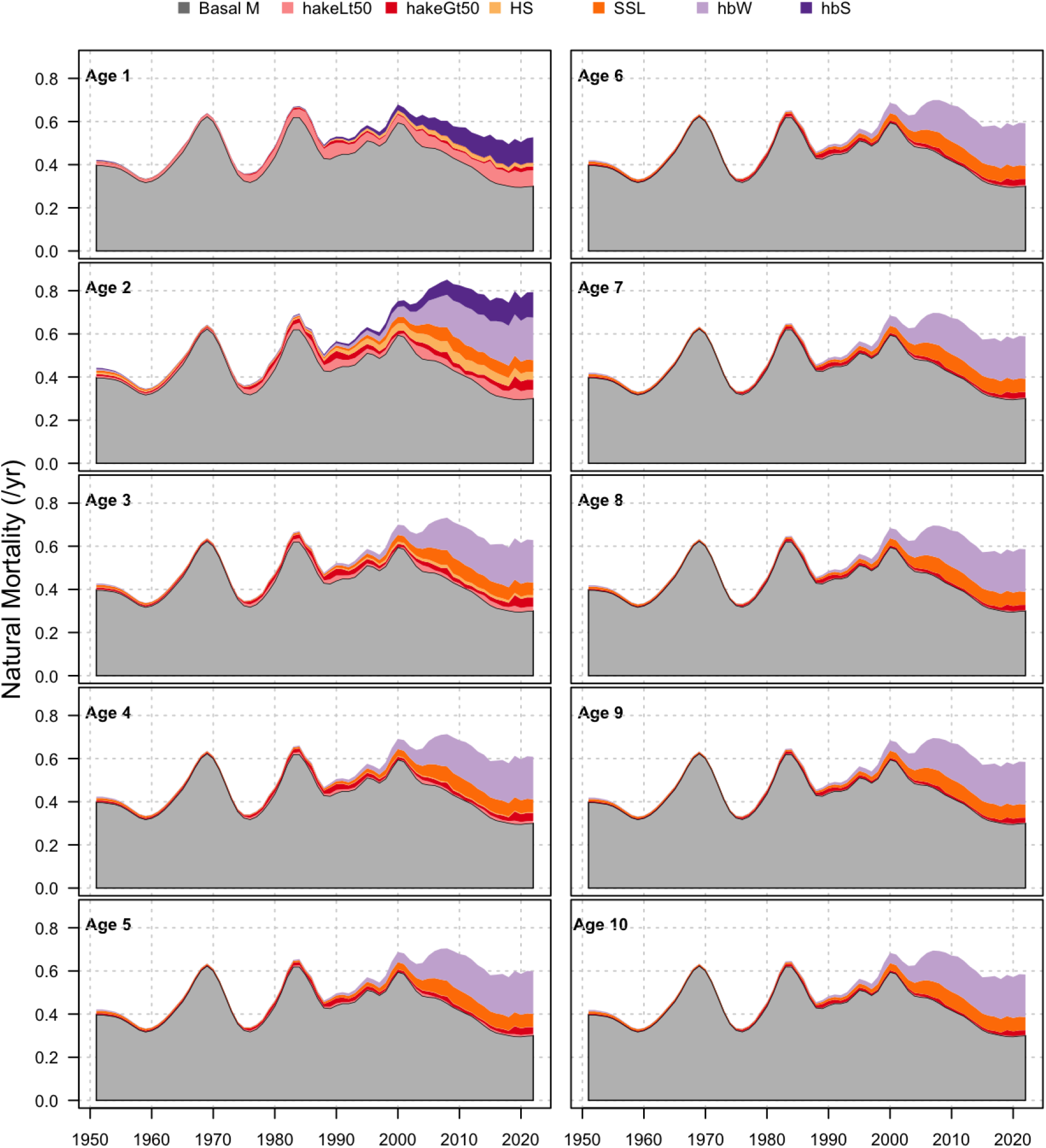
Basal and predator components of natural mortality for ages 1-10 in *predM* model.

**Figure 4:**
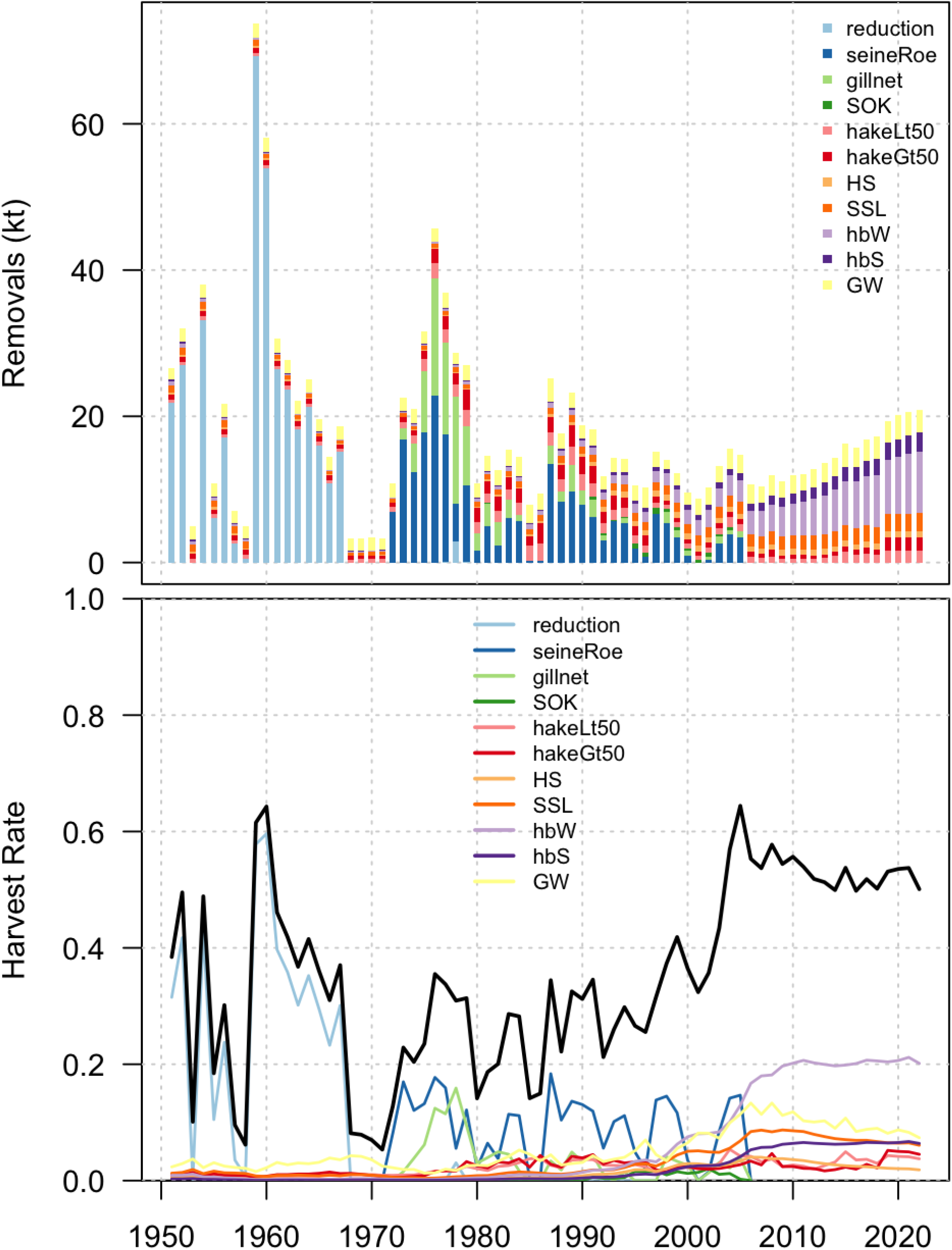
Removal and harvest rate for fisheries and predators in *predM* model. Removals (*C*_*t*_) include total ponded biomass for SOK fisheries and equivalent ponded biomass for Grey Whale egg consumption. Harvest rates are calculated as total removals divided by the sum of spawning biomass at the end of year and total removals, i.e. *C*_*t*_*/*(*SB*_*t*_ + *C*_*t*_). Black line in bottom panel indicates total harvest rates across all fisheries and predators.

**Figure 5:**
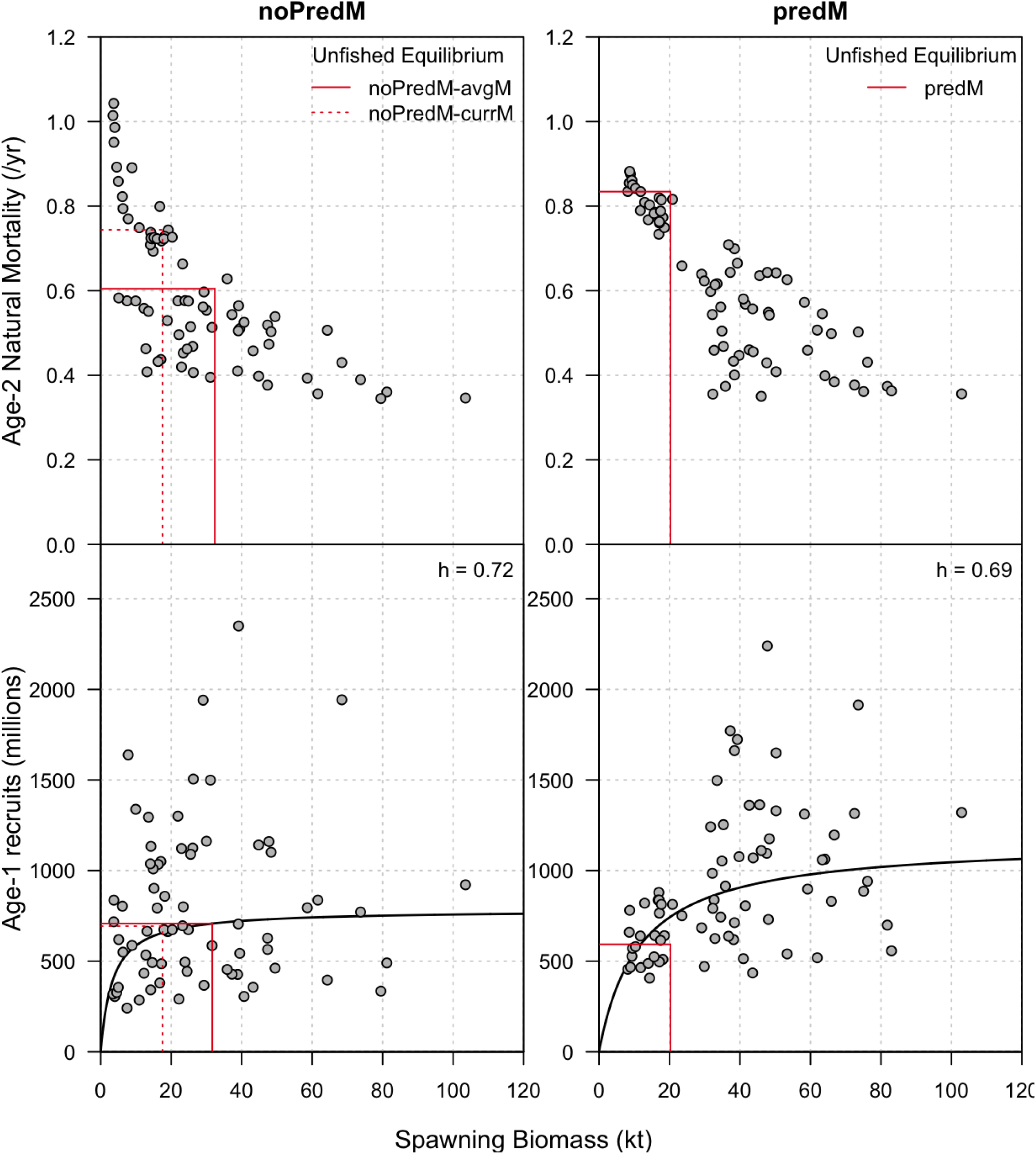
Total age-2 natural mortality (basal + predation mortality, top) and recruitment (bottom) relationships with spawning biomass for *noPredM* (left) and *predM* (right) models. Solid and dotted red lines indicate estimates of unfished equilibrium age-2 mortality, recruitment, and spawning biomass. Solid black lines in bottom panel are the estimated stock-recruit curves.

### 3.2. Goodness of fit for SCAH model

Overall, both model hypotheses fit the spawn index and age composition data well (Fig. A.3-A.4), with slightly better fits for the *noPredM* model (Table 2). The *noPredM* model has lower residual standard errors for the dive survey index, while standard errors are similar between both models for the surface survey. Surface survey catchability is 40% lower under the *predM* hypothesis, indicating slightly higher biomass during the pre-1970 period (Fig. A.3), but this does not appear to significantly affect goodness of fit to the spawn indices. Spawning biomass estimates appear unbiased overall for both hypotheses, with mean residuals at or very close to zero, and no significant residual trends (*p >* 0.05) over the 1951 - 2022 period. The age sampling standard errors are similar between both models for seine-roe and gillnet ages, while the *noPredM* model has better fits for reduction fishery ages (Table 2). The latter is largely related to a persistent under-estimate of age-3 fish in the reduction fishery in the *predM* model, and a compensatory over-estimate of ages 5+ (Fig. A.4). This is likely related to differences in how mortality is broken up across basal and predator sources, and how predation mortality is assigned to age classes via selectivity functions. Even so, the statistical properties of the recruitment process errors do not indicate any red flags, with all deviations within 3 standard errors of zero, mean residuals close to zero, and no significant time trend (Fig. A.3). Residual patterns in recruitment process errors do not show temporal trends with *ρ* = 0.09 for *noPredM* and *ρ* = 0.12 for *predM* for 1-year lag autocorrelation.

**Table 2:**
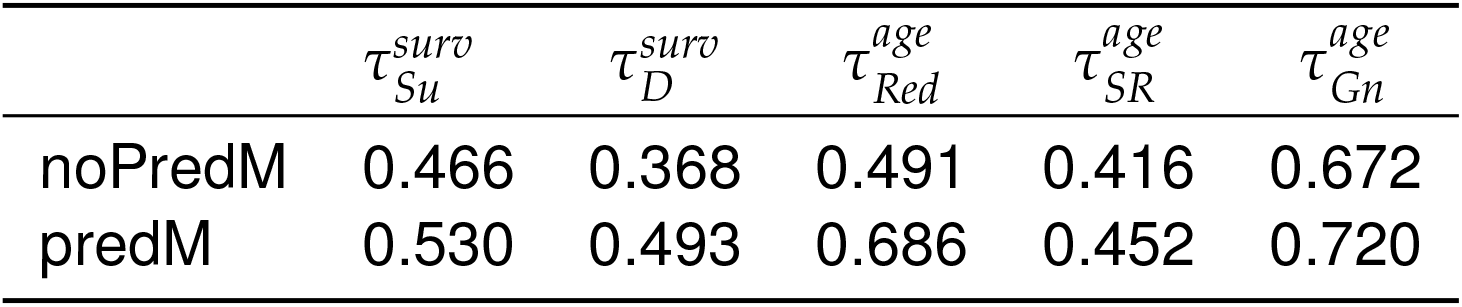
SISCAH model data likelihood function standard errors for the *noPredM* and *predM* hypotheses. The first two columns show the log-normal spawn index residual standard errors for each component of the blended spawn survey, and the last three columns show age data sampling error standard deviations from the logistic-normal compositional likelihood function.

**Table 3:**
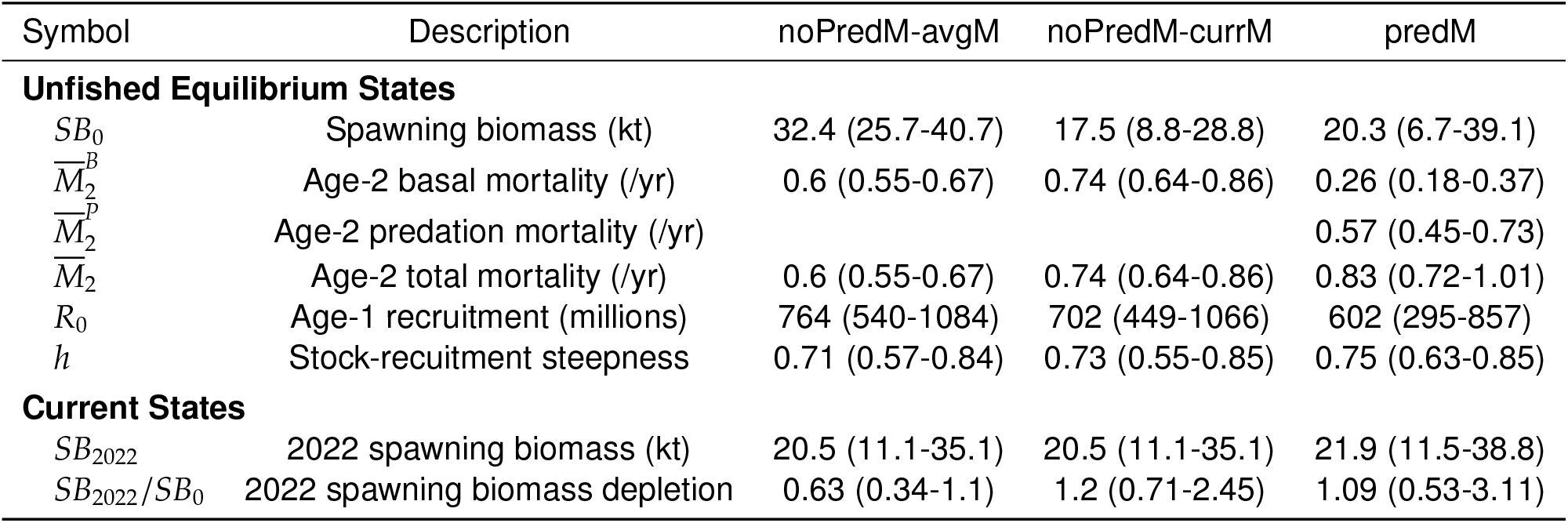
Posterior median with 95% CIs for WCVI Herring spawning biomass and life-history parameters for current and unfished equilibrium states. Unfished equilibria and depletion estimates for *noPredM-currM* and *predM* are calculated via simulation, while *noPredM-currM* are estimates from SCAH model.

| Symbol | Description | noPredM-avgM | noPredM-currM | predM |
| --- | --- | --- | --- | --- |
| **Unfished Equilibrium States** |  |  |  |  |
| $SB_0$ | Spawning biomass (kt) | 32.4 (25.7-40.7) | 17.5 (8.8-28.8) | 20.3 (6.7-39.1) |
| $\overline{M}_2^B$ | Age-2 basal mortality (/yr) | 0.6 (0.55-0.67) | 0.74 (0.64-0.86) | 0.26 (0.18-0.37) |
| $\overline{M}_2^P$ | Age-2 predation mortality (/yr) | | | 0.57 (0.45-0.73) |
| $\overline{M}_2$ | Age-2 total mortality (/yr) | 0.6 (0.55-0.67) | 0.74 (0.64-0.86) | 0.83 (0.72-1.01) |
| $R_0$ | Age-1 recruitment (millions) | 764 (540-1084) | 702 (449-1066) | 602 (295-857) |
| $h$ | Stock-recruitment steepness | 0.71 (0.57-0.84) | 0.73 (0.55-0.85) | 0.75 (0.63-0.85) |
| **Current States** |  |  |  |  |
| $SB_{2022}$ | 2022 spawning biomass (kt) | 20.5 (11.1-35.1) | 20.5 (11.1-35.1) | 21.9 (11.5-38.8) |
| $SB_{2022}/SB_0$ | 2022 spawning biomass depletion | 0.63 (0.34-1.1) | 1.2 (0.71-2.45) | 1.09 (0.53-3.11) |

### 3.3. Predator functional responses

Functional response and foraging arena theory models are post-fit to SCAH estimates of per-capita predation rates for each predator. Overall, the estimated models do sufficiently well at capturing the relationship between predation mortality, prey density, and predator density over the ranges that will be important for simulations of future predator consumption.

Functional response models of Pacific Hake and Harbour Seal predation rates tracked the SCAH model estimates of per-capita predation rates quite well (Fig. 7). As expected, there is some noise in the relationship, but the general shape of the data is matched by the estimated curves. All three curves have no observations at the lowest herring biomass, which means there aren’t observations at the the peak per-capita predation rate for the Pacific Hake predator classes with Type III responses.

**Figure 6:**
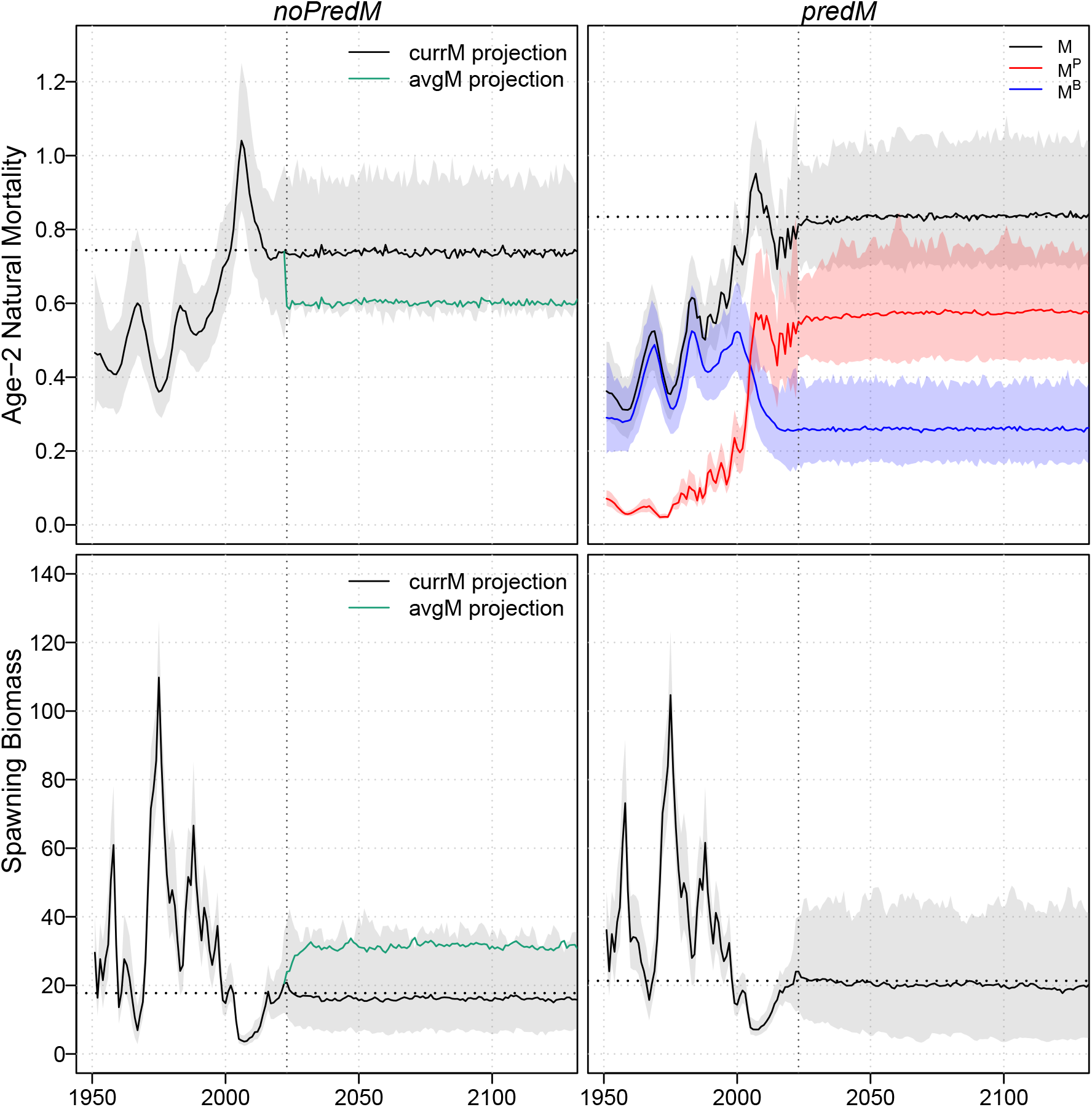
Median (solid lines) and central 95% (envelopes) of posterior distributions for spawning biomass *SB* (top) and age-2 natural mortality *M*_2_ (bottom) for *noPredM* (left) and *predM* (right) models for historical period (1951-2022) and projections (2022-2150) with no fishing. Natural mortality *M*_2_ rates for *predM* (bottom right) is separated into components for predation mortality 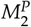 and basal mortality 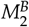. Vertical dotted grey line indicates the start of the projection period, while the horizontal dotted black line indicates median estimates of unfished equilibria states for spawning biomass *SB*_0_ and age-2 mortality 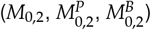from projections.

**Figure 7:**
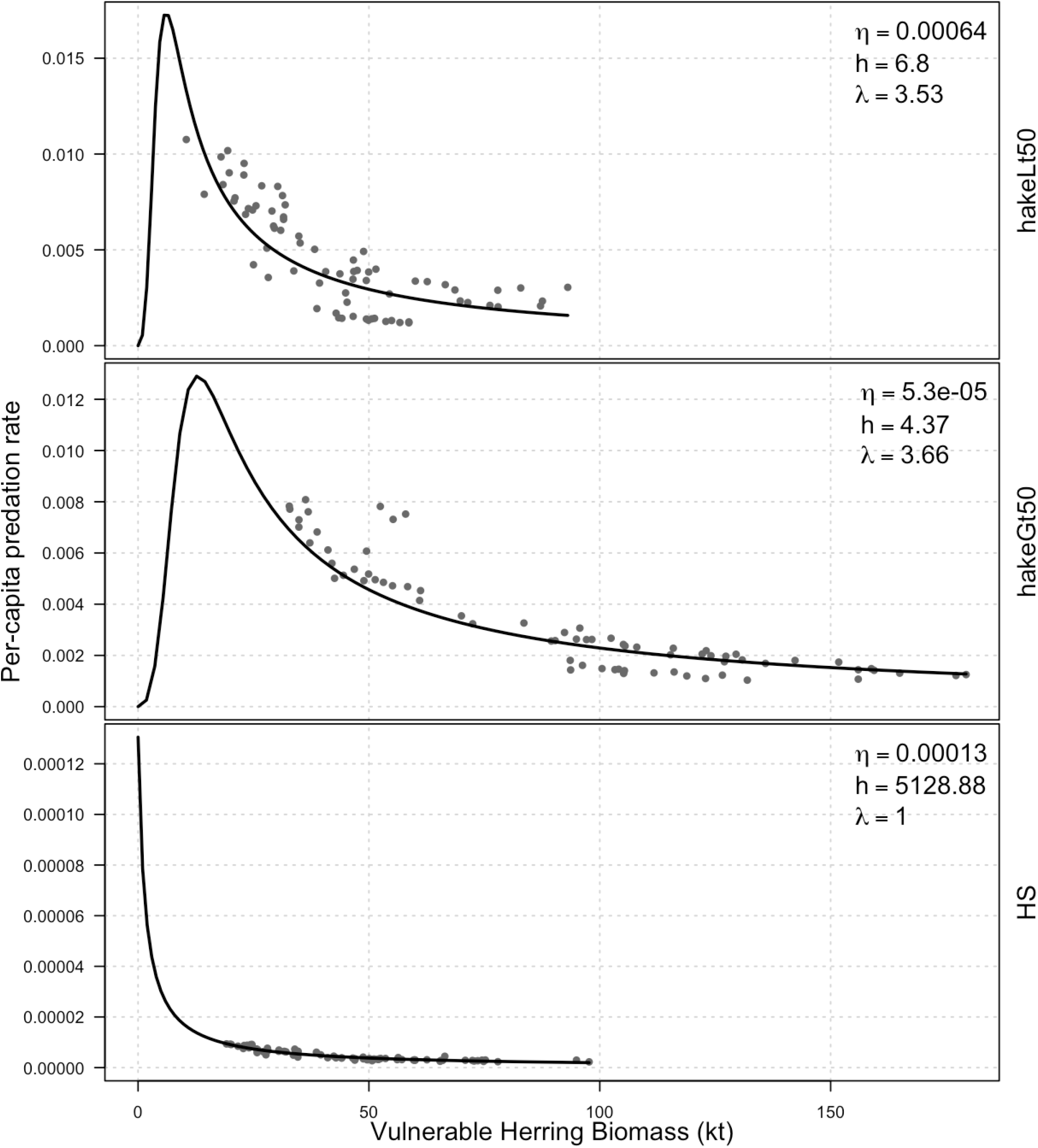
Functional response relationships between per-capita predation rate (y-axis) and herring vulnerable biomass (x-axis, MLE) for hakeLt50 (top), hakeGt50 (middle), and HS (bottom).

Foraging arena theory models had more variability and residual error with the SCAH model estimates. For SSL and hbW predator classes, the foraging arena theory models of predation mortality rate *M*_*p,t*_ fit best at low predator abundance (Fig. 8), but as predator density increases there is the perception of higher residual error. Some of the error is due to a log-normal distribution used in the model’s objective function, leading to proportional errors, but there is some indication that SCAH estimates may have a different functional form than the modified foraging arena theory model. Specifically, there is a high peak and then reduction in SCAH estimates for SSL that does not align with the foraging arena theory model neither for absolute predation mortality nor per-capita predation rates (Fig. 8, left hand column). Additionally, for all three predator classes there is a ‘bunching up’ effect, where a wide range of SCAH estimates of per-capita predation mortality rates occur over a narrow range of low predator abundances (Fig. 8, bottom row). Either of these two residual patterns could indicate model mis-specification. One possible source of mis-specification is a variability mismatch between predator population dynamics models, which are deterministic, and the Pacific Herring age-structured model, which has several sources of process variation in mortality and recruitment. For SSL specifically, the modification made to add a sigmoidal shape to the predation mortality rates may be overly restrictive on the per-capita predation rate scale, leading to the residual pattern and peak mis-alignment.

**Figure 8:**
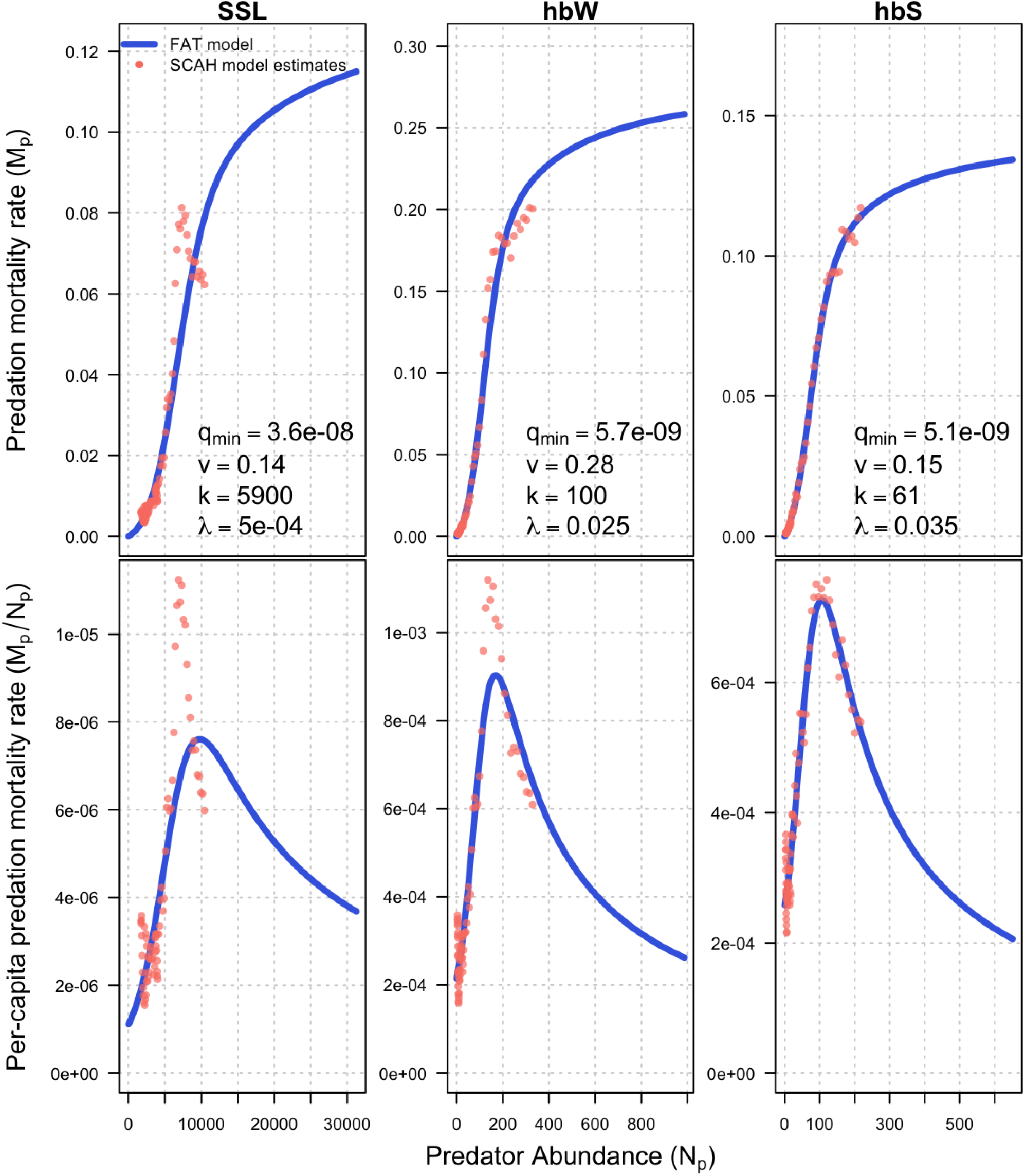
Foraging arena theory (FAT) relationships between predation rate (top row) and predator density (x-axis), and per-capita predation rate (bottom row) fit to SCAH MLE estimates (pink points) for SSL, hbW, and hbS predators.

For functional response curves, the lack of observations at low herring biomass means there is high uncertainty about the functional response of predators at low herring stock sizes; however, the relative impact of Pacific Hake and Harbour Seals is low, so the bias is not expected to have a large impact. For foraging arena theory models, most of the model mis-specification is at lower predator abundances than current levels, which have low probability of occurring in the future. Barring an unusual mortality event that reduces populations to mid-20th century levels, Steller Sea Lions and Humpback Whales should start approaching carrying capacity in the WCVI area. While there is also evidence of model mis-specification at higher predator abundance for SSL, the general behaviour of the model is desirable for the purpose of estimating unfished Pacific Herring biomass under predation.

## 4. Discussion

When predator consumption of fished species increases, there are important implications for fisheries assessment and management systems. In particular, marine mammal populations have experienced large population increases in many regions following bans on commercial harvest and predator control programs (Magera et al., 2013; Ford et al., 2009; Olesiuk, 2009, 2018). In some areas, the resulting increases are highly correlated with declines in their prey species (Swain and Benoit, 2015). Such noticeable patterns have led to hypotheses that increased predation by marine mammals is the primary cause for productivity declines in some fish stocks and their lack of recovery to historical abundance levels (Swain and Benoit, 2015; Nelson, 2020; Walters et al., 2020). In this study, we evaluate such a hypothesis for WCVI Pacific Herring by quantifying historical predation rates by key predators including cetaceans (Pacific Humpback, Grey Whales), pinnipeds (Stellar Sea Lions, Harbour Seals), and piscivorous fish (Pacific Hake). Predation mortality is quantified via a novel approach that integrates i) a single species catch-at-age assessment model for Herring, ii) predator abundance models for the 5 predators, and iii) bioenergetic models to estimates consumption rates for each predator.

Our main findings, discussed below, are two-fold. First, we found that predator consumption, largely driven by Humpback Whales, did indeed explain increasing WCVI Herring natural mortality rates in recent years and could be used to forecast future mortality rates. Second, we found that estimates of stock status and expectations for future biomass trajectories are contingent on assumptions about future natural mortality rates, extending and corroborating previous findings (Benson et al., 2023).

Indeed, post-hoc analysis of natural mortality rates and beginning-of-year age 2+ herring biomass estimated in fishery stock assessments suggests a depensatory relationship between natural mortality and age-2+ biomass (Johnson et al., 2024), which is characterised by higher mortality at low biomass levels. Such depensatory natural mortality is typical of type II or III functional response of predation mortality to prey density (Holling, 1959).

Our findings align with previous research using multi-species models for fisheries stock assessment, which found that ignoring predation or mispecifying future trends in natural mortality can produce large biases in model predictions, perception of stock status, and unfished equilibrium biomass (Trijoulet et al., 2020). Ignoring predation mortality may also lead to biased estimates of catchability *q* and absolute spawning biomass (Trijoulet et al., 2020); however, in our study this did not occur because the dive survey catchability is fixed at *q*_*d*_ = 1.

### 4.1. Natural mortality trends

Accounting for predation mortality (*predM*) produced similar trends in total natural mortality (basal + predation mortality) from 1951-2022 compared to mortality estimates obtained using a random walk without a predation component (*noPredM*). This indicates that predation mortality could be an important factor driving recent increases in WCVI Pacific Herring mortality, and offers a new approach to forecasting future mortality. Additionally, estimates of WCVI Herring mortality caused by major predators were found to have risen sharply with increasing predator populations since the 1990s. Since 2005, annual predator consumption rates of Herring have exceeded the total fisheries harvest rates during the 1950s and 1960s, when the BC reduction fishery experienced peak catches (Fig. 4). Previous research has identified Harbour Seals, Stellar Sea Lions, Pacific Hake, and Humpback Whales as significant predators of Herring (Tanasichuk et al., 1991; Olesiuk, 1993; McMillan, 2014; Tollit et al., 2015); however, specific estimates of predation mortality from each predator are lacking. This study suggests a key finding that Humpback Whales are the primary source of predation for WCVI Herring. Indeed, bioenergetic model estimates suggest that Humpback Whales consume more Herring than all other predators combined, accounting for 60-66% of the annual predator consumption over the last decade.

Looking forward, it is expected that the majority of Pacific Herring predation mortality will continue be driven by Humpback Whales. Although some recent estimates suggest North Pacific Humpback Whale populations may be nearing carrying capacity (Cheeseman et al., 2024), their carrying capacity and future growth remains uncertain. The regional model used here suggests a high probability that the population will continue to grow in British Columbia (Appendix B), in line with previous growth trends for the region (Ford et al., 2009; Calambokidis and Barlow, 2020). Such information is valuable for allocating resources in future Pacific Herring fishery science and management; it would be valuable to improve estimates of one or more factors that determine the scale of Humpback Whale predation mortality, such as Humpback Whale population size in BC, the proportion feeding in WCVI during different times of the year, their diet composition, and foraging behaviour. For example, there may be significant unobserved foraging by Humpback Whales on feeding aggregations of age-3+ Herring on La Perouse bank during the summer months, which is not captured here. Given the scale of Humpback Whale consumption, a shift in the fully selected age classes during the summer would likely have noticeable effects on estimates of Pacific Herring productivity parameters.

### 4.2. Implications for stock status and recovery trajectory

Estimates of stock status and future equilibrium states were highly sensitive to assumptions about future Pacific Herring natural mortality. Equilibrium states that reflect recent trends of higher natural mortality, whether by using an average mortality from the last 10 years (*noPredM-currM*) or forecasts of future predation mortality (*predM*), resulted in lower estimates of equilibrium biomass. Both (*noPredM-currM*) and (*predM*) models indicated that the 2022 stock status was above unfished biomass at 1.2*B*_0_ and 1.09*B*_0_, respectively. In contrast the *noPredM-avgM* model, which assumes future mortality varies around the long-term average, suggests a 2022 stock status of 0.63*B*_0_. Therefore, the choice of model has important implications for the WCVI Herring fisheries management system. The *noPredM-avgM* stock status could lead to more conservative harvest rates, with unrealistic expectations for future spawning biomass in the absence of fishing, which is unlikely to occur if recent natural mortality rates persist (Fig. 6). Consequently, in such a scenario the stock may fail to meet conservation objectives due to over-estimates of unfished biomass that expect future productivity regimes to be like the 1950-1990 period (e.g., DFO, 2021).

### 4.3. Next steps: accounting for changing future natural mortality in MSE

In this study, we present a novel method for linking single species fisheries stock assessments with predator abundance and bioenergetic models. We demonstrate how this approach can be used to forecast future natural mortality rates and unfished equilibrium states via simulation. A potential next step would be to integrate this approach into management advice via management strategy evaluation (MSE), by using closed-loop simulations to evaluate candidate harvest strategies performance for achieving Pacific Herring fisheries and ecosystem objectives.

Our results indicate that incorporating predator-prey dynamics to account for predation mortality rates can provide estimates of current stock status and simulated future biomass that may better reflect present fishery productivity. This is particularly the case for fish stocks where predator abundance in recent years differs from the historical assessment period, such as for fisheries on forage fish species close to increasing marine mammal populations on the east and west coasts of North America. Our study provides evidence that the high natural mortality regime for WCVI Herring from the last 20 years is likely influenced by increased predation mortality, which is expected to continue in the future. Under the current ecosystem conditions it appears unlikely that future WCVI Herring biomass will increase and reach historical biomass levels observed prior to mid 1990s. This finding has the potential to adjust perspectives for WCVI Pacific Herring management and manage stakeholder expectations for future biomass. For example, a biomass target was recently proposed for WCVI Herring based on a productive period in the recent history that aims to: “*maintain spawning stock biomass at or above a target biomass level equivalent to the average biomass from 1990-1999, with at least 75% probability over two herring generations*” (DFO, 2021). When tested in simulation, this conservation objective was achievable under projection scenarios that used the historical average mortality rates (i.e., similar to the *noPredM* model), but was not achievable under any scenario that assumed that future herring mortality rates remain elevated (i.e., similar to *noPredM-currM* and *predM*) like recent ecosystem conditions suggest (DFO, 2021). Therefore, the most significant factor in performance metrics for simulated harvest strategies was the assumption about future natural mortality, which poses a challenge for fisheries management decisions. Ecological models that link projections of future ecosystem conditions with fish population dynamics, such as those used here, can provide more informed choices for modelling future population trajectories.

One of the challenges with developing multi-species models that directly estimate predation mortality is that key information for predators and bioenergetic modelling are often lacking. For the key predators included in this study, only Pacific Hake had an existing age-structured model (Johnson et al., 2021), whereas population models for marine mammal predators needed to be developed from multiple data sources (Appendix B). Furthermore, bioenergetic models are highly sensitive to parameter assumptions (Moran et al., 2018), especially in cases where essential data, such as diet composition, foraging periods, and metabolic requirements, are lacking. This raises the question: can similar results for management advice be achieved with simpler models?

Practical applications for fisheries management involve striking a balance between costs (e.g., data collection, analyst time) and model complexity. This may explain why there are relatively few examples of integrating multispecies models into MSE and using them to provide tactical fisheries management advice (Punt et al., 2016; Johnson and Cox, 2021; Lucey et al., 2021; Howell et al., 2021). The need for a MICE model depends on the specific fisheries management goals, encompassing single species and broader ecosystem considerations. Another consideration is that developing more complex models by including multiple species, species interactions (such as predation or competition), or ecosystem processes may not always produce more reliable results. Models with increased complexity usually rest on more assumptions to promote model convergence given data limitations about interactions among species, and every new species factorially increases the number of potential interactions. Relatedly, higher model complexity requires longer development and analysis timelines, which may not align with management budgets and delay timely management advice. In some cases, it may be more pragmatic to modify single-species model assumptions to capture predator-prey dynamics or other ecosystem conditions affecting stock productivity via new assumptions for mortality or recruitment relationships. For example, our *noPredM-currM* scenario that assumes future natural mortality will reflect the average from the last 10 years (2013-2022), produced similar results for depletion and unfished states to the *predM* model, and similar models have already been used in harvest strategy testing for Pacific Herring in BC (Benson et al., 2023).

It should be noted that greater differences between *noPredM-currM* and *predM* models might have occured if several predator populations were not approaching carrying capacity. If instead predators had required a longer time frame to reach carrying capacity, future predator *M* might differ significantly from the recent 10-year average. For example, if projections had started in 1990, when predator populations were lower and predation mortality was around 0.13 /yr, then the *noPred-currM* model would have over-estimated carrying capacity and future biomass. Alternatively, simpler MICE models might be sufficient, such as those that focus solely on the most crucial drivers of predation. For instance, our *predM* model might achieve similar results if we only included Humpback Whale predation and did not directly estimate predation from Harbor Seals, Stellar Sea Lions, Grey Whales, and Pacific Hake, instead relying on the time-varying basal mortality to capture their influence. The challenge with choosing simpler subsets of species lies in identifying the main drivers of predation, which were not initially known before fitting the *predM* model and developing bioenergetic models. Nonetheless, future iterations or utilization of the SCAH model could prioritize Humpback Whale predation mortality, given that they dominate predator consumption, and rely on simpler models for the remaining predators. Another promising approach to account for predator-prey dynamics involves incorporating predator type II and III functional responses into single species models through a depensatory natural mortality model (Holling, 1959; Neuenhoff et al., 2019; Johnson et al., 2024). Future work could assess the benefits of using multiple models with varying complexity to provide tactical management advice by comparing outcomes when used for biomass estimation and as operating models within an MSE simulation framework.

### 4.4. Conclusions

In conclusion, this study quantifies the impact of predation mortality for WCVI Pacific Herring and reveals that including predatory-prey dynamics changes perception of stock status and expectations for future biomass. When models incorporate higher future natural mortality, reflecting the current ecosystem conditions, estimates of current stock status were higher based on lower estimates of equilibrium unfished biomass. Conversely, models that assumed mortality was more like the historical average had lower stock status and higher unfished biomass estimates.

Our novel approach for ecosystem modelling links a single-species catch-at-age model with estimates of predator consumption derived from bioenergetic models. This framework provides fertile ground for developing operating model scenarios for future harvest strategy development and MSE work, improving scientific defensibility by removing an element of analyst choice for future mortality scenarios. However, given the complexity and time-cost of developing ecosystem and multi-species models, analysts should seriously consider if simpler models that capture the same dynamics implicitly are better suited for tactical needs, such as annual catch limit calculations. Key considerations for appropriate model complexity include management goals, data availability, research questions, timeliness of advice, and budget constraints. Simpler modifications to single-species model assumptions can be more pragmatic for providing fisheries management advice, while more complex multi-species or ecosystem models might provide more nuanced insights for exploring research questions related to multi-species ecosystems and fisheries interactions.

## 5. Supplementary material

Supplementary material includes a description of the statistical catch-at-age (SCAH) herring model (supplementary material A) and predator abundance models (supplementary material B)

## 6. Acknowledgements

The authors would like to thank the Nuu-chah-nulth Tribal Council (NTC) for their feedback throughout this project, including conceptualization of research goals. Funding was provided by the British Columbia Salmon Restoration and Innovation Fund (BCSRIF) to the NTC. This research was also made possible by the numerous staff from Fisheries and Oceans Canada who assist with data management and analysis for the Pacific Herring and marine mammal research programs.

## 7. Author contributions

Conceptualization: BD, SDNJ, AJB, SPC, JSC, JL Data curation: BD, SDNJ, JSC Formal analysis: BD, SDNJ Funding acquisition: AJB, JL Methodology: BD, SDNJ, AJB, SPC Project administration: AJB Software: BD, SDNJ Supervision: AJB, SPC, JSC, JL Visualization: BD, SDNJ Writing - original draft: BD, SDNJ Writing - review & editing: BD, SDNJ, AJB, SPC, JSC, JL

## 8. Conflict of interest

The authors declare no conflicts of interest.

## 9. Data availability statement

Pacific Herring data (spawn index, biosamples, catch) and marine mammal (SSL, HS) census data are available on the Government of Canada’s open data portal: https://search.open.canada.ca/.

## Notes

### Competing Interest Statement

The authors have declared no competing interest.

