## Supplementary Material for "Influence of predation mortality on past and future dynamics of Pacific Herring: implications for stock status and future biomass"

### Supplementary Material A: statistical catch-at-age (SCAH) herring model

This appendix describes the statistical catch-at-age Herring (SCAH) model (Johnson et al., 2024) used in the main body of this paper, including notation (Table A.1), process model equations (Table A.2), and observation models and objective functions (Table A.3).

SCAH parameters, equilibrium states, and full state dynamics are given in Table A.2. Model parameters (Table A.2, P.1 - P.4) are partitioned into four subsets consisting of leading parameters $\Theta^{lead}$, nuisance catchability and variance parameters $\Theta^{cond}$ estimated conditionally on the values of leading parameters, fixed parameters $\Theta^{fixed}$ for maturity and stock-recruit steepness, prior distribution hyperparameters $\Theta^{prior}$ for leading parameters.

Equilibrium states for recruitment (EQ.4) and numbers-at-age (EQ.5) are derived via spawning biomass per recruit (EQ.3), which is itself a function of time-averaged natural mortality, weight-at-age, maturity-at-age (EQ.1), and unfished equilibrium survivorship-at-age (EQ.2).

In SCAH, an annual time step $t$ runs from July 1 - June 30 with spawning occurring at the end of each year. Annual recruitment $R_t$ (occurring on July 1) is estimated as an average recruitment $\bar{R}$ with recruitment deviations $\omega_t$, with an additional prior mean of the determinstic recruitment $\hat{R}_t$ from a Beverton-Holt stock-recruitment relationship. Deterministic prior mean recruitments are parameterized via a stock-recruit steepness $h$, unfished spawning stock biomass $B_0$, and time-varying basal natural mortality rates $M_t^B$, which are formulated as a random walk with log-normal deviations, i.e.

$$M_t^B = M_{t-1}^B \cdot e^{\sigma_M \epsilon_t} \tag{1}$$

where $M_t^B$ is the instantaneous basal natural mortality rate at time $t$, and $\epsilon_t \sim \mathcal{N}(0, 1)$ are simple random

walk deviations with $\sigma_M = 0.1$. The intial basal mortality $M_0^B$ for $t = 1$ has a normal hyperprior, with a mean $m_M = 0.4$ and a standard deviation of $s_M = .4$.

Age-class abundances are initialised in a fished state by estimating initialisation multipliers $\omega_{init,a,2}$ for each age-class $a \in \{1, ..., 10\}$ in the first year. Initalisation multipliers scale the unfished equilibrium age-structure to non-equilibrium fished age-structure (Table A.2, NEQ.3), which in turn is scaled by either an estimated initial recruitment $R_{init}$ (predM) or the overall average recruitment $\bar{R}$ (noPredM). Estimating initialisation multipliers is equivalent to estimating recruitment deviations for the 10 years prior to model initialisation.

#### *A.1. Fisheries*

Roe fisheries (seine-roe, gillnet, spawn-on-kelp) target spawning aggregations just before spawning occurs between March 1 and April 15, while the reduction fishery historically occured a few months earlier in the winter. Removals from reduction, seine-roe, gillnet, andspawn-on-kelp (SOK) fisheries are represented as discrete fisheries occurring at a fractional time of year $\delta_g$ (Table A.2, C.1 - C.9), where $\delta_1 = 0.50$ for the reduction fishery, $\delta_2 = 0.65$ for the seine-roe fishery, $\delta_3 = 0.67$ for the gillnet fishery, and $\delta_6 = 0.68$ for the SOK fishery. Pacific Herring removals are calculated by converting landed commericial catch to landed numbers-at-age via the annual empirical weight-at-age observations (C.7), and then removing these numbers-at-age from the vulnerable numbers-at-age (C.8). Annual harvest rates by commercial fisheries $U_{g,t}$ are calculated as the ratio of landed catch to vulnerable biomass (Table A.2, C.9).

Age-selectivity in each fishery is modeled as a logistic function of age (Table A.2, C.2), with parameters age-at-50% s electivity $s_g^{50}$ and the difference between age-at-50% and age-at-95% $s_g^{Step}$. Selectivity-at-age 1 is set to 0 by default. There are log-normal priors on $s_g^{50}$ and $s_g^{Step}$, both with a standard deviation of 0.3, and ad-hoc prior mean values found via parameter tuning. Priors were needed to improve fits to age composition data under the predation model, where high rates of predation mortality on young fish increased recruitments, biasing catch-at-age estimates in preliminary model fits. Additionally, selectivity for the SOK fishery (see below) is assumed to be the same as the seine-roe fleet, because impounded fish are collected via purse-seine and there are no age composition observations for that fishery. Finally, selectivity curves for predators are fixed based on information on size or age-composition in diets obtained from the litterature, as described in the main text.

##### *A.1.1. Fertilised herring egg removals model*

Fertilised Pacific Herring egg removals are converted to numbers of herring at-age using the model equations K.1-K.6 in Table A.2. The first step involves defining a unitless conversion factor $\Psi_{g,t}$ (K.1) to convert between landed or consumed eggs and the equivalent biomass temporarily removed from the spawning stock to represent egg removals. The conversion factor $\Psi_{g,t}$ combines fecundity $\mathfrak{F} = 200$ eggs/gram of spawning biomass for an average female herring, the proportion of mature individuals in the pond $p_{m,t}$, the proportion of female individuals in the aggregate population $p_f = 0.5$, the proportion of effective spawners in the aggregate population $p_{eff} = .35$, and the average weight in grams of yield/consumption per unfertilised egg $\gamma_g$, which converted to spawn-on-kelp/bough product for the closed-pond spawn fishery with $\gamma_g = 0.0024$ (Schweigert et al., 2018), and fertilised egg weight for grey whale consumption with $\gamma_g = 0.0023$ (Bishop and Green, 2001). The proportion mature $p_{m,t}$ is the only time varying quantity, as it is based on the vulnerable Herring biomass age structure. A log-normal prior is assumed with $\Psi_{g,t} \sim \log N(0.06, 0.1)$, where $0.06$ is the average observed ratio between ponded biomass and landed SOK product (Schweigert et al., 2018).

The $\Psi_t$ conversion factor converted landed SOK product $K_t$ to an estimated biomass of ponded fish $P_{g,t}$ (Table A.2, K.2), and ponded biomass-at-age $P_{a,g,t}$ (Table A.2, K.3). Ponded biomass is converted to ponded numbers-at-age by scaling by weight-at-age. While grey whale egg consumption is not from ponded fish, we still calculate the equivalent 'ponded' fish for grey whale egg consumption since those fish are temporarily separated from the population for the purpose of spawning biomass calculations (Table A.2, K.4). Equivalent ponded fish are then returned to the general population (Table A.2, A.3) after applying a ponding-induced mortality rate $M_{SOK} = 0.315$ for SOK and the remainder of the natural and predation mortality for each model year (Table A.2, K.5) (Schweigert et al., 2018; Shields and Kingston, 1982).

#### *A.2. Observation models and likelihood functions*

Temporal variation in Pacific Herring stock abundance and population composition are monitored via surface and dive survey spawn survey indices, and three age composition series collected from the reduction, seine-roe, and gillnet commercial fisheries. There is also a purse-seine test fishery that is included in the seine-roe age data (Fig. A.1)

*A.2.0.1. Blended spawner index observation model.* Assumed observation models and statistical functions for likelihoods and prior distributions are given in Table A.3. Historically, the spawn survey was considered two separate indices: a pre-1987 surface survey and a post-1987 dive survey; however, the two designs have overlapped in several years (Fig. A.2). Here, we treat the two spawn survey designs as components of a single blended index, with a yearly catchability scalar derived via weighted averaging of surface and dive survey catchability parameters $q_s$, $q_d$, i.e.,

$$q_t = \eta_t q_s + (1 - \eta_t) q_d, \quad (2)$$

where the weights $\eta_t, 1 - \eta_t \in [0, 1]$ are the proportions of total observed spawn from the surface and dive surveys, respectively (Johnson et al., 2024).

Surface survey catchability is estimated with a vague log-normal prior ($\mu_{q_s} = 0.5$, $\tau_{q_s} = 0.5$), while the dive survey is assumed to be absolute with $q_d = 1$, which matches current operating model assumptions (Johnson et al., 2024). The assumption that the dive survey measures absolute spawning biomass (on average) is appropriate given the coverage of the survey, although there is some debate on the impact of egg survival and hatching rates on estimated total spawn. Nevertheless, a fixed assumption (or informative prior) is needed for spawn survey catchability because the highly variable nature of herring abundance, combined with high observation uncertainty and recently low fishery removals, make it difficult to discern the size of populations from herring dive survey data alone.

#### *A.3. Age composition observation models*

Proportion-at-age (i.e., age composition) observations are modeled via a logistic-normal likelihood function (Schnute and Haigh, 2007; Francis, 2014), with expected values calculated as proportions of the catch-at-age (Table A.3, O.2). Annual age data samples are weighted relative to the average annual sample size for each fleet (Table A.3, A.4), and gear-specific lag-1 auto-correlation matrices are estimated for age composition residuals (Table A.3, A.1). To avoid zeroes in the age composition data, we applied a tail-compression procedure (Francis, 2014) to combine data from age classes with less than 2% of the samples with neighbouring age classes that are above that threshold, creating a variable number of bins at each time step (Table A.3, A.5). Gear and area specific age sampling error standard deviations are considered nuisance parameters and estimated as conditional maximum likelihood estimates (Table A.3, A.7).

#### *A.4. Objective function and optimisation*

The SCAH objective function is proportional to the negative log posterior density function (Table A.3, F.1), and defined as the sum of the observation data negative log likelihood function values (L.6, A.8, and Co.2), negative log prior densities for process errors (P.1 - P.3), shrinkage prior negative log densities (P.4 and P.5), and priors on other leading parameters (P.6 - P.8).

The SCAH TMB model objective function is optimised via the nlminb() function in the R statistical package (R Core Team, 2015; Kristensen et al., 2015). We considered the model parameters converged when the maximum gradient component of the likelihood surface had absolute value less than $10^{-2}$, and the Hessian matrix is positive definite. Bayes posterior distributions are then sampled as 4 independent chains of 1000 samples each using Hamiltonian Monte-Carlo (Monnahan and Kristensen, 2018), or No U-turn Sampling. Hamiltonian Monte-Carlo differs from Markov-Chain Monte-Carlo by minimising the auto-correlation between successive posterior samples, thereby producing a mixed model posterior sample with lower absolute sample sizes and little or no thinning (Monnahan et al., 2017). Chains are considered converged when $\widehat{R} < 1.01$ and effective sample sizes are greater than 400 (i.e., 100 samples per chain) for all parameters.

#### A.5. SCAH model tables

Table A.1: Notation used in the SISCAH OM.

| Symbol | Value | Description |
|---|---|---|
| $T$ | 72 | Total number of time steps 1951 - 2022 |
| $A$ | 10 | Plus group age-class |
| $t$ | $1, 2, \ldots, T$ | Time step |
| $a$ | $1, 2, \ldots, A$ | Age-class index |
| $g$ | $1, 2, \ldots, 6$ | Gear index for (1) reduction, (2) seine-roe, and (3) gillnet fisheries, (4) Surface and (5) dive spawn survey indices, and (6) the closed pond SOK fishery. |
| $B_0$ | | Unfished spawning stock biomass |
| $h$ | | Beverton-Holt stock-recruitment steepness |
| $R_0$ | | Unfished equilibrium recruitment |
| $R_{init}$ | | Initial recruitment parameter |
| $\bar{R}$ | | Average recruitment |
| $\omega_a^{init}$ | | Initial recruitment log-deviations |
| $S_a$ | | Unfished equilibrium survivorship-at-age |
| $\beta_1, \beta_2$ | $10, 4.92$ | Beta prior parameters for stock-recruit steepness |
| $\phi_0$ | | Unfished equilibrium spawning biomass per recruit |
| $\omega_t$ | | Annual recruitment process error log-deviations |
| $\sigma_R$ | 0.8 | Standard error of $\omega_t$ recruitment deviations |
| $q_s$ | | Catchability coefficient for the surface survey design |
| $q_d$ | 1 | Catchability coefficient for the dive survey design |
| $q_t'$ | | Annual convex-combination of survey catchabilities |
| $q_p^{pred}$ | | Per-capita predation rate for predator $p$ |
| $\sigma^{pred}$ | 0.5 | Standard deviation for simple random walk in log per-capita predation rate |
| $\epsilon_{p,t}^{pred}$ | | Standard normal residuals for random walk in log per-capita predation rate |
| $\eta_t$ | | Proportion of annual spawn survey index that is observed by the surface survey design |
| $\mu_{q_s}$ | 0.5 | Prior mean catchability on the log-scale for surface survey |
| $\tau_{q_s}$ | 0.5 | Log-normal prior standard deviation on catchability coefficient for surface survey |
| $M_{t_0}$ | | Initial natural mortality rate |

Table A.1: Notation used in the SISCAH OM. *(continued)*

| Symbol | Value | Description |
|---|---|---|
| $\epsilon_t$ | | Annual log-deviations in natural mortality rate |
| $\sigma_M$ | .1 | Standard error in annual natural mortality log-deviations |
| $M_{a,t}$ | | Annual natural mortality rate for age $a$ |
| $w_{a,t}$ | | Annual average weight-at-age observations |
| $s_g^{50}$ | | Age-at-50% selectivity for gear $g$ |
| $s_g^{Step}$ | | Difference between age-at-50% and age-at-95% selectivity for Gear $g$ |
| $s_{a,g,t}$ | | Selectivity-at-age $a$ for gear $g$ in year $t$ |
| $\delta_g$ | | Fractional time-step at which catch from gear type $g$ is removed from the population |
| $R_t$ | | Estimated Age-1 recruitment (with process error) at time $t$ |
| $\hat{R}_t$ | | Expected age-1 recruitment (deterministic) based on a Beverton-Holt stock-recruit relationship |
| $N_{a,t+\delta_g}$ | | Total numbers-at-age $a$ in area $p$ in year $t$ at fractional time-step $\delta_g$ |
| $N_{a,g,t+\delta_g}$ | | Total numbers-at-age $a$ in area $p$ vulnerable to gear $g$ in year $t$ at fractional time-step $\delta_g$ |
| $B_t$ | | Spawning biomass in year $t$ |
| $C_{a,g,t}$ | | Expected catch-at-age $a$ in numbers by gear $g$ in year $t$ |
| $C'_{a,g,t}$ | | Expected catch-at-age $a$ in biomass units by gear $g$ in year $t$ |
| $\tau_p^C$ | | log-scaled standard deviations from residuals between MLE estimates of consumption for predator $p$ and the estimated consumption from bioenergetic models |
| $U_{g,t}$ | | Harvest rate of population by gear $g$ in year $t$ |
| $K_t$ | | Total landed weight of Spawn-on-Kelp product at time $t$ |
| $P_t$ | | Biomass of fish ponded for Spawn-on-Kelp fishery at time $t$ |
| $\Psi_t$ | | Conversion factor from ponded fish to landed SOK |
| $\mathfrak{F}$ | 200 | Average fecundity of mature, female herring (eggs per gram of spawning biomass) |
| $p_{m,t}$ | | Proportion of individuals that are mature at time $t$ |
| $p_{f,t}$ | 0.5 | Proportion of individuals that are female at time $t$ |
| $p_{eff,t}$ | 0.35 | Proportion of individuals at time $t$ that will spawn in the closed pond |

Table A.1: Notation used in the SISCAH OM. *(continued)*

| Symbol | Value | Description |
|---|---|---|
| $\gamma_g$ | $0.0024, 0.0023$ | Conversion factor from eggs to SOK product |
| $I_{g,t}$ | | Observed spawn index for survey $g \in \{4, 5\}$ at time $t$ |
| $\hat{I}_{g,t}$ | | Expected spawn index for survey $g \in \{4, 5\}$ at time $t$ |
| $\tau^I_t$ | | Standard deviation of blended spawn index observation log-residuals |
| $u_{a,g,t}$ | | Observed composition data for age $a$ by gear $g$ at time $t$ |
| $\hat{u}_{a,g,t}$ | | Expected composition data for age $a$ by gear $g$ at time $t$ |
| $\mathfrak{B}_{g,t}$ | | Total number of age classes with age observations above 2% of the total sample size in year $t$ |
| $\tau^{age}_g$ | | Conditional MLE of age composition sampling error |
| $m_M$ | 0.45 | Prior mean initial natural mortality |
| $s_M$ | 0.4 | log-scale standard deviation on initial mortality |
| $m^{s^{50}}_g$ | | Population mean age-at-50% selectivity for gear $g$ |
| $s^{s^{50}}_g$ | 0.3 | log-deviation of $s^{50}_p$ from $\mu_{s^{50},g}$ |
| $m^{s^{Step}}_g$ | | Population mean step from age-at-50% to age-at-95% selectivity for gear $g$ |
| $s^{s^{Step}}_g$ | 0.3 | log-deviation of $s^{Step}_g$ from $\mu_{s^{Step},g}$ |
| $\rho_g$ | | Correlation-at-lag-1 coefficient for age composition residuals |
| $\mathfrak{C}$ | | Lag-1 correlation matrix for age composition residuals |
| $\mathfrak{K}$ | | Dimension transformation matrix for age composition logistic normal likelihood |
| $\iota_{g,t}$ | | Centred logistic normal age-composition log-residuals for gear $g$ at time step $t$ |

Table A.2: Process model equations for the SISCAH OM.

| No. | Equation |
|---|---|
| **Model Parameters** | |
| (P.1) | $\Theta^{lead} = \left(B_0, \bar{R}, R_{init}, \{\omega_t\}_{t\in 1:T}, M_0, \{\epsilon_t\}_{t\in 1:T}, s^{(50)}, s^{(step)}, \{\psi_g\}_{g\in 1:3}, \{\tau_g\}_{g\in\{4,5\}}, h\right)$ |
| (P.2) | $\Theta^{cond} = \left(\{\log q_g\}_{g\in\{4,5\}}, \{\tau_g^{age}\}_{g\in 1:3}\right)$ |
| (P.3) | $\Theta^{fixed} = (\{m_a\}_{a\in 1:10}, \sigma_R, \sigma_M)$ |
| (P.4) | $\Theta^{priors} = \left(m_M, s_M, \{m_g^{s^{50}}, s_g^{s^{50}}, m_g^{s^{Step}}, s_g^{s^{Step}}, \sigma_g^{Sel}\}_{g\in 1:3}, \beta_1, \beta_2\right)$ |
| **Unfished Equilibrium States** | |
| (EQ.1) | $m_a = \left(1 + e^{-\log 19 \frac{a - a_{50}^{mat}}{a_{95}^{mat} - a_{50}^{mat}}}\right)^{-1}$ |
| (EQ.2) | $S_a = \begin{cases} 1 & a = 1, \\ S_{a-1} e^{-\bar{M}_{a-1}} & 1 < a < A \\ S'_{a-1,g} e^{-\bar{M}_{a-1}} / (1 - e^{-\bar{M}_A}) & a = A. \end{cases}$ |
| (EQ.3) | $\phi = e^{-\bar{M}} \cdot \sum_a S_a \cdot \bar{w}_a \cdot m_a$ |
| (EQ.4) | $R_0 = B_0 / \phi$ |
| (EQ.5) | $N_a^{eq} = R_0 \cdot S_a$ |
| **Non-equilibrium Initial States** | |
| (NEQ.1) | $Z_{init,a} = \bar{M}_a + s_{a,g} * F_{init}$ |
| (NEQ.2) | $S'_a = \begin{cases} 1 & a = 1, \\ S'_{a-1} e^{-Z_{a-1}} & 1 < a < A \\ S'_{A-1,g} e^{-Z_{A-1}} / (1 - e^{-Z_A}) & a = A. \end{cases}$ s |
| (NEQ.3) | $N_{a,1} = R_{init} \cdot S'_a \cdot e^{\omega_a^{init}}$ |
| **Catch from discrete fisheries** | |
| (C.1) | $N_{a,t+\delta_g^-} = N_{a,t+\delta_{g-1}} \cdot e^{-1\cdot(\delta_g - \delta_{g-1})M_t}$ |
| (C.2) | $s_{a,g} = \left(1 + e^{-\log 19 \frac{a - a_{50}^{sel}}{a_{95}^{sel} - a_{50}^{sel}}}\right)^{-1}$ |
| (C.3) | $N_{a,g,t} = N_{a,t+\delta_g^-} \cdot s_{a,g}$ |
| (C.4) | $B_{a,g,t} = N_{a,g,t} \cdot w_{a,t}$ |
| (C.5) | $B_{g,t} = \sum_a B_{a,g,t}$ |
| (C.6) | $C'_{a,g,t} = C_{g,t} \cdot \frac{B_{a,g,t}}{\sum_{a'} B_{a',g,t}}$ |

Table A.2: Process model equations for the SISCAH OM. *(continued)*

| No. | Equation |
|---|---|
| (C.7) | $C_{a,g,t} = C'_{a,g,t}/w_{a,g,t}$ |
| (C.8) | $N_{a,t+\delta_g} = e^{-(\delta_g-\delta_{g-1})\cdot M_{a,t}} \cdot N_{a,t+\delta_{g-1}} - C_{a,g,t}$ |
| (C.9) | $U_{g,t} = C_{g,t}/B_{g,t}$ |
| **Fertilised Herring egg removals** | |
| (K.1) | $\Psi_t = \mathfrak{F} \cdot p_{m,t} \cdot p_{f,t} \cdot p_{eff,t} \cdot \gamma$ |
| (K.2) | $P_t = \hat{K}_t/\Psi_t$ |
| (K.3) | $P_{a,t} = P_t \cdot \frac{B_{a,g,t}}{\sum_{a'} B_{a',g,t}}$ |
| (K.4) | $N_{a,t+\delta_{SOK}} = e^{-(\delta_{SOK}-\delta_{g-1})\cdot M_t} \cdot N_{a,t+\delta_{g-1}} - P_{a,t}/w_{a,t}$ |
| (K.5) | $P'_{a,t} = P_{a,t} \cdot e^{-M_{SOK}}$ |
| (K.6) | $U^{SOK}_t = P_{a,t} \cdot (1 - e^{-M_{SOK}})/B_{SOK,t}$ |
| **Spawning Biomass, Recruitment, and Numbers-at-age** | |
| (A.1) | $B_t = e^{-(1-\delta_G)M_t} \cdot \sum_a N_{a,t+\delta_G} \cdot w_{a,t} \cdot m_a$ |
| (A.2) | $R_{t+1} = \bar{R} \cdot e^{\omega_{R,t}}$ |
| (A.3) | $\hat{R}_{t+1} = \frac{4 \cdot R_0 \cdot B_t}{B_0 \cdot (1-h) + (5h-1) \cdot SB_t}$ |
| (A.4) | $N_{a,t+1} = \begin{cases} R_{t+1} & a = 1 \\ e^{-(1-\delta_G)M_t} \cdot N_{a-1,t+\delta_G} + P'_{a-1,t} & 2 \leq a \leq A-1 \\ e^{-(1-\delta_G)M_t} \cdot (N_{a-1,t+\delta_G} + N_{a,t+\delta_G}) + (P'_{a-1,t} + P'_{a,t-1})e^{-M_{SOK}} & a = A. \end{cases}$ |

Table A.3: Data likelihoods, model prior density functions, and the final objective function for the SISCAH OM. $\mathbf{1}(X)$ is the indicator function that takes value 1 when the statement $X$ is true, and $0$ otherwise.

| No. | Equation |
|---|---|
| **Observation model** | |
| (O.1) | $\hat{I}_t = q'_t \cdot B_t \cdot \mathcal{P}_t$ |
| (O.2) | $\hat{u}_{a,g,t} = \dfrac{C_{a,g,t}}{\sum_{a'} C_{a',g,t}}$ |
| **Spawn index data likelihood** | |
| (L.1) | $z_{1,t} = \log \dfrac{I_t}{\hat{I}_t}$ |
| (L.2) | $q'_t = \eta_t q_4 + (1 - \eta_t) q_5$ |
| (L.3) | $n = \sum_{t=t_0}^{T} \mathbf{1}(I_t > 0)$ |
| (L.4) | $Z = \dfrac{1}{2\tau^2} \sum_{t=t_0}^{T} \mathbf{1}(I_t > 0) z_{1,t}^2$ |
| (L.5) | $l_1 = \dfrac{1}{2}\left(n \cdot \log \tau^2 + Z\right)$ |
| **Age composition data likelihood** | |
| (A.1) | $\mathfrak{C}_g = \left[\rho_g^{j-i}\right]_{i,j}$ |
| (A.2) | $\mathfrak{K} = [I_{A-1} \vert 1]$ |
| (A.3) | $V_g = \mathfrak{K} \cdot C_g \cdot \mathfrak{K}^T$ |
| (A.4) | $W_{g,t} = \dfrac{\text{mean}(n_{g,t}^{Age})}{n_{g,t}^{Age}}$ |
| (A.5) | $\mathfrak{B}_{g,t} = \sum_{a=1}^{A} \mathbf{1}(u_{a,g,t} > 0.02)$ |
| (A.6) | $\vec{\iota}_{g,t} = \langle log(u_{a,g,t}/u_{\mathfrak{B}_{g,t},g,t}) - log(\hat{u}_{a,g,t}/\hat{u}_{\mathfrak{B}_{g,t},g,t} \rangle_a$ |
| (A.7) | $\hat{\tau}_g^{age} = \left(\dfrac{\sum_t \frac{1}{W_{g,t}^2} (\vec{\iota}_{g,t})^T \cdot V_g^{-1} \cdot \vec{\iota}_{g,t}}{\sum_t \mathfrak{B}_{g,t}}\right)^{0.5}$ |
| (A.8) | $l_2 = \sum_g \left(\sum_{a,t} \log u_{a,g,t} + \log \hat{\tau}_g^{age} \sum_t (\mathfrak{B}_{g,t} - 1) + \dfrac{1}{2} \sum_t \log \vert V_g \vert + \sum_t (\mathfrak{B}_{g,t} - 1) \log W_{g,t}\right)$ |
| **Predator consumption likelihood** | |
| (Co.1) | $z_{3,p,t} = \dfrac{\log \hat{C}_{p,t} - \log C_{p,t}}{\tau_p^C}$ |

Table A.3: Data likelihoods, model prior density functions, and the final objective function for the SISCAH OM. $\mathbf{1}(X)$ is the indicator function that takes value 1 when the statement $X$ is true, and $0$ otherwise. *(continued)*

| No. | Equation |
|---|---|
| (Co.2) | $l_3 = \sum_{p,t} (\log \frac{1}{\sqrt{2\pi}} - \frac{z^2_{3,p,t}}{2})$ |
| **Process error prior density functions** | |
| (Pr.1) | $p_1 = \sum_t (\log \frac{1}{\sqrt{2\pi}} - \frac{\omega_t^2}{2})$ |
| (Pr.2) | $p_2 = \sum_a \left( \log \frac{1}{\sqrt{2\pi}} - \frac{\left(\omega_a^{init}\right)^2}{2} \right)$ |
| (Pr.3) | $p_3 = \sum_t (\frac{1}{\sqrt{2\pi}} - \frac{\epsilon_t^2}{2})$ |
| **Priors on other parameters** | |
| (Pr.4) | $p_4 : M_{1951} \sim \log N(0.45, 0.4)$ |
| (Pr.5) | $p_5 : h \sim \beta(20, 9.84)$ |
| (Pr.6) | $p_6 : s_g^{50} \sim \log N(m_g^{s^{50}}, s_g^{s^{50}})$ |
| (Pr.7) | $p_7 : s_g^{Step} \sim \log N(m_g^{s^{Step}}, s_g^{s^{Step}})$ |
| (Pr.8) | $p_8 : q_s \sim \log N(0.5, 0.5)$ |
| (Pr.9) | $p_9 : \tau_g^2 \sim IG(2, 0.27)$ |
| (Pr.10) | $p_{10} : \log q_{p,init}^{pred} \sim N(-4, -3)$ |
| (Pr.11) | $p_{11} : \epsilon_{p,t} \sim N(0, 1)$ |
| (Pr.12) | $p_{12} : \psi_t \sim \log N(0.06, 0.1)$ |
| (Pr.13) | $p_{13} = \sum_{g=1}^{3} \rho_g^2$ |
| (Pr.14) | $p_{14} = \sum_t (\log \frac{1}{\sqrt{2\pi}} - \frac{\left(\log R_t - \log \hat{R}_t\right)^2}{2\sigma_R^2})$ |
| (Pr.15) | $p_{15} : -\log B_0$ |
| (Pr.16) | $p_{16} = -\log R_{bar}$ |
| (Pr.17) | $p_{17} = -\log R_{init}$ |
| **Objective function (negative posterior density)** | |
| (F.1) | $f_{noPredM} = l_1 + l_2 + l_3 + \sum_{j=1}^{16} p_j$ |

Table A.3: Data likelihoods, model prior density functions, and the final objective function for the SISCAH OM. $\mathbf{1}(X)$ is the indicator function that takes value 1 when the statement $X$ is true, and $0$ otherwise. *(continued)*

| No. | Equation |
|---|---|
| (F.2) | $f_{predM} = l_1 + l_2 + l_3 + \sum_{j=1}^{17} p_j$ |

### A.6. SCAH model figures

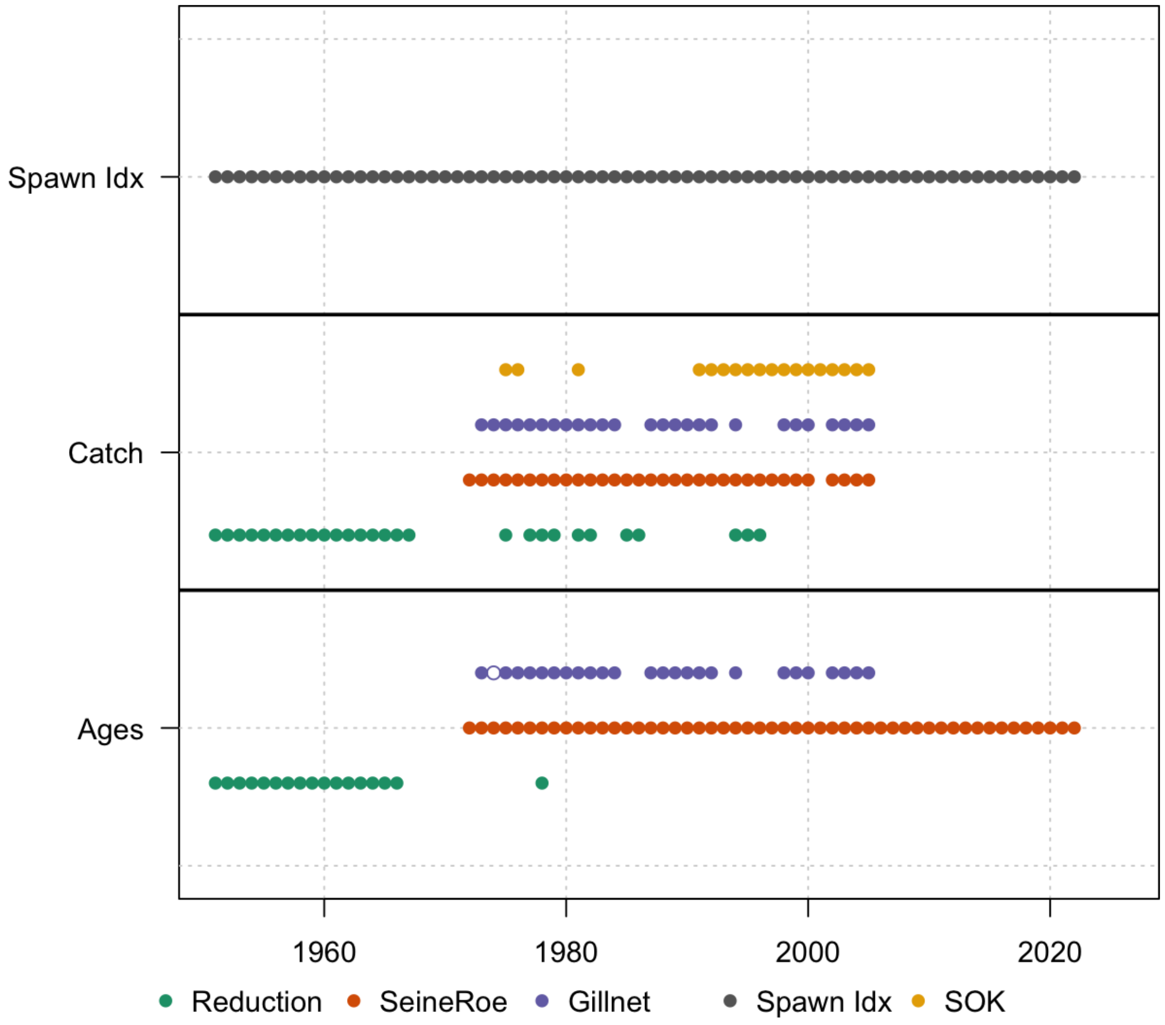


Figure A.1: A summary of data available for fitting the SCAH *noPredM* model. Panels show, from top to bottom, years with spawn survey index data, catch data, and age-composition data for each fishery. Circles in a cell indicate that data of that type exists in that year from the indicated fleet (colour). Open circles for ages indicate years with less than 200 samples.

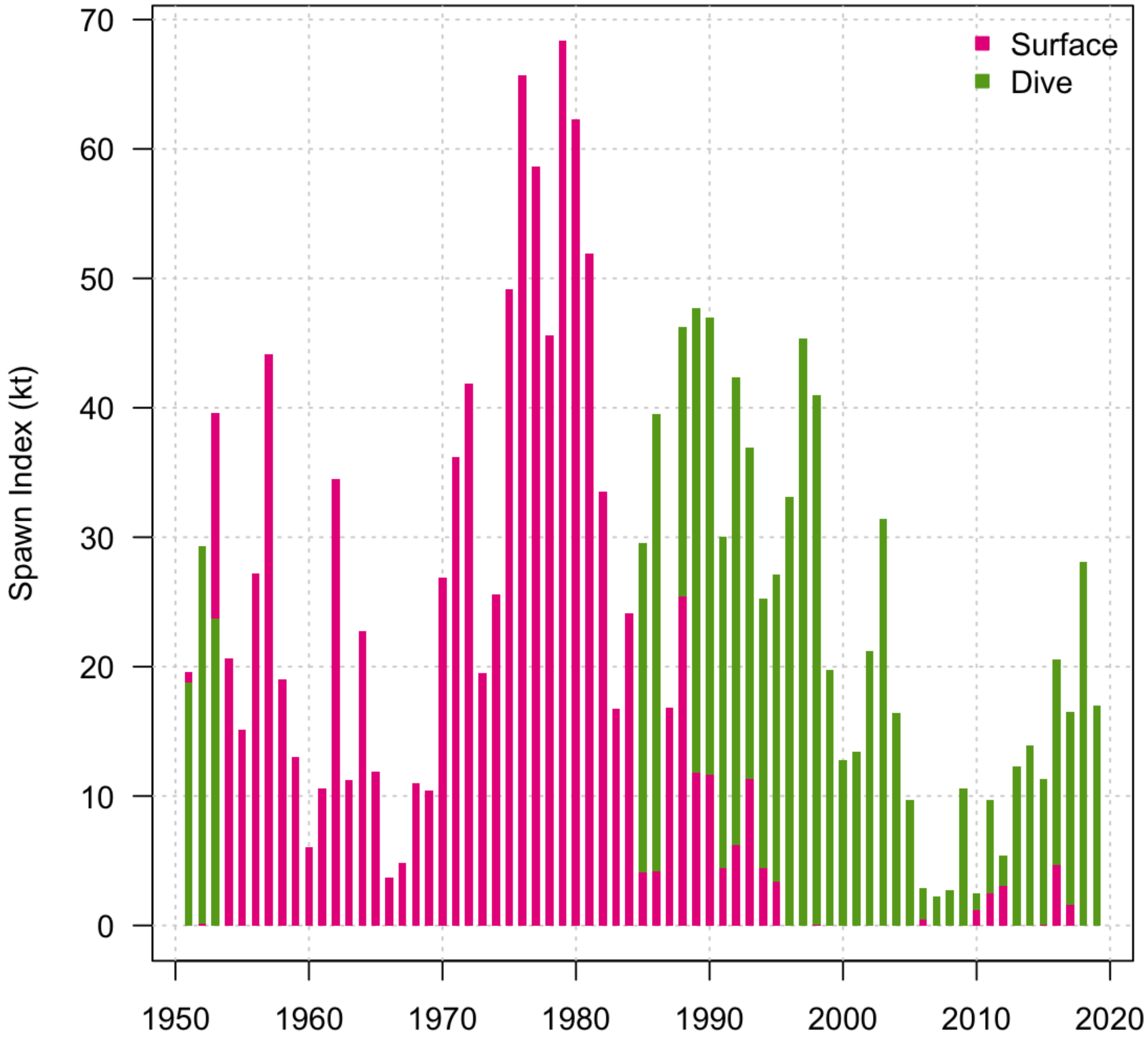


Figure A.2: Spawn survey index contributions from the Surface and Dive survey designs for each sub-area.

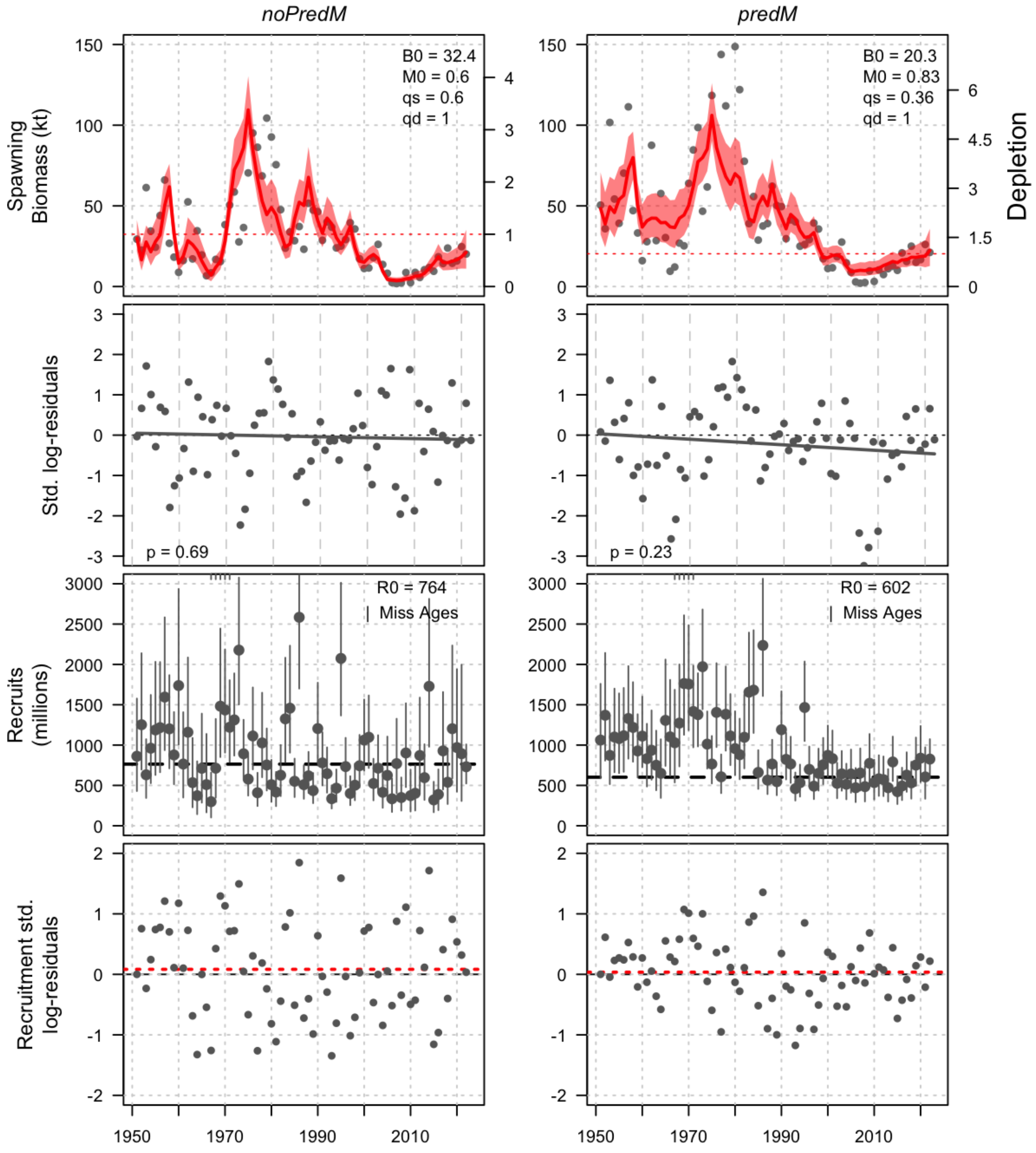


Figure A.3: Comparison of SISCAH model Bayes posterior estimates across the *noPredM* (left) and *predM* (right) hypotheses. Panels show: spawning stock biomass with scaled blended spawn indices, as well as posterior median estimates of unfished biomass $SB_0$, mean (basal) mortality $M_0$, and surface survey catchability $q_s$ (1st row); standardised blended spawn index log-residuals (2nd row), age-1 recruitments (3rd row), and standardised annual age-1 recruitment deviations (points) and the mean deviation (red dashed line) log-scale recruitment process errors (last row).

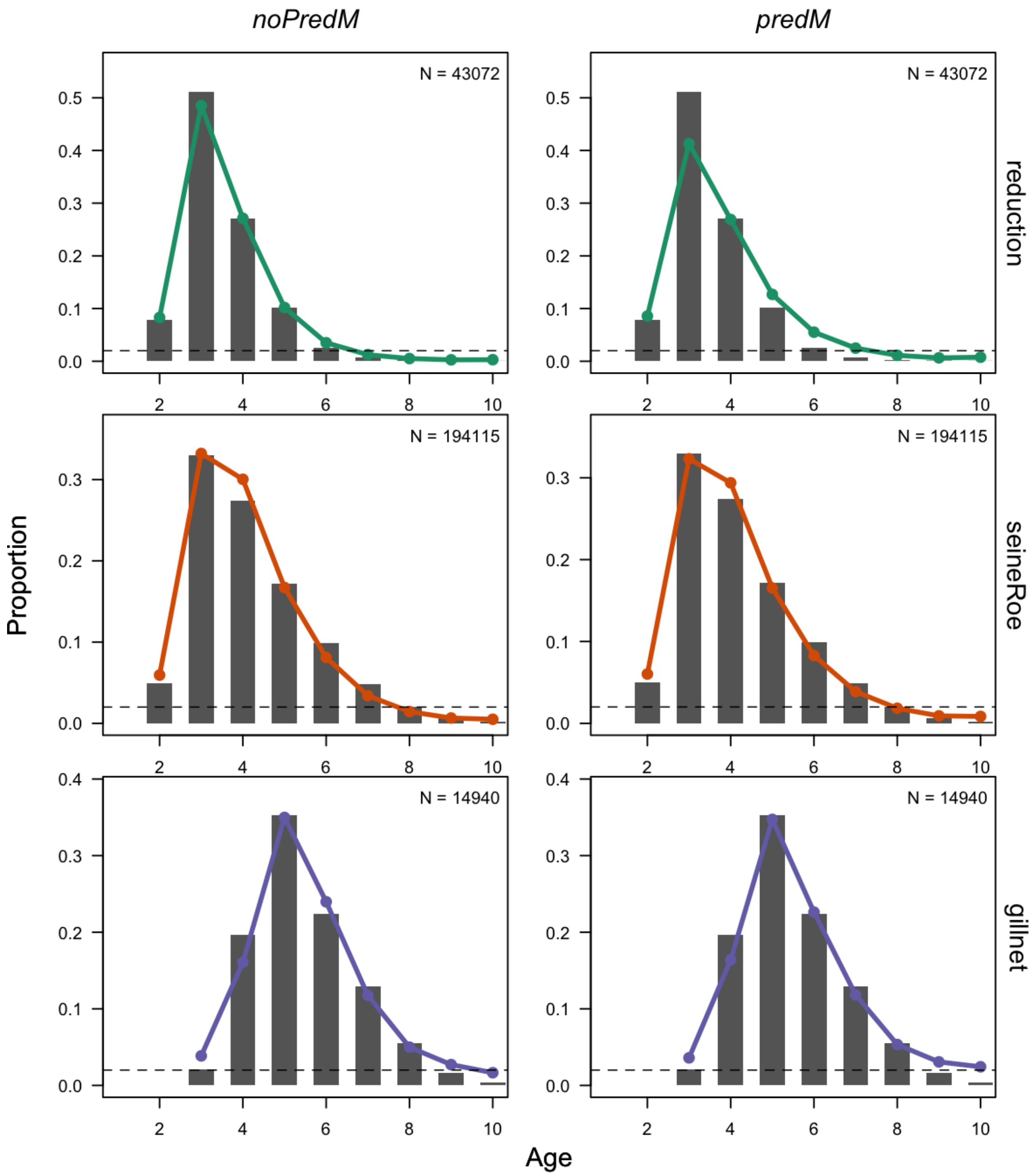


Figure A.4: Time-averaged age composition fits from *noPredM* (left) and *predM* (right) models for reduction, seine-roe, and gilllnet fisheries. Total age samples across all years are shown in top right.

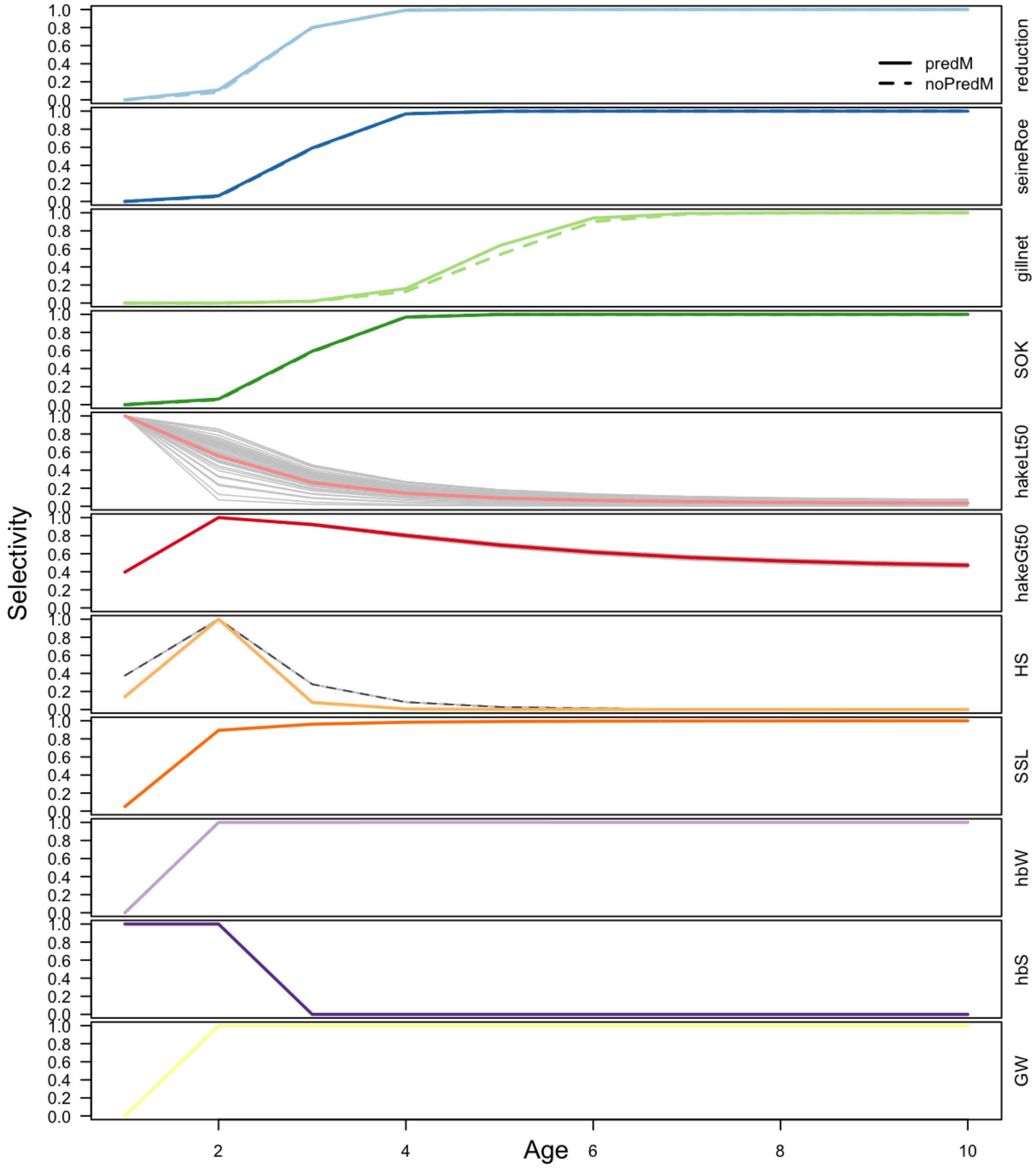


Figure A.5: Selectivity-at-age for each fishery and predator included in *noPredM* (dashed lines) and *predM* (solid lines) models. Thin grey lines indicate time-varying selectivity for the small hake size class *hakeLt50*.

### Supplementary Material B: Modelling predator abundance

#### *B.7. Generalized theta-logistic population model*

Humpback Whale *(Megaptera novaeanglia)*, Steller Sea Lion *(Eumetopias jubatus)*, and Harbour Seal *(Phoca vitulina)* populations that feed on WCVI Pacific Herring were all modeled using the deterministic Bayesian generalized $\theta$-logistic growth model of the form

$$N_{t+1} = N_t + rN_t\left(1 - \left(\frac{N_t}{K}\right)^{\theta}\right) - C_t \tag{3}$$

where $N_t$ is the number of individuals at the start of year $t$, $C_t$ is the annual numbers killed during year $t$, $r$ is the intrinsic growth rate of the population, $K$ is the population carrying capacity, and $\theta$ is the shape parameter that determines the population size at maximum net productivity level (MNPL). Model notations and equations are provided in Table B.1 and Table B.2, respectively. A standard logistic model assumes that MNPL occurs at K/2, whereas the shape parameter $\theta$ on the generalized logistic model controls skew in the production relationship, producing MNPL above K/2 when $\theta > 1$ and below $K/2$ when $\theta < 1$. For all populations, parameters were unable to be estimated assuming an unexploited state, so initial numbers were estimated as a scalar multiple $\kappa$ of carrying capacity.

The generalised $\theta$-logistic population model was specified in Template Model Builder (Kristensen et al., 2015), and leading model parameters $r$, $K$, $\theta$, and $\kappa$ were estimated via Hamiltonian Monte-Carlo using the tmbstan package (Monnahan and Kristensen, 2018). Posterior log-density functions were the sum of log-likelihood functions for observational indices, and $\log$-normal prior density functions for leading model parameters $r$, $K$, and $\theta$. The initialization scalar $\kappa$ was freely estimated.

Observational data for the likelihood function calculation were indices of abundance for surveys $g \in \{1, \ldots, G\}$, modeled via a linear observation model

$$I_{t,g} = q_g B_t e^{\xi_{t,g}} \tag{4}$$

where $q_g$ is catchability and $\xi_{t,g}$ $N(0, \tau_g)$ is a normally distributed random observation deviation in year $t$ for survey index $g$, with observation deviation variance $\tau_g^2$. Nuisance parameters $q_g$ and $\tau_g^2$ were derived from model residuals as conditional maximum likelihood estimates (Table B.1), resulting in a concentrated data likelihood (Table B.2).

*B.7.1. Humpback Whales*

The Humpback Whale population was modeled over the whole British Columbia coast, with 13% of the coastwide total assumed to be present in West Coast of Vancouver Island (WCVI) region year round (Nichol et al., 2010). That value is further reduced for each of the feeding seasons, which is scaled in proportion to seasonal abundance survey estimates for the area relative to maximum numbers observed in the fall (McMillan et al., 2022). We assumed that 53% of the WCVI proportion of Humpback Whales are present from October to March, which is based on an average number of 146 whales from Fall/Winter surveys scaled relative to the maximum numbers of 274 whales in the Fall (McMillan et al., 2022). Similarly, we assumed that 35% of the WCVI proportion of Humpback Whales are present in summer months, which is based on survey estimates of 96 whales during summer relative to the Fall maximum of 274 (McMillan et al., 2022).

We used prior hyperparameters $\mu^r = 0.085$ and $\sigma^r = 0.015$ for the intrinsic rate of growth $r$ based on global annual growth rate estimates for Humpback Whales (Mobley et al., 1999, 2001; Mizroch et al., 2004; Calambokidis et al., 2004; Zerbini et al., 2006; Calambokidis et al., 2008; Calambokidis, 2009; Ford et al., 2009; Zerbini et al., 2010; Wedekin et al., 2017). The prior structure assumes that most populations were heavily depleted post-whaling, so that estimates of annual growth rates should be similar to values for $r$ when $N_t/K$ is small. A vague prior with 100% CV was used for $K$, with $\mu^K = \sigma^K = 5,500$ based on the sum of historical catch from 1905-1930 during the initial period of modern shore-based commercial whaling in BC. The parameter $\theta$ had prior hyperparameters $\mu^r = 2.39$ and $\sigma^r = 0.4$, corresponding to an informative prior around a MNPL of $0.6K$, which is typically assumed for whale populations (Baker and Clapham, 2004; Ivashchenko et al., 2016).

Absolute abundance indices were taken from mark-recapture estimates (Chapman-modified Lincoln-Petersen) from 1992-2005 (Ford et al., 2009), where $q_1$ is fixed at 1. Annual observation error standard deviations (SDs) $\tau_t$ were scaled in proportion to annual estimates from survey CVs from abundance estimates to account for reduced uncertainty in more recent years with increased sampling effort (Ford et al., 2009)

$$w_{t,g} = \frac{\tau_t}{\tau} = \frac{\sqrt{CV_t^2 + 1}}{\tau}$$

where $\tau$ is the average observation error SD and $w_{t,g}$ is a scalar for annual observation errors estimated in the model and $CV_t$ is the estimated annual CVs from abundance estimates ranging from 0.54 to 0.06 with increasing precision in more recent years (Table 1, Ford et al. (2009)).

Humpback Whale removals were taken from commercial whaling records from BC for 5 whaling stations in BC from 1908-1965 (Naden Harbour, Rose Harbour, Coal Harbour, Kyuquot, Sechart; (Nichol and Heise, 1992), after which legal whaling operations ceased internationally. While the population model is not fit to data before 1950, to generate the prior mean carrying capacity we assume that 200 whales were killed per year over 1905-1907 (Ford et al., 2009). Humpback catches were adjusted upwards by 2% to account for whales that were struck and lost (Reeves et al., 1985, in Ford et al. 2009).

Posterior estimates of leading generalized logistic model parameters are provided in Table B.3 and abundance estimates are shown in Figure B.1. The mean posterior estimate for BC Humpback Whales in 2005 is 1684 whales (95% CI=1476-1893) and the forward projection for 2022 is 4833 whales (95% CI= 2191-7996), which are 32% (95% CI=0.11-0.9) and 80% (95% CI=0.43-0.995) of carrying capacity $K$, respectively. The lower limits of the 95% posterior distribution indicate that carrying capacity could be reached as early as 2031 around 2200 whales, while upper limits of 15,000 whales indicate carrying capacity will not be reached until 2065.

#### *B.7.2. Steller Sea Lions*

The WCVI population of Stellar Sea Lions (SSL) is explicitly modeled, and therefore, unlike Humpback Whales or Harbour Seals that use coastwide models, no scaling is needed to estimate the proportion of the modelled SSL population feeding in the WCVI herring stock assessment region.

We used a weakly informative prior for growth rate with mean $\mu^r$=0.04 based on previous estimates for BC Steller Sea Lions (Olesiuk, 2018) and standard deviation $\sigma^r$=0.012. A vague prior with 100% CV was defined for $K$, with $\mu^K$=$\sigma^K$=10,000, derived from the sum of historical kills of WCVI adult Stellar Sea Lions during intensive culling programs from 1913-1939. Prior hyperparameters $\mu^\theta$=3.48 and $\sigma^\theta$=2 are based on prior knowledge that MNLP levels for marine mammals range from $0.5K - 0.8K$, corresponding to values for $\theta$ ranging from $1 - 11$ (Ragen, 1995; Read and Wade, 2000; Maunder et al., 2000; Olesiuk, 1999, 2009).

The Steller Sea Lion population was fit to two indices of abundance; i) a relative abundance index of historic counts from Scott Island rookeries (Triangle Island, Sartine Island, Beresford Island, Maggot Island) from 1913-1982 (Biggs 1985), and ii) an absolute abundance (i.e., $q_2$=1) index of summer adult counts from rookeries and haul out sites for WCVI from 1971-2013 (Olesiuk, 2018).

Annual removals of Stellar Sea Lions for research, commercial, and population control programs were compiled for WCVI sites from 1913-1968 (Biggs 1984), after which the species was protected. All index observations for both gears were weighted equally (i.e., $w_{t,g}$= 1).

Posterior estimates of leading generalized logistic model parameters are provided in Table B.3 and abundance estimates are shown in Figure B.2. The mean posterior estimate for WCVI Stellar sea lions in

2013 is 8,300 (95% CI=6,800-9,900) and the forward projection for 2022 is 10,400 (95% CI= 8,400-13,300), which are 56% (95% CI=0.27-0.89) and 70% (95% CI=0.35-0.98) of carrying capacity $K$, respectively. There is large uncertainty in carrying capacity estimates with mean posterior of 15,400 sea lions (95% CI= 9.8 - 29.8). The lower limits of the 95% posterior distribution indicate that 99% of carrying capacity could be reached by 2039 at 9730 Stellar sea lions, while upper limits of 29,700 indicate 99% of carrying capacity won't be reached until 2155 (Figure B.2).

#### *B.7.3. Harbour Seals*

We used a coastwide model for the BC Harbour Seal population that combined census and removal data for all areas inside and outside Strait of Georgia. The number of seals feeding in the WCVI stock assessment region was assumed to be 5.1% of the total BC population based on the 2008 survey numbers for Pacific Fishery Management Areas 23, 24 and 25 (Olesiuk, 2009).

The intrinsic rate of growth $r$ prior hyperparameters $\mu^r$=0.15,$\sigma^r$=0.015 are based on a weighted average of $r$ estimates for two distinct Harbour seal populations in outside waters and in the Strait of Georgia (Olesiuk, 2009). A vague normal prior with 100% CV was used for $K$, with $\mu^K$=$\sigma^K$=105,000, which is the 2008 coastwide abundance estimate for British Columbia (DFO, 2010), chosen because logistic models indicated populations in Strait of Georgia and outer BC were at or near carrying capacity, respectively (Olesiuk, 2009). The prior mean $\mu^\theta$=5.77 and SD $\sigma^\theta$=0.577 is based on the estimate for the Strait of Georgia (Olesiuk, 2009). with an assumed 10% CV.

The Harbour Seal population model was fit to three indices of abundance; i) a relative abundance index of historic counts from Strait of Georgia from 1973-2008 (Fig. 16 in Olesiuk 2010), ii) a relative abundance index of historic counts from areas outside Strait of Georgia from 1976-2008 (Fig. 16 in Olesiuk, 2009), and iii) an absolute abundance index from 1950-1972 based on a backwards model extrapolation (similar to stock reduction analysis) used to reconstruct historical abundance (Fig. 2 in DFO, 2010). All observations were equally weighted (i.e., $w_{t,g}$=1).

Annual removals of Harbour Seals were based on historical data for seal pelts processed, bounties paid for seal pelts, and predator control records (Olesiuk 2009) as

$$C_t = \frac{B_t + P_t}{R} + D_t + P_k \cdot D'_t$$

where $B_t$ is bounties paid for seal pelts, $P_t$ is additional processed seal pelts, $R$ = 0.616 the recovery rate of carcasses, $D_t$ and $D'_t$ are, respectively, the confirmed and probable numbers of dead seals killed by Fisheries and Oceans predator control actions, and $P_k$=0.75 is the assumed death rate of probable kills during predator control (Eq 13 and Table 7 in Olesiuk, 2009).

Posterior estimates of leading generalized logistic model parameters are provided in Table B.3 and abundance estimates are shown in Figure B.3. The mean posterior estimate for BC Harbour Seals in 2008 is 109,000 (95% CI=94,000-126,000) and the forward projection for 2022 is 109,000 (95% CI=94,000-126,000), both of which are at carrying capacity $K$. There is a lower range of uncertainty in carrying capacity estimates for Harbour Seals relative to the Humpback Whales and Stellar Sea Lion models with 95% posterior distributions indicating carrying capacity was reached between 1998-2000 (Figure B.3). Note that census estimates for outside Strait of Georgia (bcCoast, Figure B.3) shows an increasing trend for the last 3 surveys (2005,2007,2008), whereas the Strait of Georgia (SOG) census does not. This might indicate that the SOG population has reached carrying capacity and that the outside population is still increasing, in which case future Harbour Seal modelling might consider separate population models for areas outside SOG and inside SOG (e.g., Olesiuk, 2009).

*B.7.4. Pacific Hake*

Pacific Hake *(Merluccius productus)* abundance on the WCVI was estimated from the outputs of recent stock assessments (Johnson et al., 2021). Hake are a transboundary stock ranging from southern California to Alaska managed jointly by the USA and Canada according to a treaty. As such, stock assessments for Pacific Hake estimate biomass and abundance of females over the entire range of the species (Johnson et al., 2021), so we used spatial data from acoustic surveys to estimate the proportion of male and female biomass-at-age within WCVI. Survey observations of Pacific Hake are collected via a standardised Hake Survey, which spans the range of the stock from California to northern British Columbia. In addition to the acoustic survey, there are non-standardized observations of Hake feeding aggregations along La Perouse bank off the coast of WCVI, which we use for estimating diet related quantities for Hake bio-energetic models.

While the WCVI herring spawn events occur in inshore areas (i.e., PFMAS 23, 24, and 25), the interactions between Pacific Hake and Pacific Herring occur predominatly offshore. Therefore, the spatial boundary used to define observations of Pacific Hake in the WCVI area was defined by aggregating PFMAs 23, 24, 25 and the neighbouring offshore PFMAS 123, 124, and 125 (Map of PFMAs available at https://www.pac.dfo-mpo.gc.ca/fm-gp/maps-cartes/areas-secteurs/index-eng.html)

Male and female biomass at age in WCVI were estimated from coast-wide female biomass-at-age using the following steps

1. Convert yearly coastwide biomass-at-age $B_{a,t}$ to WCVI yearly biomass-at-age $B^{W}_{a,t}$ by estimating the

annual proportion of biomass in each age-class in the WCVI for each year

a. In years with biomass-at-age data from the acoustic survey, use observed proportions $p^W_{a,t}$=$B^W_{a,t}/B_{a,t}$ inside WCVI. Within a year $t$ with data, fill missing ages with the average proportion over years with data for that age
b. For age-1, we set $B^W_{a,t} = 0$, as young of the year will most likely remain in California
c. Post 1995, interpolate missing years within an age-class, as there is no evidence of cohort signal
d. Pre 1995, use average of observed data post 1995 (before filling missing age classes in a)

2. Multiply coastwide biomass at age by the proportions estimated above to get biomass-at-age in WCVI, i.e.

$$\widehat{B^W_{a,t}} = p^W_{a,t} \cdot B_{a,t}$$

3. Estimate the proportion $\rho_{f,a,t}$=$B_{f,a,t}/B_{a,t}$ of female Hake within the WCVI area in each age class and year of acoustic survey data,
   a. Fill missing age classes in years with data with average for that year class over the data.
   b. Given low sample sizes at older ages, ratios for age 12+ were calculated from pooled data.
   c. Post 1995, fill missing years using linear interpolation within an age class across years
   d. Pre 1995, use average from observed values 1995+
4. Multiply total WCVI biomass-at-age by $\rho_{f,a,t}$ and $(1 - \rho_{f,a,t})$ to get WCVI biomass-at-age for males and females.

This procedure produces a time-series of total male and female biomass-at-age for the WCVI area. Total WCVI Hake Biomass is given in Figure B.4 (second panel).

Pacific Hake male and female biomass-at-age were then used to generate two piscivorous Hake sex-aggregated predator length-classes split at 50 cm fork length. The 50 cm break was based on the changes in diets around that length. For Pacific Hake above 50 cm fork length, the proportion of stomach contents made up by Pacific Herring and the proportion of piscivorous Hake (both defined below) both increased quickly (Figures B.5 and B.6). Biomass within each length class is estimated from sex-structured biomass-at-age via age-length keys, which are in turn estimated via fitting Von Bertalanffy growth models to biological data collected from the Hake Acoustic Survey (Figure B.7).

For each length-class $l$ of Hake, the proportion piscivorous $p^{Pisc}_l$ is calculated simply as

$$p^{Pisc}_l = \frac{O^{Herr}_l}{O_l}, \tag{5}$$

where $O_l$ is the total number of hake observed across both surveys in length class $l$, and $O_l^{Herr}$ is number of Hake observed in length class $l$ containing Herring in their stomach contents, found by aggregating over length classes shown in Figure B.6. Similarly, diet proportion for each length class is calculated as

$$p_l^{Diet} = \frac{V_l^{Herr}}{V_l}, \tag{6}$$

where $V_l$ is the total volume of all stomach contents observed for sampled Hake in length class $l$, and $V_l^{Herr}$ is the total volume of Herring found in the stomach contents of the same sampled Hake, again found by aggregating data shown in Figure B.5. For both proportion piscivorous and the proportion of Hake diet made up by herring, the samples are taken from both the Hake Acoustic and La Perouse Bank surveys.

For simulations, we condition a coastwide combined-sex Pacific Hake population using the parameters from the stock assessment (Johnson et al., 2021). A simulated biomass for each Pacific Hake predator size class (piscivorous Hake, below and above 50 cm in fork length) are then derived from the coastwide combined-sex biomass using the process above. Predator size class biomasses are then used as inputs to the predation mortality models described in the main body of this paper.

*B.7.5. Gray Whales*

The population model for Pacific Gray Whales feeding on herring roe in WCVI draws on estimates of the abundance of two Grey Whale populations that migrate through the WCVI area. There are three distinct designable units of Pacific Gray Whales that have been observed migrating through the WCVI area, namely the Pacific Coast Feeding Group (PCFG), the North Pacific Migratory (NPM) population, and the Western Pacific group (WP) (Gavrilchuk and Doniol-Valcroze, 2021). As an additional complication, distinct abundance estimates for each group are only available beginning in 1999 (Gavrilchuk and Doniol-Valcroze, 2021), while prior to 1998 abundances estimates are only available for the Eastern North Pacific (ENP) population, which aggregates the NPM and PCFG, beginning in 1967 (Durban et al., 2015, 2017). To simplify our approach, we assume that (i) only the PCFG whales are feeding on Herring roe in WCVI, and (ii) Gray Whales feeding on the WCVI during Herring spawning aggregations feed exclusively on Herring spawn. Such assumptions are reasonable because PCFG whales are strongly associated with early spring feeding on Pacific Herring roe in Barkley and Clayoquot Sounds (COSEWIC, 2004; Darling et al., 1998; Darling, 1984; Ford et al., 1994, 2013), and a whale that died in the area during herring spawning season was found to have its stomach filled with herring eggs (Darling et al., 1998).

A parametric theta-logistic model of PCFG abundance has already been defined for Grey Whale abundance after 2015 (Gavrilchuk and Doniol-Valcroze, 2021). The model closely matches the general model used above for Harbour Seals, Steller Sea Lions, and Humpback Whales, but includes an extra yearly mortality term $M_t$ for non-anthropogenic sources of mortality, i.e.,

$$N_{t+1} = N_t + R_{max} N_t \left(1 - \left(\frac{N_t}{K}\right)^{\theta}\right) - C_t - M_t, \quad (7)$$

where $N_{t+1}$ is the population abundance, $R_{max}$ is the exponential growth rate at low population size, $K$ is carrying capacity, $\theta$ controls skew (and therefore density dependence) in the logistic production relationship, and $C_t$ is the anthropogenic mortality, largely representing aboriginal harvest (Gavrilchuk and Doniol-Valcroze, 2021). For projections of Grey Whale abundance, parameter uncertainty is simulated by drawing random model parameter values from distributions defined in Gavrilchuk and Doniol-Valcroze (2021).

Before 2015, interpolation is used to estimate PCFG abundance from ENP abundance estimates between 1967 and 1997. After 1997, there are nine years with abundance estimates for both populations, to which we fit a simple linear model to estimate the ratio $r$ of population sizes ($R^2 = 0.84$, (Fig B.10 upper panel)). Then, PCFG population sizes are interpolated as $N_{PCFG} = rN_{ENP}$ in the years where $N_{ENP}$ esitmates exist prior to 1999 (filled points, Fig B.10 lower panel). For years where no ENP estimates exist, existing estimates are used to linearly interpolate missing years using weighted averages (open points, Fig B.10 lower panel). The interpolation method produces a steep decline in whales from around 200 individuals in 1997 to around 135 individuals in 1998. However, this decline roughly coincides with an unusual mortality event for the ENP gray whale population, which left around 634 emaciated gray whales stranded along the coast between Mexico and Alaska in 1999 and 2000 (Gavrilchuk and Doniol-Valcroze, 2021), so we deemed the large decline in abundance acceptable for the purposes of estimating Herring spawn consumption by PCFG Gray Whales. Finally, prior to 1967 the PCFG is assumed to have a constant abundance of 154 individuals.

##### B.7.6. Tables

Table B.1: Notation for the generalized logistic population dynamics model for marine mammals.

| Symbol | Description |
|---|---|
| | **Indices and index ranges** |
| $T$ | Year in which stock assessment is performed |
| $t$ | Year, where $t = t_1, ..., T$ |
| $g$ | Survey index where $g = 1, ..., G$ |
| $n_g$ | Number of non-missing observations for the index $g$ |
| | **Data** |
| $C_t$ | Catch numbers removed during year $t$ |
| $I_{t,g}$ | Stock relative or absolute abundance observation for year $t$ and index $g$ |
| $w_{t,g}$ | Scalar for observation error standard deviations for year $t$ and gear $g$ |
| | **Leading model parameters** |
| $K$ | Carrying capacity |
| $r$ | Intrinsic growth rate |
| $\theta$ | Shape parameter |
| $\kappa$ | Initial population scalar |
| | **Nuisance parameters** |
| $q_g$ | Catchability coefficient for abundance index $g$ |
| $\tau$ | Average observation error standard deviation |
| | **State variables** |
| $N_t$ | Numbers at the beginning of year $t$ |
| | **Prior distributions** |
| $N(\mu^r, \sigma^r)$ | Normal prior on $r$ |

Table B.1: Notation for the generalized logistic population dynamics model for marine mammals. *(continued)*

| Symbol | Description |
|---|---|
| $N(\mu^K, \sigma^K)$ | Normal prior on $K$ |
| $N(\mu^\theta, \sigma^\theta)$ | Normal prior on $\theta$ |
| | **Error distributions** |
| $\xi_{t,g} \sim N(0, \tau^2)$ | Observation error in year $t$ for index $g$ |

Table B.2: Process model equations for the generalized logistic population dynamics model for marine mammals.

| No. | Equation |
|---|---|
| **Model Parameters** | |
| E2.1 | $\Phi = (K', r', \theta', log\kappa)$ |
| **Parameter Transformation** | |
| E2.2 | $K = exp(K')$ |
| E2.3 | $r = exp(r')$ |
| E2.4 | $\theta = exp(\theta')$ |
| E2.5 | $\kappa = exp(\kappa')$ |
| **Population Dynamics Model** | |
| E2.6 | $N_{t_1} = \kappa \cdot K$ <br> $N_{t+1} = N_t + rN_t(1 - (\frac{N_t}{K})^\theta) - C_t, \quad t > t_1$ |
| **Observation Model Residuals** | |
| E2.7 | $\xi_{t,g} = ln(I_{t,g}/N_t)$ |
| **Conditional maximum likelihood estimates** | |
| E2.8 | $n_g = \sum_{t=t_1}^{T} \mathbb{1}(I_{t,g} > 0)$ |
| E2.9 | $\widehat{ln(q_g)} = \frac{1}{n_g} \sum_{t=t_1}^{T} \mathbb{1}(I_{t,g} > 0) \cdot \xi_{t,g}$ |
| E2.10 | $\widehat{\tau_g^2} = \frac{1}{n_g} \sum_{t=t_1}^{T} \mathbb{1}(I_{t,g} > 0) \cdot (\frac{\xi_{t,g} - \widehat{ln(q_g)}}{W_{t,g}})^2$ |
| **Negative log-likelihood and objective function** | |
| E2.11 | $l(\Phi \vert I_{t,g}) = \frac{\sum n_g}{2} ln\tau^2$ |
| E2.12 | $G(\Phi \vert I_{t,g}) \propto l(\Phi \vert I_{t,g}) + \frac{1}{2(\sigma^K)^2}(K - \mu^K) + \frac{1}{2(\sigma^r)^2}(r - \mu^r) + \frac{1}{2(\sigma^\theta)^2}(\theta - \mu^\theta)$ |

Table B.3: Posterior estimates of leading generalized logistic model parameters $K$, $r$, $\theta$ and $\kappa$ for Humpback Whales, Steller Sea Lions, and Harbour seals. Posterior mean estimates are shown for each parameter with 95% credibility intervals. Population carrying capacity are shown in 1000s.

| Parameter | Humpback Whales | Steller Sea Lions | Harbour seals |
|---|---|---|---|
| $K$ | 5.9 (2.2 - 15.0) | 15.4 (9.8 - 29.8) | 109 (94 - 126) |
| $r$ | 0.09 (0.06 - 0.11) | 0.04 (0.03 - 0.05) | 0.12 (0.11 - 0.13) |
| $\theta$ | 2.3 (1.6 - 3.2) | 2.9 (0.8 - 7.0) | 5.2 (4.0 - 6.5) |
| $\kappa$ | 0.07 (0.03 - 0.19) | 0.57 (0.27 - 1.07) | 0.39 (0.32 - 0.46) |

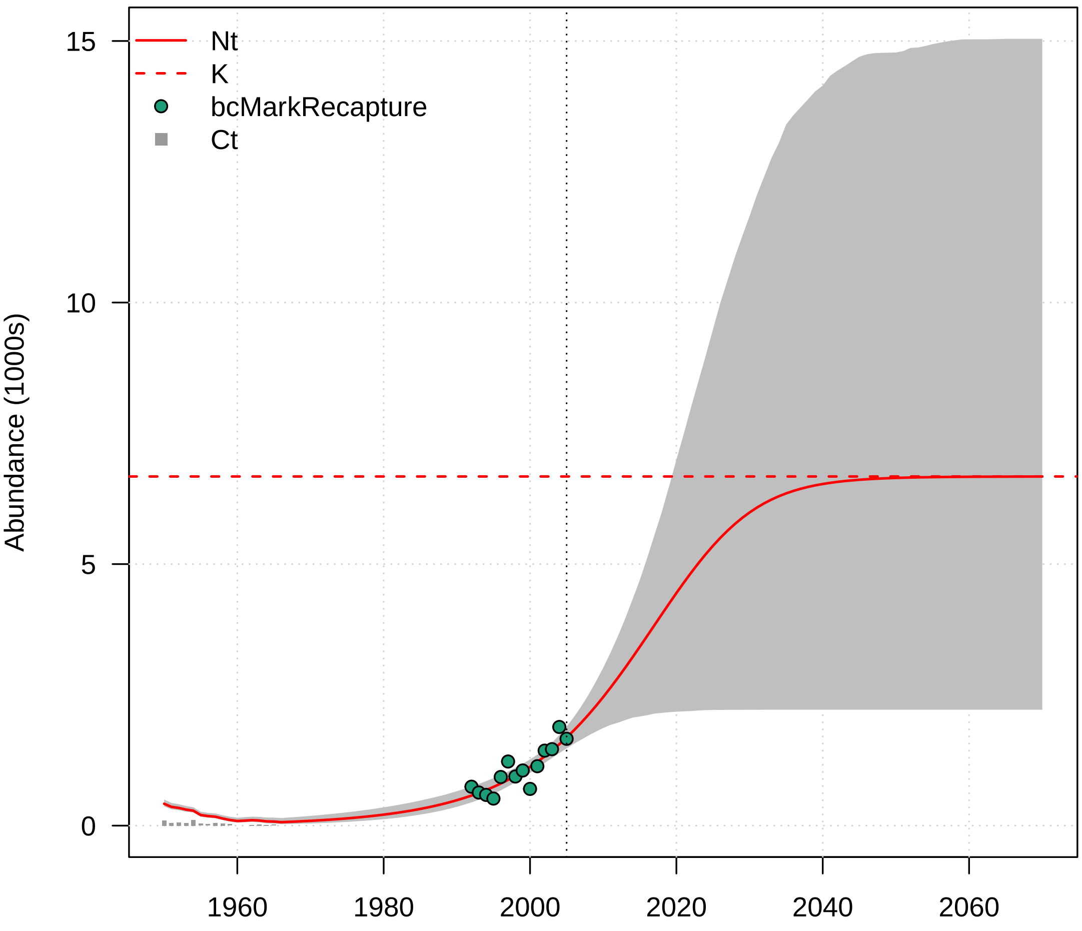


Figure B.1: Mean (red line) and central 95% (grey envelope) of the posterior distribution of Humpback Whale abundance estimates for the BC coast from 1950 until 2005 (vertical dashed line) with projections to 2070 with no removals. Points show abundance indices coloured according to the legend, while grey bars at the bottom of the plot show historical removals.

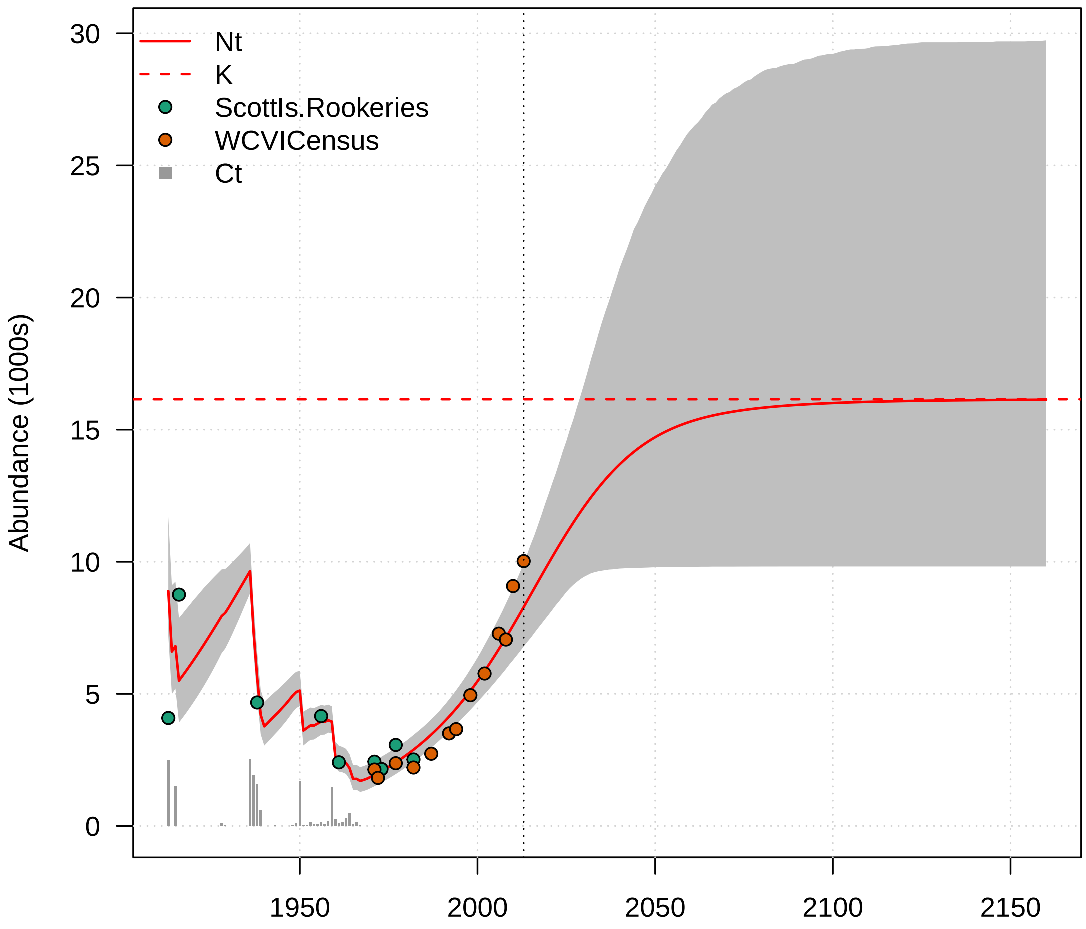


Figure B.2: Mean (red line) and central 95% (grey envelope) of the posterior distribution of Steller Sea Lion abundance estimates on the West Coast of Vancouver Island from 1913 until 2013 (vertical dashed line) with projections to 2160 with no removals. Points show abundance indices coloured according to the legend, while grey bars at the bottom of the plot show historical removals.

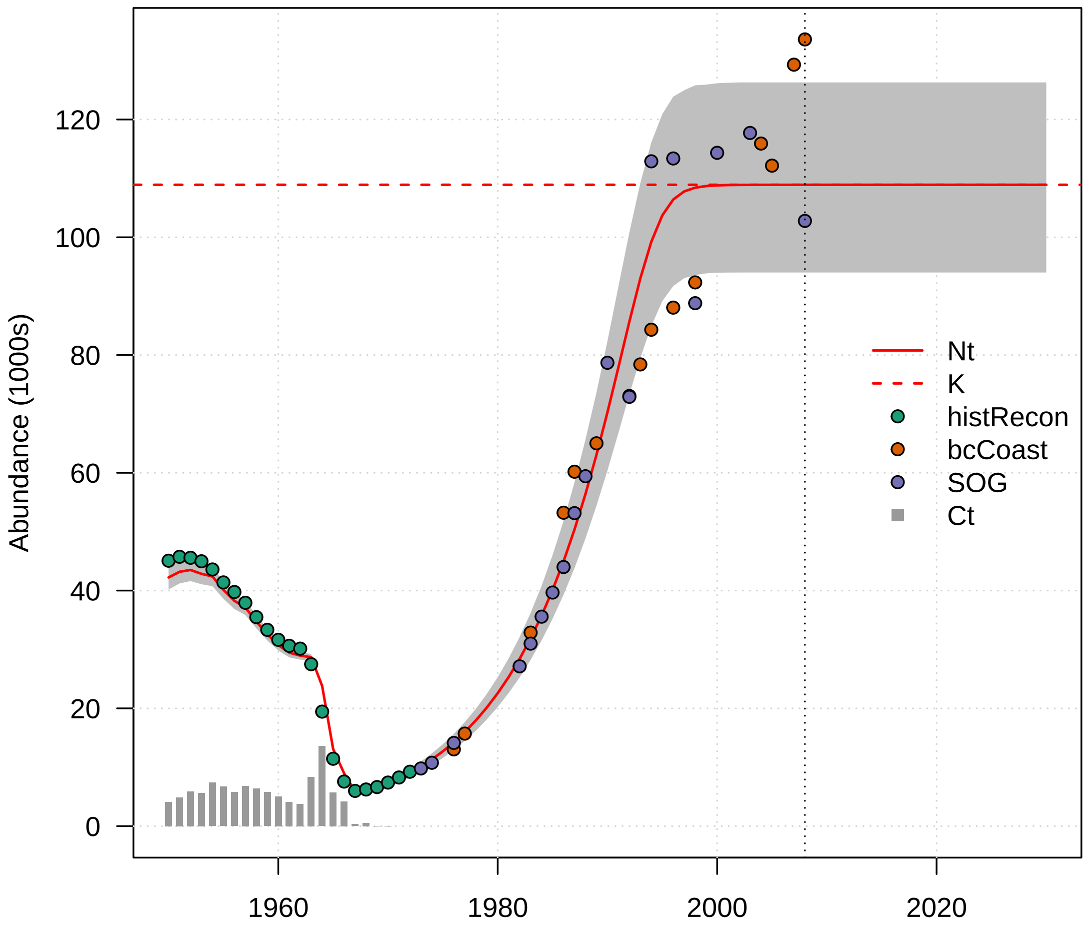


Figure B.3: Mean (red line) and central 95% (grey envelope) of the posterior distribution of Harbour seal abundance estimates for all of BC from 1950 until 2008 (vertical dashed line) with projections until 2030 with no removals. Points show abundance indices coloured according to the legend, while grey bars at the bottom of the plot show historical removals.

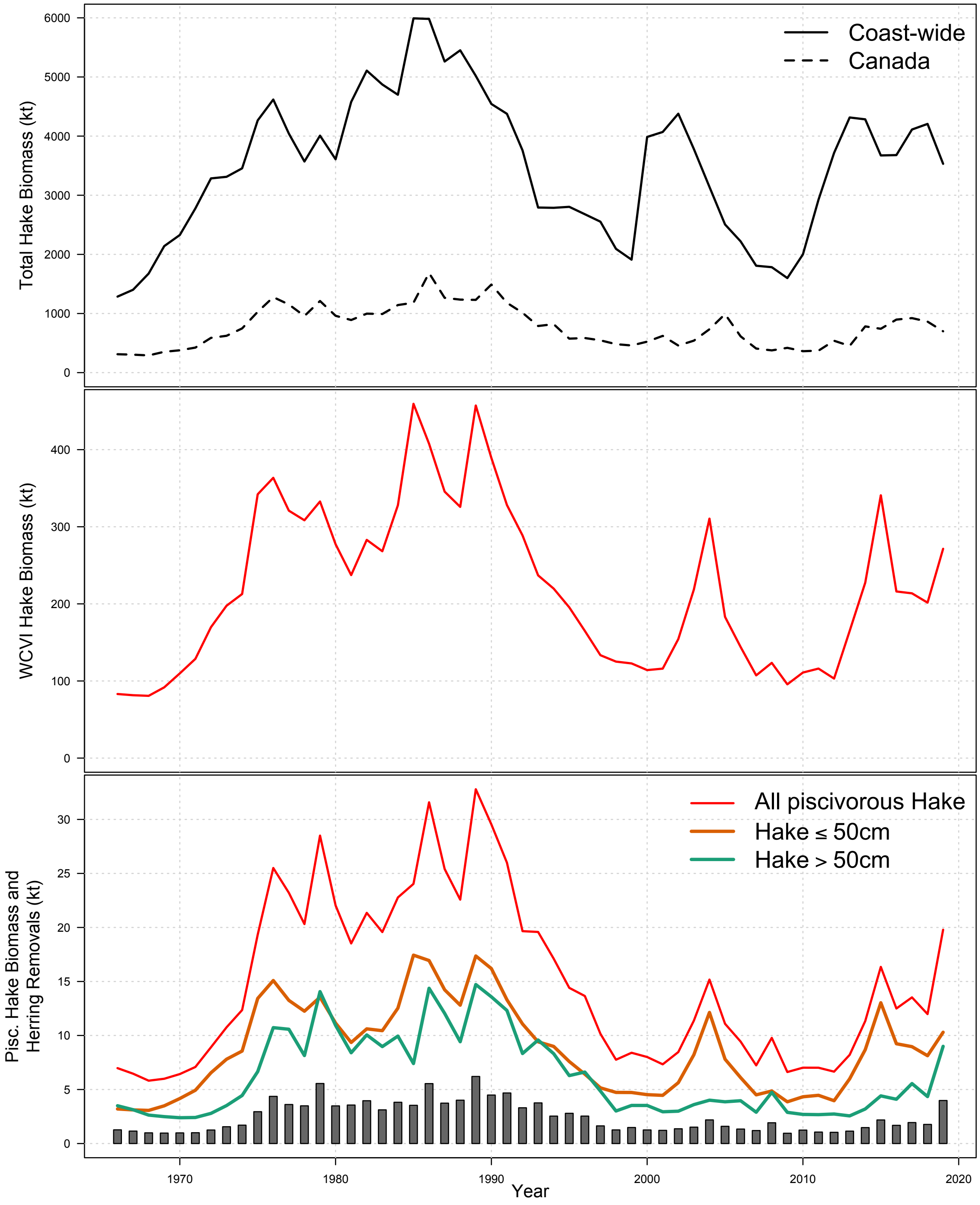


Figure B.4: Estimates for total biomass of Pacific Hake (top), WCVI biomass (middle), and Piscivorous Hake biomass for two size class (Hake ≤ 50 cm and Hake > 50 cm) with total consumption (grey vertical bars) of Herring (bottom).

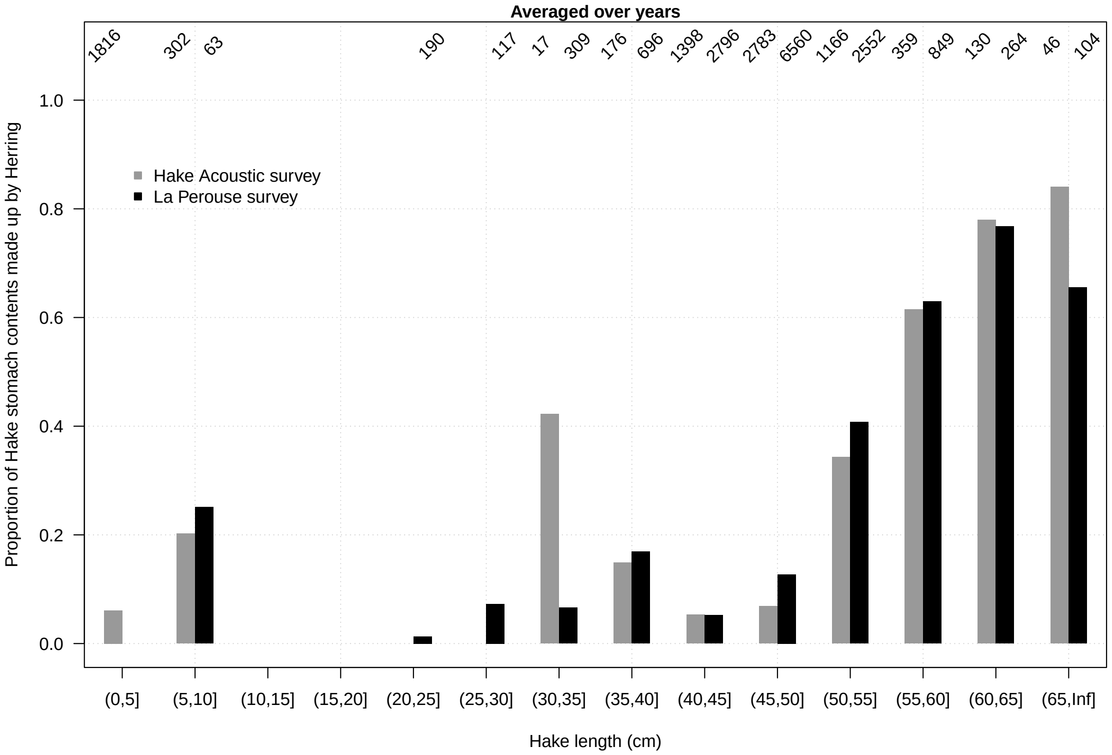


Figure B.5: Proportion of Hake stomach contents (y-axis) made up by Herring, broken up into 5cm Hake length bins (x-axis) with all Hake above 65 cm combined. Data are split into Hake Acoustic and La Perouse surveys, and aggregated over years.

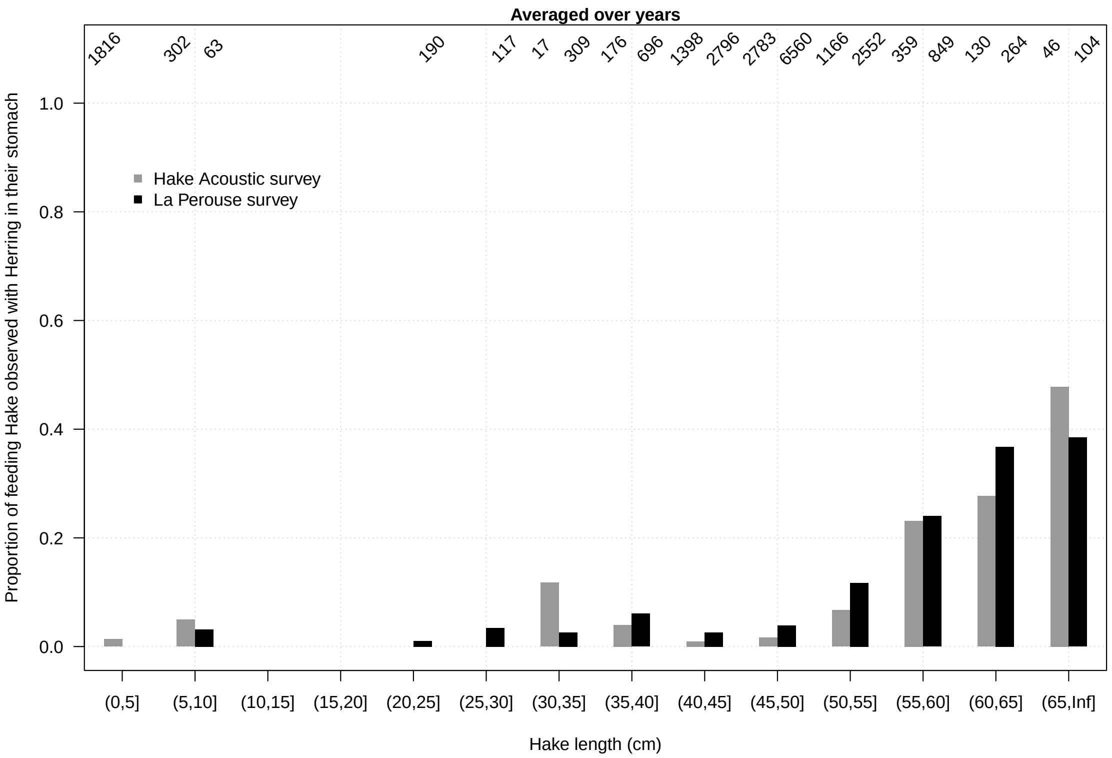


Figure B.6: Proportion of feeding Hake observed with herring in their stomach contents (y-axis), broken up into 5cm Hake length bins (x-axis) with all Hake above 65 cm combined. Data are split into Hake Acoustic and La Perouse surveys, and aggregated over years.

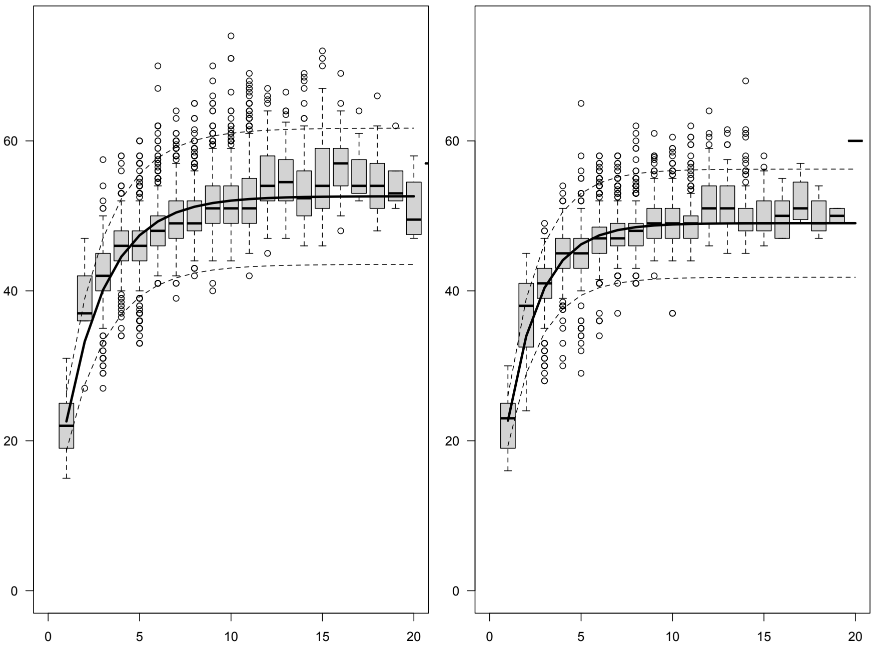


Figure B.7: Pacific Hake size-at-age data (box plots) and von Bertalanffy growth models (lines) for female (left) and male (right) Hake.

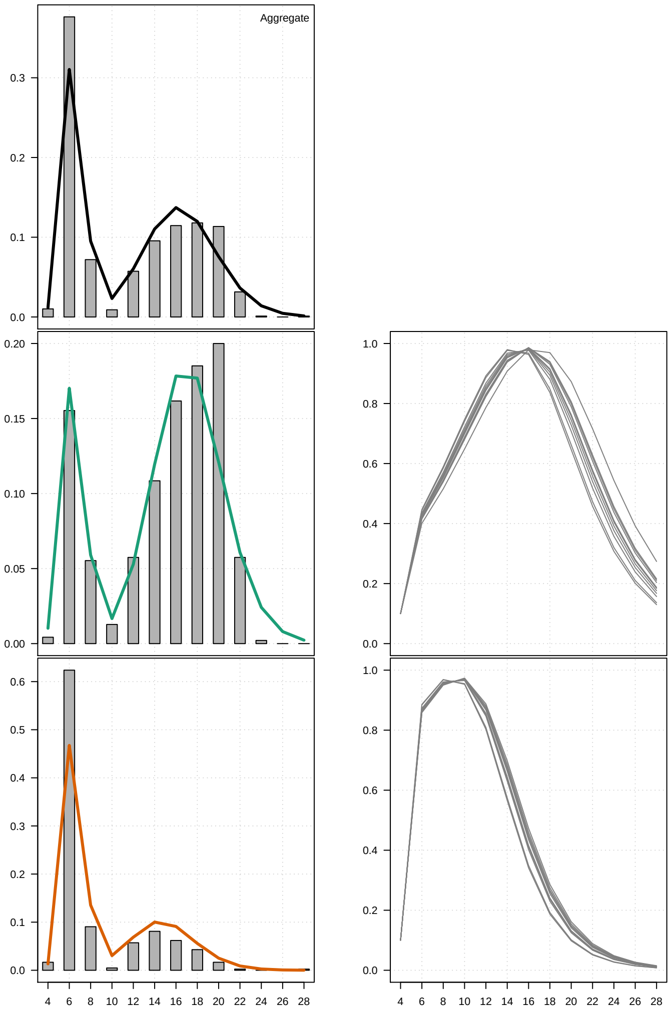


Figure B.8: Left column: Pacific Hake length compositions from stomach contents observational data collected off the coast of WCVI (rectangles) aggregated across all Hake (top panel) and split by Hake size class (above and below 50 cm fork length), with predicted length compositions from the fitted dome-shaped size-selectivity models (lines). Right column: time-varying dome-shaped size-selectivity of Herring by Pacific Hake on La Perouse Bank.

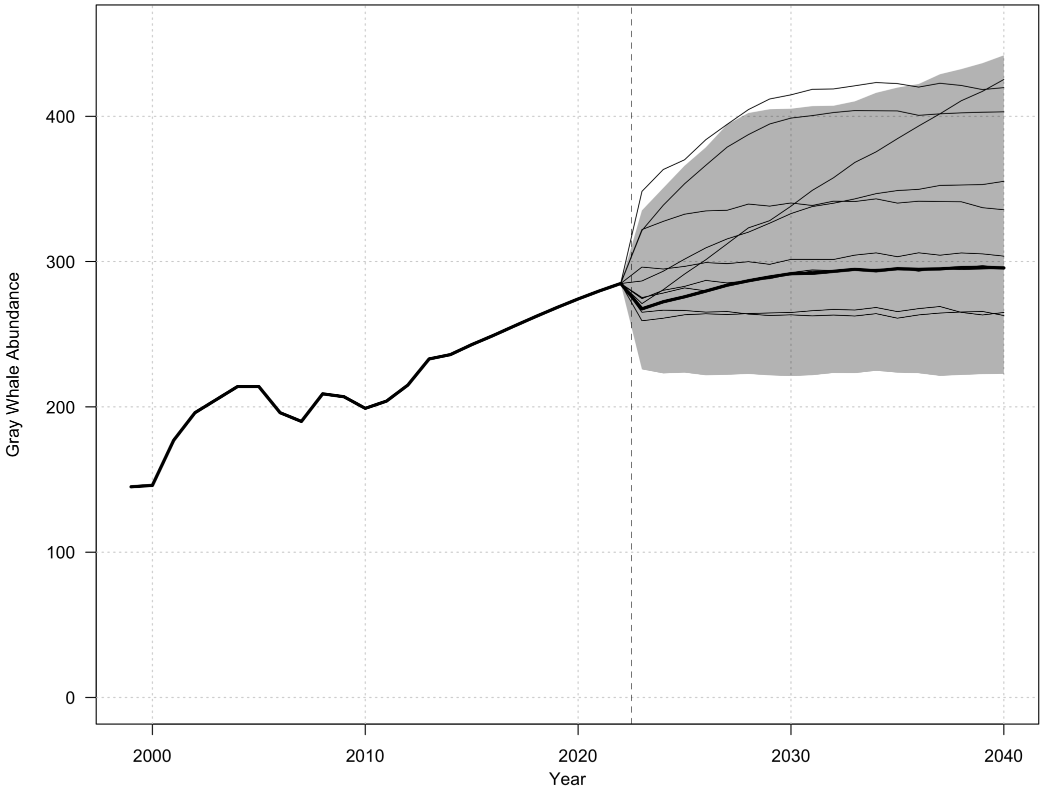


Figure B.9: Capture-mark-recapture estimates of PCFG Grey Whale abundance and simulated projections of PCFG abundance, reproduced from data and $\theta$-logistic model parameters supplied by Gavrilchuk and Doniol-Valcroze (2021).

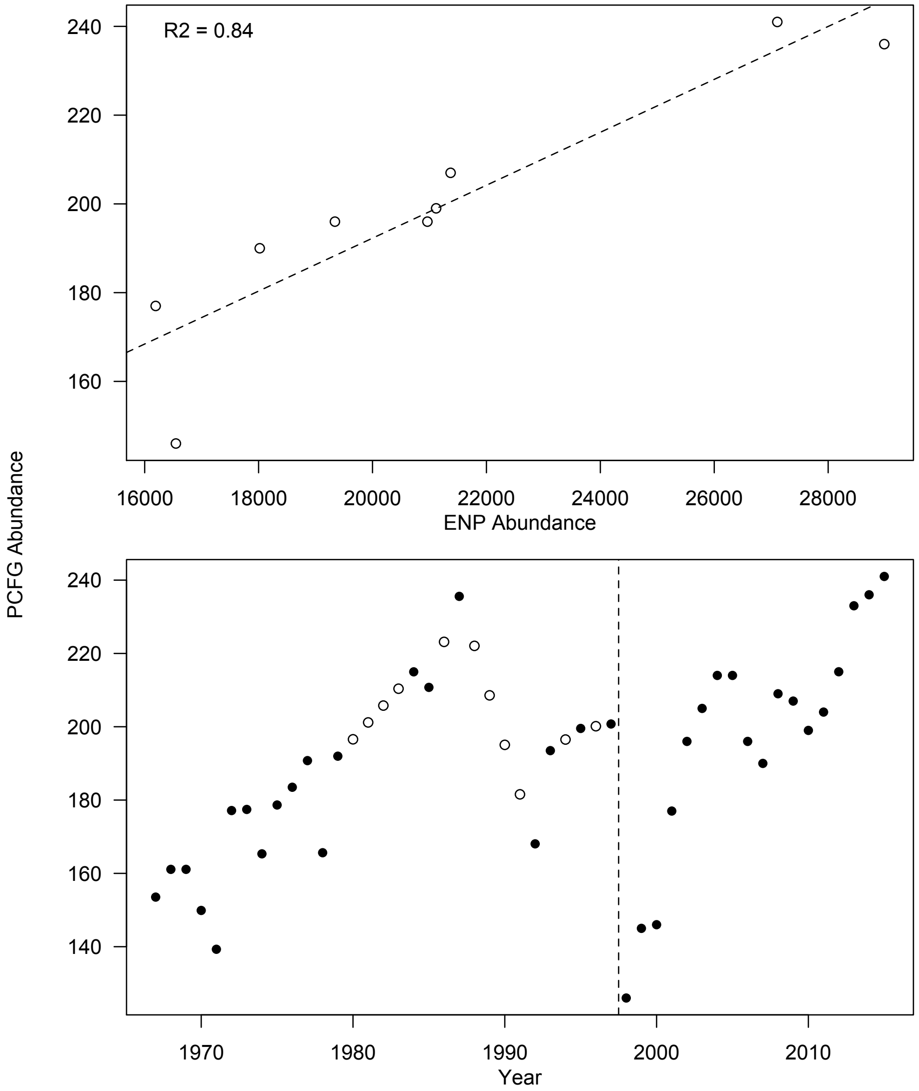


Figure B.10: The two steps of estimating PCFG abundance from ENP abundance in years before 1998, showing the linear model (top) estimating the proportion PCFG whales within the ENP population, and (bottom) the interpolation of PCFG abundance from ENP abundance estimates prior to 1998 (vertical dashed line).

John Calambokidis. Abundance estimates of humpback and blue whales off the us west coast based on mark-recapture of photo-identified individuals through 2008. *Cascadia Research, Olympia, Washington*, 2009.

John Calambokidis, Gretchen H Steiger, David K Ellifrit, Barry L Troutman, and C Edward Bowlby. Distribution and abundance of humpback whales (*Megaptera novaeangliae*) and other marine mammals off the northern washington coast. 2004.

John Calambokidis, Erin A Falcone, Terrance J Quinn, Alexander M Burdin, Phillip J Clapham, John KB Ford, Christine M Gabriele, Richard LeDuc, David Mattila, Lorenzo Rojas-Bracho, et al. Splash: Structure of populations, levels of abundance and status of humpback whales in the north pacific. *Final report for Contract AB133F-03-RP-00078*, 57, 2008.

COSEWIC. COSEWIC assessment and update status report on the grey whale (Eastern North Pacific population) *Eschrichtius robustus* in Canada. pages vii + 31 p. Committee on the Status of Endangered Wildlife in Canada, Ottawa, 2004.

James D Darling. Gray whales off vancouver island, british columbia. *The gray whale, Eschrichtius robustus*, pages 267–287, 1984.

James D Darling, Kathleen E Keogh, and Tammy E Steeves. Gray whale (*Eschrichtius robustus*) habitat utilization and prey species off vancouver island, bc. *Marine Mammal Science*, 14(4):692–720, 1998.

DFO. Population assessment pacific harbour seal (*Phoca vitulina richardsi*). *DFO Can. Sci. Advis. Sec. Sci. Advis. Rep.*, 2009/011, 2010.

JW Durban, DW Weller, AR Lang, and WL Perryman. Estimating gray whale abundance from shore-based counts using a multilevel bayesian model. *J. Cetacean Res. Manag*, 15:61–68, 2015.

JW Durban, DW Weller, and WL Perryman. Gray whale abundance estimates from shore-based counts off california in 2014/15 and 2015/16. Technical report, Paper SC, 2017.

JKB Ford, AL Rambeau, RM Abernethy, MD Boogaards, LM Nichol, and LD Spaven. An assessment of the potential for recovery of humpback whales off the Pacific coast of Canada. *DFO Canadian Science Advisory Secretariat Research Document*, 15:33, 2009.

John KB Ford, Kathy Heise, Lance G Barrett-Lennard, and Graeme M Ellis. Killer whales and other cetaceans of the queen charlotte islands/haida gwaii. *Report to the Gwaii Haanas National Park Reserve, Parks Canada*, 1994.

John KB Ford, John W Durban, Graeme M Ellis, Jared R Towers, James F Pilkington, Lance G Barrett-Lennard, and Russel D Andrews. New insights into the northward migration route of gray whales between vancouver island, british columbia, and southeastern alaska. *Marine Mammal Science*, 29(2):325–337, 2013.

RIC Chris Francis. Replacing the multinomial in stock assessment models: A first step. *Fisheries Research*, 151:70–84, 2014.

Katherine Gavrilchuk and Thomas Doniol-Valcroze. Recovery Potential Assessment for the Grey Whale (*Eschrichtius robustus*): Pacific Coast Feeding Group and Western Pacific Population in Canadian Waters. Technical Report 20, Fisheries and Oceans, Canada, 2021.

YULIA V Ivashchenko, ALEX N Zerbini, and PHILLIP J Clapham. Assessing the status and preexploitation abundance of north pacific humpback whales. *International Whaling Commission Paper SC/66a/IA/16*, 2016.

Kelli F Johnson, AM Edwards, Aaron M Berger, and CJ Grandin. Status of the pacifc hake (whiting) stock in u.s. and canadian waters in 2021. Technical report, Prepared by the Joint Technical Committee of the U.S. and Canada Pacifc Hake/Whiting Agreement, National Marine Fisheries Service and Fisheries and Oceans Canada, 2021.

Samuel D N Johnson, Sean P Cox, Jaclyn S Cleary, AJ Benson, SJH Power, and Steven P Rossi. Application of a new modelling framework for the assessment of Pacific Herring (*Clupea pallasii*) major stocks and implementation in the management strategy evaluation process. *DFO Can. Sci. Advis. Sec. Res. Doc.*, 2024/nnn(iv + 233 p.), 2024.

Kasper Kristensen, Anders Nielsen, Casper W Berg, Hans Skaug, and Brad Bell. Tmb: Automatic differentiation and laplace approximation. *arXiv preprint arXiv:1509.00660*, 2015.

Mark N Maunder, Paul J Starr, and Ray Hilborn. A bayesian analysis to estimate loss in squid catch due to the implementation of a sea lion population management plan. *Marine Mammal Science*, 16(2):413–426, 2000.

Christie McMillan, Elise A Keppel, Lisa D Spaven, and Thomas Doniol-Valcroze. Preliminary report on the seasonal abundance and distribution of cetaceans in the southern salish sea in response to tmx recommendations 5 and 6 (year 1). Technical report, Can. Tech. Rep. Fish. Aquat. Sci. 3474: vi + 33 p., 2022.

SA Mizroch, LM Herman, JM Straley, DA Glockner-Ferrari, C Jurasz, J Darling, S Cerchio, CM Gabriele, DR Salden, and O Von Ziegesar. Estimating the adult survival rate of central north pacific humpback whales (*Megaptera novaeangliae*). *Journal of Mammalogy*, 85(5):963–972, 2004.

JM Mobley, Scott Spitz, Richard Grotefendt, Paul Forestell, Adam Frankel, and Gordon Bauer. Abundance of humpback whales in hawaiian waters: Results of 1993-2000 aerial surveys. *Report to the Hawaiian Islands Humpback Whale National Marine Sanctuary*, 9, 2001.

JR Mobley, Gordon B Bauer, and Louis M Herman. Changes over a ten-year interval in the distribution and relative abundance of humpback whales (*Megaptera novaeangliae*) wintering in hawaiian waters. *Aquatic Mammals*, 25:63–72, 1999.

Cole C Monnahan and Kasper Kristensen. No-u-turn sampling for fast bayesian inference in admb and tmb: Introducing the adnuts and tmbstan r packages. *PloS one*, 13(5):e0197954, 2018.

Cole C Monnahan, James T Thorson, and Trevor A Branch. Faster estimation of bayesian models in ecology using hamiltonian monte carlo. *Methods in Ecology and Evolution*, 8(3):339–348, 2017.

L Nichol and K Heise. The historical occurrence of large whales off the queen charlotte islands. *Unpublished report prepared for the South Moresby/Gwaii Haanas National Park Reserve, Canada Parks Service, Queen Charlotte City, BC*, 1992.

Linda Marie Nichol, R Abernethy, L Flostrand, T. S. Lee, and J. K. B. Ford. Information relevant to the identification of critical habitats of North Pacific humpback whales (*Megaptera novaeangliae*) in British Columbia. *DFO Can. Sci. Advis. Sec. Res. Doc*, (2009/116), 2010.

Peter F Olesiuk. An assessment of the status of harbour seals (*Phoca vitulina*) in british columbia. *Canadian Stock Assessment Secretariat Research Document 99/33*, 1999.

Peter F Olesiuk. An assessment of population trends and abundance of harbour seals (*Phoca Vitulia*) in British Columbia. *DFO Canadian Science Advisory Secretariat Research Document*, (105), 2009.

Peter F Olesiuk. Recent trends in abundance of steller sea lions (*Eumetopias jubatus*) in British Columbia. *DFO Canadian Science Advisory Secretariat Research Document*, (6):67, 2018.

R Core Team. *R: A Language and Environment for Statistical Computing*. R Foundation for Statistical Computing, Vienna, Austria, 2015.

Timothy J Ragen. Maximum net productivity level estimation for the northern fur seal (*Callorhinus ursinus*) population of St. Paul Island, Alaska. *Marine Mammal Science*, 11(3):275–300, 1995.

Andrew J Read and Paul R Wade. Status of marine mammals in the united states. *Conservation Biology*, 14 (4):929–940, 2000.

Randall R Reeves, Stephen Leatherwood, Stephen A Karl, and Evelyn R Yohe. Whaling results at akutan (1912-39) and port hobron (1926-37), alaska. *Report of the International Whaling Commission*, 35: 441–457, 1985.

Jon T Schnute and Rowan Haigh. Compositional analysis of catch curve data, with an application to sebastes maliger. *ICES Journal of Marine Science: Journal du Conseil*, 64(2):218–233, 2007.

JF Schweigert, Jaclyn S Cleary, and P. Midgley. Synopsis of the Pacific Herring Spawn-on-Kelp Fishery in British Columbia. *Canadian Manuscript Reports of Fisheries and Aquatic Sciences*, (3148):vi + 33, 2018.

Thomas Shields and Gary Kingston. Herring impoundment and spawn-on-kelp production in british columbia. *Victoria: Archipelago Marine Research*, 1982.

LL Wedekin, MH Engel, A Andriolo, PI Prado, AN Zerbini, MMC Marcondes, PG Kinas, and PC Simões-Lopes. Running fast in the slow lane: rapid population growth of humpback whales after exploitation. *Marine Ecology Progress Series*, 575:195–206, 2017.

Alexandre N Zerbini, Janice M Waite, Jeffrey L Laake, and Paul R Wade. Abundance, trends and distribution of baleen whales off western alaska and the central aleutian islands. *Deep Sea Research Part I: Oceanographic Research Papers*, 53(11):1772–1790, 2006.

Alexandre N Zerbini, Phillip J Clapham, and Paul R Wade. Assessing plausible rates of population growth in humpback whales from life-history data. *Marine Biology*, 157:1225–1236, 2010.
